# Dopamine D1 receptor activation shapes anterior insula neural coding of anxiety

**DOI:** 10.1101/2024.10.25.620186

**Authors:** Yoni Couderc, Tanmai Dhani Reddy, Archi Garg, Daria Ricci, Giovanni Vardiero, Marion d’Almeida, Céline Nicolas, Joeri Bordes, Tina Habchi, Camille Penet, Yue Kris Wu, Julijana Gjorgjieva, Yulong Li, Emmanuel Valjent, Anna Beyeler

## Abstract

The anterior insula^1–3^ and dopamine neuromodulation^4–11^ both play key roles in the control of anxiety, yet how dopamine shapes anterior insula function to regulate anxiety remains unknown. Here we show that dopaminergic neurons of the ventral tegmental area preferentially target the anterior relative to the posterior insula, and that optogenetic activation of these neurons elicits dopamine signals in the anterior insula. Behaviorally, dopamine signals increased in the anterior insula during risk assessment and exploration of exposed spaces. Interestingly, neurons expressing the type-1 dopamine receptor (D1) are enriched in the anterior insula subdivision, where their optogenetic activation is anxiogenic. At the molecular level, direct D1 activation or blockade in the anterior insula bidirectionally controls anxiety, demonstrating a causal anxiogenic function of D1 in the anterior insula. Remarkably, systemic D1 activation also increased anxiety-related behaviors, together with a cellular activation of the anterior insula, and a disruption of neural coding in this region. As an example of the latter, systemic D1 activation oppositely regulated the coding reliability of exposed and protected areas, increasing the reliability of the neural code for exposed spaces. Together, our findings reveal an anterior insula D1-dependent mechanism by which dopamine can control anxiety, providing a framework for investigating dopamine dysregulation in models of anxiety disorders. Our study also introduces quantitative metrics applied to AI-based computational representations of neural activity to identify how neuromodulation reshapes neural coding of behaviors and contexts.

## Main

Anxiety is an adaptive response to potential threats, yet persistent and high levels of anxiety can become pathological^12,13^. Anxiety disorders are the most prevalent psychiatric conditions, characterized by chronic and excessive worry, along with other symptoms. Heightened insular cortex (or insula) activity has been reported in patients with anxiety disorders. Indeed, neuroimaging studies consistently report insular overactivation in response to salient and negative stimuli in patients with anxiety disorders, supporting a contribution of insular circuits in the pathophysiology of anxiety^14–19^. Moreover, pharmacological and optogenetic manipulations in rodents have further demonstrated that activation of the anterior insula is anxiogenic^1,2,20^. While dopamine is best known for its role in reward and motivation^21,22^, it also modulates anxiety levels through actions on D1 and D2 receptors in humans^11^ and preclinical models^4–10^. In cortical and subcortical regions, the predominance of D1 over D2 receptors^23^ suggests that D1 signaling may be a primary driver of dopaminergic modulation. Interestingly, pharmacological activation of D1 in the basolateral amygdala (BLA) is anxiogenic^4^. However, activation of D1 receptors in the medial prefrontal cortex (mPFC) is anxiolytic^24^, underscoring the complex and region-specific nature of dopaminergic control of anxiety. Yet, the topography of D1+ neurons in the insula, and whether D1 receptor signaling in the anterior insula causally controls anxiety-related behaviors, remain unknown. Here, combining anatomical mapping, pharmacology, optogenetics, *in vivo* electrophysiology, as well as advanced computational methods, we identify a D1-dependent mechanism by which dopamine shapes anterior insula neural dynamics for protective and risk-assessment behaviors.

### VTA dopamine neurons preferentially innervate the anterior over posterior insula

To dissect the dopaminergic innervation of the insular cortex, we performed both retrograde and anterograde mapping of dopaminergic projections to the insula. First, we injected Cholera Toxin B (CTB) retrograde tracers into the anterior and posterior insular cortex (aIC and pIC, respectively) combined with tyrosine hydroxylase (TH) immunostaining to identify dopaminergic neurons. We then quantified insula-projecting dopaminergic neurons (co-labeled with TH and CTB) in the ventral tegmental area (VTA) as well as substantia nigra pars compacta (SNc, **Fig. 1a-b and Extended Data Fig. 1a**). We found comparable numbers of VTA dopaminergic neurons (VTA^DA^) and SNc dopaminergic neurons (SNc^DA^) projecting to the insula overall (**Extended Data Fig. 1b**). However, we found more VTA^DA^ and SNc^DA^ neurons projecting to the anterior insula compared to posterior insula (**Fig. 1c-d**). Next, we tested whether dopaminergic innervation of the insula shows laminar specificity depending on its VTA or SNc origin. To this end, we expressed eYFP or mCherry in a cre-dependent manner in the VTA or SNc of DAT-cre mice (**Fig. 1e-f and Extended Data Fig. 1c**). Interestingly, we revealed a denser axonal arborization from VTA^DA^ neurons in the anterior than the posterior insula. SNc^DA^ terminal density was overall similar across cortical layers in both anterior and posterior insula, except for the layer 6 with a higher density in the posterior insula, compared to anterior insula (**Fig. 1g-h and Extended Data Fig. 1c-d**).

**Fig. 1:**
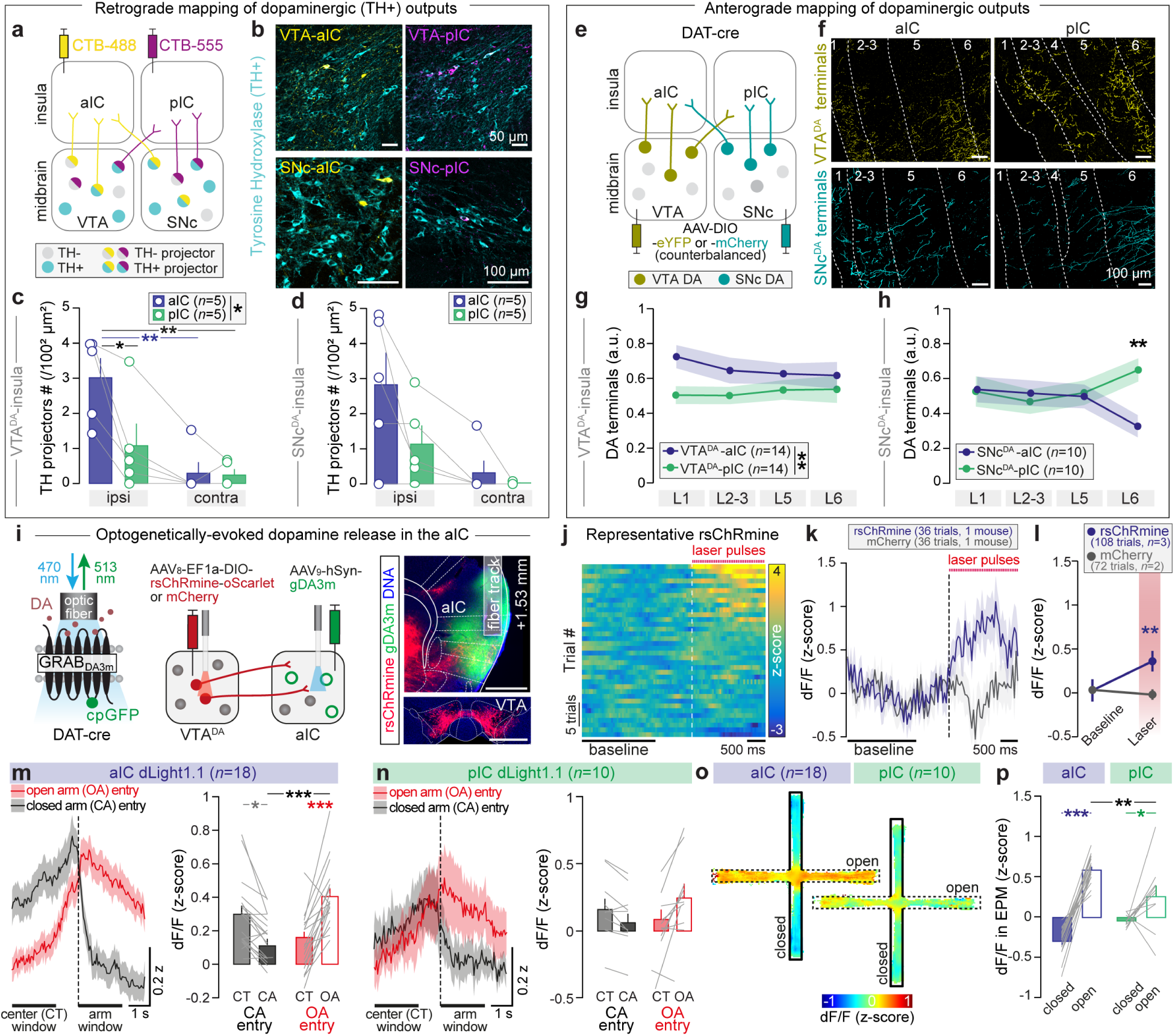
Dopaminergic system architecture of the insular cortex. **a,** Retrograde mapping of dopaminergic projections to the anterior (aIC) and posterior (pIC) insula, combined with Tyrosine Hydroxylase (TH) immunostaining. **b,** Confocal images of VTA^DA^ (left) and SNc^DA^ (right) neurons projecting to the insula. **c,** Higher number of VTA^DA^ neurons projecting to the aIC (\**p*=0.0297) relative to pIC (Main effect Region: F_1,16_=4.91, \**p=*0.0415, Main effect Side: F_1,16_=15.7, \*\**p=*0.0011, with a Region x Side interaction: F_1,16_=4.44, \**p=*0.05). **d,** No difference in the number of SNc^DA^ neurons projecting to either aIC or pIC (Region: F_1,16_=3.34, *p=*0.0863, Main effect Side: F_1,16_=11, \*\**p=*0.0044, and no Region x Side interaction: F_1,16_=1.57, *p=*0.2288). **e,** Anterograde mapping of dopaminergic projections to aIC and pIC. **f,** Confocal images of VTA^DA^ and VTA^DA^ axonal terminals in aIC and pIC. **g,** No layer specificity observed from VTA^DA^ terminals in both aIC and pIC (Main effect Region: F_1,104_=8.87, \*\**p=*0.0036, Layer: F_3,104_=0.175, *p=*0.9131, with no Region x Layer interaction: F_3,104_=0.489, *p=*0.6909). **h,** SNc^DA^ terminals are denser in pIC layer 6 (\*\**p*=0.0012) compared to aIC (Region: F_1,72_=2.17, *p=*0.1450, Layer: F_3,72_=0.127, *p=*0.9440, with a Region x Layer interaction: F_3,72_=3.19, \**p=*0.0288). **i,** Left: Viral approach to evoke dopamine signals in the aIC (hSyn-gDA3m) by optogenetic activation of VTA^DA^ neurons expressing rsChRmine-oScarlet (or the control vector expressing mCherry) in DAT-cre mice. Right: Representative fluorescence image of injection sites. **j,** Representative heatmap of an rsChRmine-expressing mouse showing z-scored dopamine signal in the aIC, aligned to the onset of light stimulation (20 Hz, 10 mW, 2s, 10 ms pulses). Trials were sorted based on dopamine signal activity during the laser stimulation period (0-2s). **k**, Averaged z-scored dopamine signal increased during laser stimulation for one rsChRmine-expressing mouse compared to baseline, while no change is observed for the mCherry-expressing mouse. **l,** VTA^DA^ cell bodies optogenetic stimulation of rsChRmine-expressing mice increases dopamine signals in the aIC (\*\**p*=0.0062) compared to controls (Group: F_1,356_=2.33, *p*=0.1278, Main effect Period: F_1,356_=4.22, \**p*=0.0407, with a Group x Period interaction: F_1,356_=4.41, \**p*=0.0365). Quantification panels for **j-l** panels: baseline window [-3s to -1s] and laser windows [0s to 2s]. **m,** DA signal in the aIC decreases upon closed arm (CA) entries (\**p*=0.0108) and increases upon open arm (OA) entries (\*\*\**p*=0.0005), with a stronger increase (\*\*\**p*<0.0001) at OA entry compared to CA entry in aIC (Space: F_1,68_=3.55, *p*=0.0638, Event: F_1,68_=0.4605, *p*=0.4997, with a Space x Event interaction: F_1,68_=27.15, \*\*\**p*<0.0001). **n,** Although dopamine signal in the pIC at entries into open and closed follows similar direction than in the aIC, no statistical difference was found between entries into both arms (Space: F_1,36_=0.5154, *p*=0.4774, Event: F_1,36_=0.1555, *p*=0.6957, with no Space x Event interaction: F_1,36_=1.783, *p*=0.1902). Quantification windows for **m-n** panels: center (CT) window: [-3s to -1s] and arm entry window [0s to 2s]. **o-p,** Averaged heatmaps (**o**) and quantification (**p**) of z-scored DA signal in aIC and pIC during EPM exploration showed increased DA signal in open vs. closed arms in aIC (\*\*\**p<*0.0001) and pIC (\**p=*0.0160), with a stronger signal in aIC (\*\**p=*0.0028) relative to pIC (**p,** Main effect Arm: F_1,52_=92.6, \*\*\**p<*0.0001, Region: F_1,52_=0.35, *p=*0.5565, with an Arm x Region interaction: F_1,52_=21.6, \*\**p=*0.0001). **a-d**: aIC and pIC (n=5 mice). **e-h:** VTA (n=14 mice) and SNc (n=10 mice). **i-l:** rsChRmine (n=3 mice) and mCherry (n=2 mice). **m-p**: aIC (n=18 mice) and pIC (n=10 mice). **c-d, g-h, l-n, p**: Two-way ANOVA followed by Tukey’s/Holm-Sidak’s post hoc tests. Data are: mean ± SEM.

Given that VTA^DA^ neurons preferentially innervate the anterior over the posterior insula, we tested whether optogenetic activation of VTA^DA^ neurons is sufficient to elicit dopamine signal in the anterior insula. DAT-cre mice were injected with a viral vector expressing the red-shifted opsin rsChRmine fused to oScarlet (or a control vector expressing mCherry) into the VTA, together with a viral vector expressing the genetically-encoded dopamine sensor gDA3m^25,26^ into the anterior insula (**Fig. 1i**). Optogenetic activation of VTA^DA^ cell bodies increased dopamine signals in the anterior insula compared to controls (**Fig. 1j-l** and **Extended Data Fig. 1e**), demonstrating that VTA dopaminergic inputs can drive dopamine release in the anterior insula.

### Dopamine signals are higher in the anterior insula at entry into and during exploration of exposed areas

To determine how anatomical differences of dopaminergic inputs to the insula translate into dopamine dynamics during anxiety-related behaviors, we expressed the genetically-encoded dopamine sensor dLight1.1^27^ under the pan-neuronal hSyn promoter in either the anterior or posterior insula (**Extended Data Fig. 1f**). We then recorded dopamine dynamics in the anterior or posterior insula during the elevated-plus maze (EPM, **Fig 1m-p**, **Extended Data Fig. 1g-h**) and the open field test (OFT, **Extended Data Fig. 1i-j**). Dopamine signals were time-locked to entries into open (exposed) and closed (safe) arms in the EPM. In the anterior insula, dopamine signals sharply decreased at entry into closed arms, and increased at entry into open arms (**Fig. 1m**). A similar bidirectional response to safe versus exposed environments was observed in the posterior insula, although with smaller amplitude (**Fig. 1n**). Moreover, dopamine signals were higher in the anterior compared to posterior insula, when mice were located in the open arms of the EPM(**Fig. 1o-p**), and similarly, when mice were in the center of the OFT **(Extended Data Fig. 1i-j**). However, the magnitude of open arm evoked dopamine signal did not correlate with the level of anxiety in mice (estimated by the time mice spent in EPM open arms or OFT center, **Extended Data Fig. 1k-l**). Importantly, these dopamine changes were not correlated with velocity in open arms or OFT center (**Extended Data Fig. 1m-n**). Interestingly, dopamine transient frequency was higher in the posterior compared to anterior insula in the EPM, but independently of whether mice were in protected or exposed areas (**Extended Data Fig. 1o**). Together, these findings reveal stronger dopamine signals in the anterior relative to posterior insula during exploration of exposed environments, consistent with the preferential VTA dopaminergic innervation of the anterior compared to posterior insula.

### Dopamine signaling onto D1+ neurons in the anterior insula scales with anxiety levels

To parse the role of dopamine signaling in the insula, we examined the topography of dopaminoceptive neurons across anterior and posterior subdivisions and cortical layers. We used D1-cre and D2-cre mouse lines crossed with the Ai14 reporter line to generate double transgenic mice expressing tdTomato selectively in D1+ or D2+ neurons, respectively (**Fig. 2a and Extended Data Fig. 2a**). We observed a higher density of D1+ neurons in the anterior compared to the posterior insula. Moreover, D1+ neuron density was more than sevenfold greater than D2+ neuron density in both anterior and posterior insula (**Fig. 2b**). Across cortical layers, D1+ neurons were particularly enriched in layer 2/3 of the anterior relative to the posterior insula. Conversely, D2+ neurons were equally abundant in the anterior and posterior insula (**Fig. 2c**). Although the absolute number of D1+ but not D2+ neurons was higher among excitatory than inhibitory neurons (**Extended Data Fig. 2b**), the proportion of D1+ and D2+ neurons within inhibitory populations was higher in both anterior and posterior insula (**Extended Data Fig. 2c**). To assess potential overlap between dopamine receptor subtypes, we performed fluorescent *in situ* hybridization and detected limited co-expression of *Drd1* and *Drd2* transcripts, with 10% of cells in the anterior insula and 6% in the posterior insula co-expressing both receptors (**Extended Data Fig. 2d-f**).

**Fig. 2:**
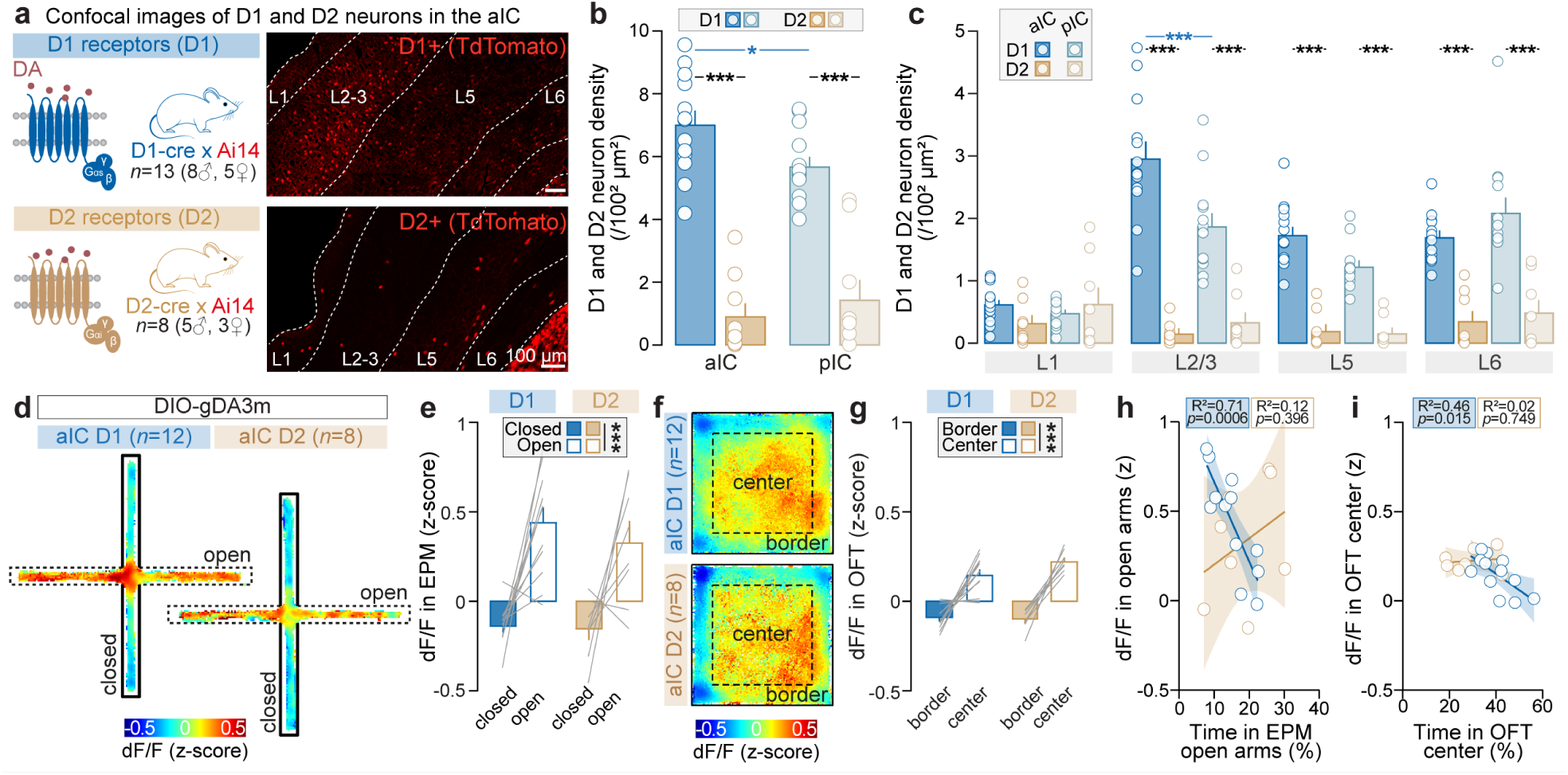
Dopamine signal in the anterior insula encodes anxiety-related states. **a,** Confocal images of tdTomato expression in D1+ and D2+ neurons across aIC layers. **b,** Density of dopaminoceptive neurons reveals a predominance of D1+ over D2+ neurons in aIC (\*\*\**p*<0.0001) and pIC (\*\*\**p*<0.0001), and a higher density of D1+ neurons in the aIC (\**p*=0.0480) relative to pIC (Main effect Receptor: F_1,40_ = 134, \*\*\**p*< 0.0001, Region: F_1,40_=0.821, *p*=0.3703, with a Receptor x Region interaction: F_1,40_=4.36, \**p=*0.0432). **c,** Laminar distribution of dopaminoceptive neurons shows an enrichment of D1+ neurons in the aIC layer 2/3 (\*\*\**p*<0.0001) compared to pIC (Main effect Receptor: F_1,152_=203.3, \*\*\**p*<0.0001, Region: F_1,152_=1.113, *p=*0.2930, Main effect Layer: F_3,152_=16.88, \*\*\**p*<0.0001, with a Receptor x Region x Layer interaction : F_3,152_=3.138, \**p=*0.0272). **d-e,** Averaged heatmaps (**d**) and quantification (**e**) of z-scored DA signal onto D1+ and D2+ aIC neurons in the EPM shows increased DA signal targeting both D1+ and D2+ neurons in open (vs. closed) arms of the EPM (Main effect Arm: F_1,36_=48.9, \*\*\**p<*0.0001, Receptor: F_1,36_=0.695, *p=*0.41, with no Arm x Receptor interaction: F_1,36_=0.414, *p=*0.5243). **f-g,** Averaged heatmaps (**f**) and quantification (**g**) of z-scored DA signal onto D1+ and D2+ aIC neurons in the OFT shows increased DA signals onto D1+ and D2+ aIC neurons in center (vs. borders) of the OFT (Main effect Zone: F_1,36_=131, \*\*\**p<*0.0001, Receptor: F_1,36_=1.84, *p=*0.1833, with no Zone x Receptor interaction: F_1,36_=2.90, *p=*0.0973). **h,** DA signal onto D1+ aIC neurons when mice are in open arms negatively correlates with the time spent in open arms (two-tailed Pearson’s correlation, D1: R²=0.71, \*\*\**p=*0.0006; D2: R²=0.1222, *p=*0.3960). **i,** DA signal onto D1+ aIC neurons when mice are in OFT center negatively correlates with the time spent in OFT center (two-tailed Pearson’s correlation, D1: R²=0.4596, \**p=*0.0156; D2: R²=0.0183, *p=*0.7493). **a-c:** D1 (n=13 mice) and D2 (n=8 mice). **d-i:** aIC D1 (n=12 mice) et aIC D2 (n=8 mice). **b-c, e, g**: Two-way ANOVA (three-way in **c**) and Tukey’s (or Holm-Sidak’s) post hoc tests. Data are: mean ± SEM.

Given the divergent densities and distribution of D1+ and D2+ neurons in the insula, we next examined dopamine dynamics specifically in these neuronal subtypes during anxiety-related behaviors by selectively expressing the dopamine sensor gDA3m^25,26^ in D1-cre or D2-cre mice. Because dopamine signals were higher in the anterior compared to posterior insula when mice explored exposed spaces, we performed recordings in the anterior insula (**Extended Data Fig. 2g**). Time-locked analyses during entry into EPM open arms revealed neuron subtype-specific effects. Indeed, dopamine preferentially targeted D1+ neurons at open arm entry, whereas this effect was not detected in D2+ neurons. Conversely, the decrease in dopamine signal at closed arm entry was significant only in D1+ neurons (**Extended Data Fig. 2h-i**).

When mice were in open arms of the EPM (**Fig. 2d-e**) or in the center of the OFT (**Fig. 2f-g**), dopamine signals onto both D1+ and D2+ anterior insula neurons increased. However, only the amplitude of dopamine signal measured in D1+ neurons during the exposure to open arms or OFT center was negatively correlated with the time spent in these exposed spaces (**Fig. 2h-i**). This indicates a selective link between the anxiety levels and the amount of dopamine released targeting D1 anterior insula neurons. We did not detect changes in dopamine transient frequency or amplitude in the anterior insula during the EPM (**Extended Data Fig. 2j-k**). Importantly, dopamine signals were not correlated with the velocity in the open arms or OFT center (**Extended Data Fig. 2l-m**), suggesting that these dopamine fluctuations reflect anxiety state rather than locomotor activity.

Previous studies have shown that glutamatergic neurons of the insula project to downstream regions involved in anxiety^2,28,29^, including the basolateral amygdala (BLA), the bed nucleus stria terminalis and central amygdala nuclei. We therefore examined whether D1+ neurons in the anterior and posterior insula share similar downstream targets. Long-range synaptic tracing of D1 anterior and posterior insula neurons revealed strong projections to ipsilateral amygdala nuclei and to contralateral insula (**Extended Data Fig. 2n-q**), mirroring the projection patterns of glutamatergic insula neurons.

Overall, these findings demonstrate that the insular cortex, particularly its anterior subdivision, is enriched in D1+ neurons and that dopamine signaling onto D1+ anterior insula neurons scales with anxiety levels during exploration of exposed spaces, supporting the hypothesis of a D1-dependent regulation of anxiety in the anterior insula.

### D1 receptor activation in the anterior insula bidirectionally controls anxiety

Based on the positive correlation between dopamine signaling onto D1+ anterior insula neurons and anxiety levels, and prior evidence supporting an anxiogenic role of the anterior insula^1,2^, we tested whether D1 receptor activation promotes anxiety at both systemic and intra-anterior insula levels. First, we administered the selective D1 agonist SKF38393 or vehicle, intraperitoneally 30 minutes before behavioral testing (10 mg/kg, **Fig. 3a**). Systemic D1 activation reduced the time spent in the open arms of the EPM (**Fig. 3b-c**) and in the center of the OFT (**Fig. 3e-f**), without affecting total distance travelled in either test (**Fig. 3d,g**). To evaluate the impact of systemic D1 activation on neuronal activity, we performed cFOS immunostaining 90 minutes after intraperitoneal injection of SKF38393 or vehicle (**Fig. 3h**). Compared to vehicle, D1 activation increased the number of cFOS+ cells in both anterior and posterior insula, as well as and in the nucleus accumbens (NAc), but not the BLA or S1 somatosensory cortex (**Fig. 3j**), Remarkably, the increase of cFOS+ cells was the highest in the anterior insula compared to the posterior insula and NAc. Collectively, our results indicate that systemic D1 activation increases anxiety-related behaviors, as well as an activation of anterior insula activity.

**Fig. 3:**
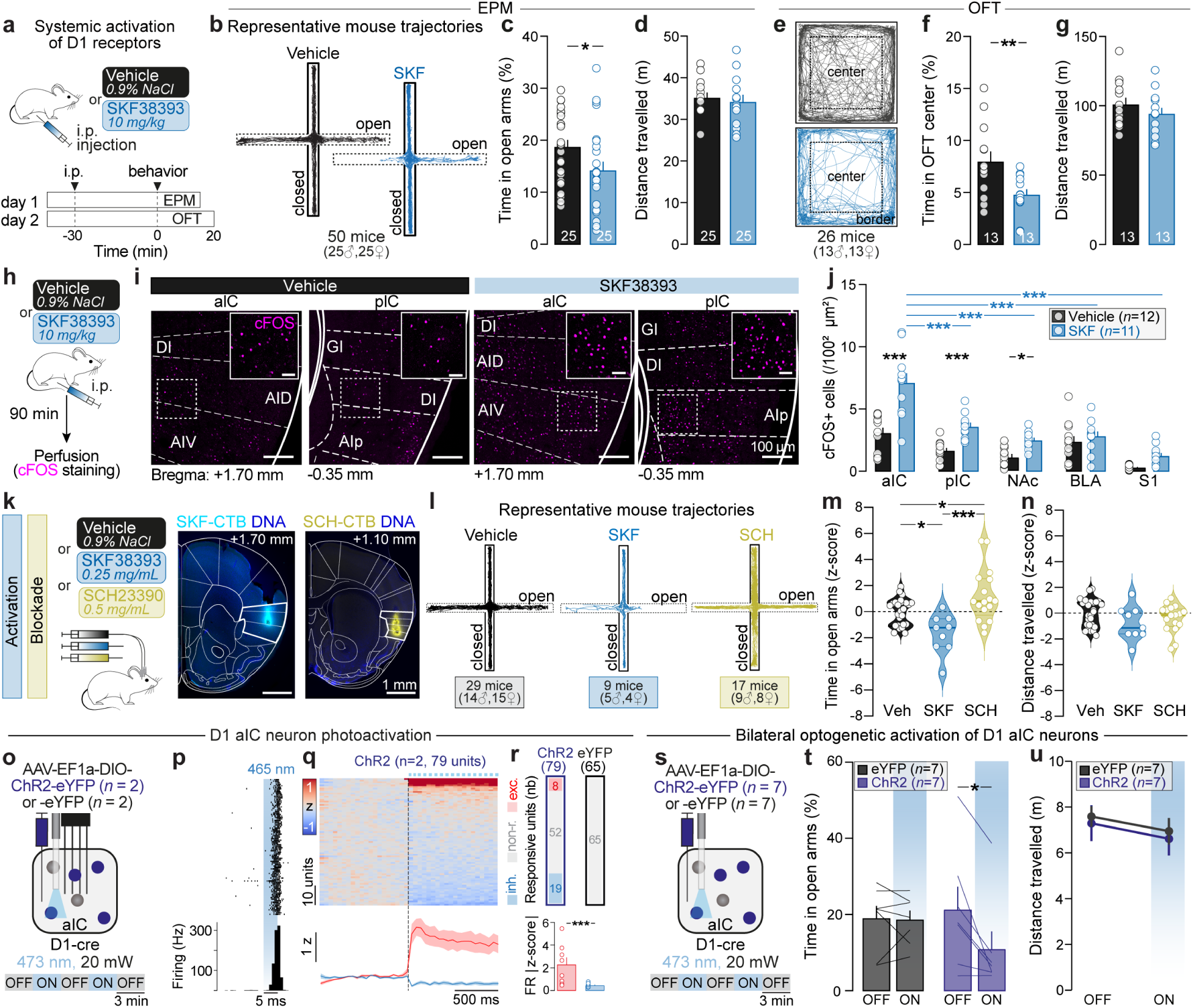
D1 receptors of the anterior insula drive anxiety-related behaviors. **a,** Systemic activation of D1 receptors. Mice received an intraperitoneal (i.p.) injection of vehicle or D1 agonist SKF38393 (SKF, 10 mg/kg) 30 minutes before EPM testing. **b-e,** Representative mice trajectory treated with vehicle or SKF38393 in the EPM (**b**) and OFT (**e**). **c-f,** SKF-treated mice spend less time in the open arms (**c,** two-tailed unpaired *t*-test, *t*=2.106, \**p=*0.0405) and OFT center (**f,** two-tailed unpaired *t*-test, *t*=2.811, \*\**p=*0.0097) compared to vehicle-treated mice. **d-g,** No impairment of distance travelled in the EPM (**d,** two-tailed unpaired *t*-test, *t*=0.1008, *p=*0.9201) and OFT (**g,** two-tailed unpaired *t*-test, *t*=1.042, *p=*0.3078) between vehicle- and SKF-treated mice. **h,** Vehicle- or SKF-treated mice were perfused 90 minutes later for cFOS immunostaining. **i,** Representative confocal images of cFOS expression in the aIC and pIC for vehicle- and SKF-treated mice. **j,** SKF38383 i.p. injection increases the proportion of cFOS+ cells in the aIC (\*\*\**p*<0.0001), pIC (\*\*\**p*=0.0008) and NAc (\**p*=0.0234) compared to vehicle, with a greater effect in the aIC compared to pIC (\*\*\**p*<0.0001), NAc (\*\*\**p*<0.0001), BLA (\*\*\**p*<0.0001) and S1 (\*\*\**p*<0.0001) (Main effect Region: F_4,103_=34.2, \*\*\**p*<0.0001, Main effect Group: F_1,103_=49.2, \*\*\**p*<0.0001, with a Region x Group interaction: F_4,103_=6.54, \*\*\**p*<0.0001). **k,** Design and histological verification of intra-aIC bilateral activation and blockade of D1. Mice received a bilateral intra-aIC infusion of vehicle, SKF38393 (SKF, 0.25 mg/ml) or SCH23390 (SCH, 0.5 mg/ml) 6 minutes before the EPM. Each injection was mixed with a fluorescent marker (CTB488). **l-n,** Bilateral infusions of D1 receptor agonist (SKF) and antagonist (SCH) into the aIC decrease (\**p*=0.0123) and increase (\**p*=0.0259), respectively, the time mice spent in open arms (**m**) relative to vehicle, without affecting their total distance (**n**) travelled (One-way ANOVA, F_2,52_=11.37, \*\*\**p*<0.0001). **o,** Strategy to photoactivate D1 aIC neurons. **p,** Raster plot of action potential and peri-event stimulus histogram of the firing rate of one unit identified as a D1 aIC neuron. To identify light-responsive units, we compared each neuron z-scored firing rate during a baseline window (45 ms before light onset) to that during a 50 ms experimental window, starting at light onset, using a Wilcoxon signed-rank test (*p*<0.01). Units with a significant increase in firing (*p*<0.01) and a positive mean z-score during stimulation were classified as excited, whereas those showing a significant decrease (*p*<0.01) and a negative mean z-score were classified as inhibited. **q,** Heatmap representing the population z-score activity (top) of aIC neurons in response to the light stimulation (20 Hz, 1s, 5 ms pulses) and peri-event z-score firing of excited and inhibited (bottom) aIC neurons. **r,** Number of responsive units following the optogenetic activation of D1 aIC neurons in ChR2 and eYFP conditions (ChR2: 8/79 units excited (10%), 19/79 units inhibited (24%), and 52/79 units non-responding (66%); eYFP: 65/65 units non-responding (100%)). Absolute z-scored firing rate showed greater excitation compared to inhibition effect (two-tailed unpaired Mann-Whitney test, U=11, \*\*\**p*=0.0002). **s,** Strategy to optogenetically activate D1 aIC neurons (20 Hz, 1s, 5 ms pulses). **t-u,** Optogenetic activation of D1 aIC neurons decreases the time in open arms (**t**) compared to the light OFF period (two-tailed paired *t*-test, *t*=3.361, \**p*=0.0152), without affecting the total distance travelled (**u**). **b-d:** Vehicle (n=25 mice) and SKF (n=25 mice). **e-g:** Vehicle (n=13 mice) and SKF (n=13 mice). **h-j:** Vehicle (n=12 mice) and SKF (n=11 mice). **k-n:** Vehicle (n=29 mice), SKF (n=9 mice), SCH (n=17 mice). **o-r:** ChR2 (n=2 mice, 79 units) and eYFP (n=2 mice, 65 units). **s-u:** ChR2 (n=7 mice) and eYFP (n=8 mice). Data are: mean ± SEM.

To directly probe the contribution of anterior insula D1 receptors to anxiety regulation, we locally manipulated D1 signaling. Mice were bilaterally implanted with cannulae targeting the anterior insula and received infusions of the D1 agonist SKF38393 (0.25 mg/mL, **Fig. 3k and Extended Data Fig. 3a**), the D1 antagonist SCH23390 (0.5 mg/mL), or vehicle. In line with systemic D1 activation, local activation of D1 receptors in the anterior insula decreased time spent in open arms, whereas D1 blockade in the anterior insula increased time spent in open arms (**Fig. 3l-m and Extended Data Fig. 3b-c**), without affecting total distance travelled (**Fig. 3n**).

To assess the impact of D1+ neuron activation to local neural dynamics in the anterior insula, we combined *in vivo* electrophysiological recordings and optogenetic activation of D1+ anterior insula neurons (**Fig. 3o-p**). A 1s train of optogenetic stimulation at 20 Hz increased the firing of 10% of anterior insula neurons, while around 20% were inhibited, relative to eYFP controls (**Fig. 3q and Extended Data Fig. 3d**). Importantly, the magnitude of excitation exceeded inhibition by more than fourfold (**Fig. 3r**), suggesting a net excitatory effect on anterior insula network activity. These observations are consistent with the activation of the anterior insula induced by systemic SKF38393 injection and measured with cFOS expression. Finally, we assessed the behavioral impact of selective activation of D1+ anterior insula neurons using bilateral optogenetic activation. Remarkably, photoactivation of D1 anterior insula neurons reduced time spent in open arms (**Fig. 3s-t**) compared to controls, without altering total distance travelled by the animals (**Fig. 3u and Extended Data Fig. 3e-g**). Collectively, these findings demonstrate that activation of D1 receptors in the anterior insula promotes anxiety-related behaviors, supporting a causal anxiogenic role of D1 signaling in this region.

### Systemic D1 activation increases the proportion of anxiety- and salience-encoding units in the anterior insula

As systemic D1 activation increased anxiety-related behaviors and cFOS activation in the anterior insula, we sought to identify the impact of systemic D1 activation on neural coding of anxiety-related behaviors in this region. Thus, we conducted *in vivo* electrophysiology in the anterior insula of vehicle- and SKF38393-treated mice, which received systemic injections 30 minutes before EPM testing while being recorded (**Fig. 4a and Extended Data Fig. 4a**). Consistent with observations in non-implanted mice (**Fig. 3c**), systemic D1 activation decreased open arm entries (**Fig. 4b**) and reduced time spent in open arms compared to vehicle-treated mice, without affecting closed arm entries or total distance travelled (**Fig. 4b and Extended Data Fig. 4b**).

**Fig. 4:**
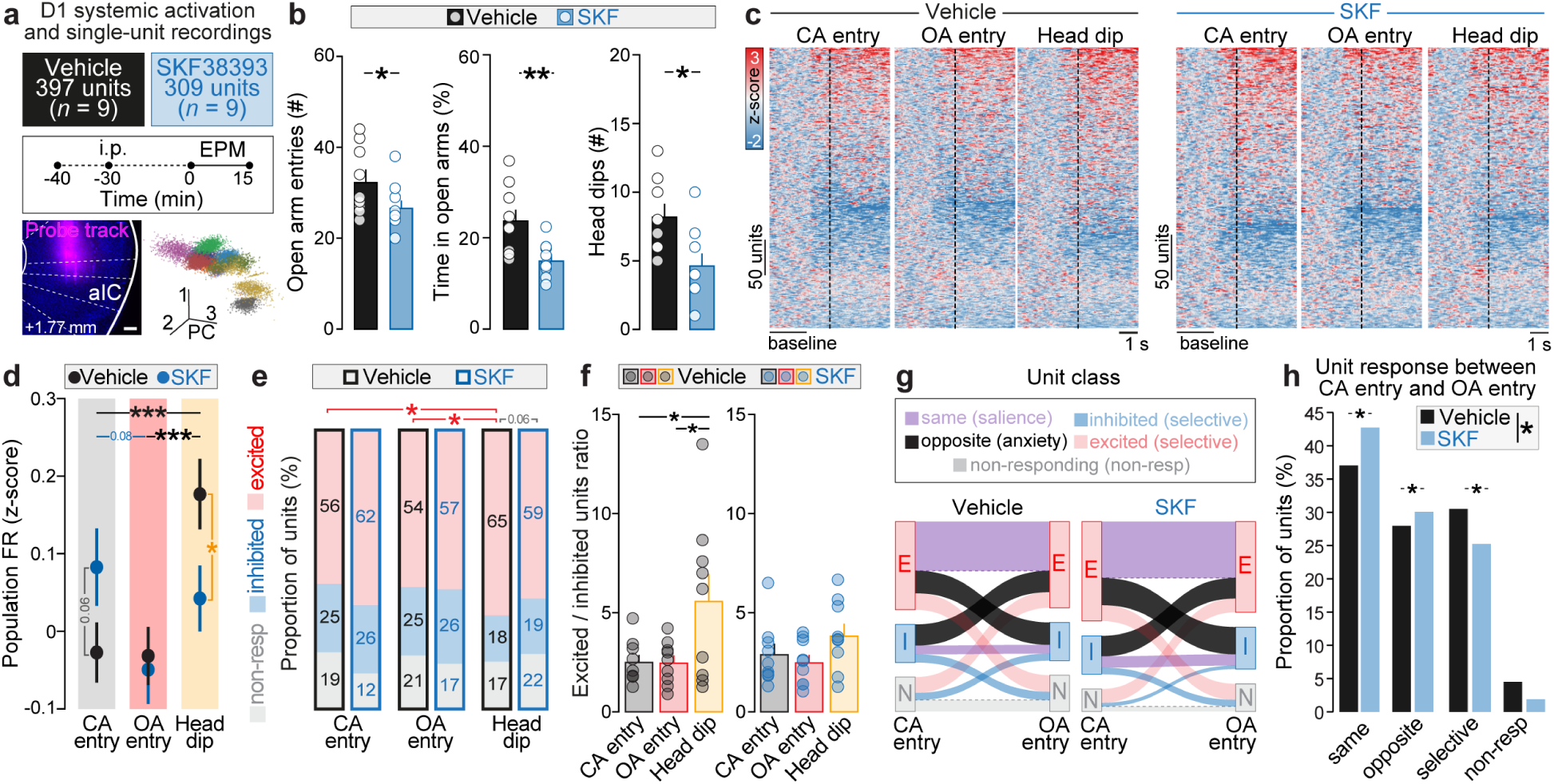
Single-unit activity of anterior insula neurons in response to anxiety-like behaviors. **a,** Extracellular recordings in the aIC using 16-channel electrodes in vehicle-treated (n=9, 397 units) and SKF38393-treated mice (n=9, 309 units). A 10-min baseline was recorded prior to i.p. injection. Representative histological confirmation of the electrode placement (top) and spike sorting representation from one mouse (bottom). **b,** SKF-treated mice perform less entries (one-tailed *t*-test, *t*=1.870, \**p*=0.0399) and spend less time in open arms (two-tailed *t*-test, *t*=3.144 \*\**p*=0.0063) and do less head dips (two-tailed *t*-test, *t*=2.856, \**p*=0.0114) compared to vehicle. **c,** Heatmaps of *z*-scored peri-event firing rate aligned to closed arm (CA) entry, open arm (OA) entry and head dips (HD) in open arms, in vehicle and SKF groups. Baseline window: [-3s to -1s]. **d,** Population z-scored firing rate of vehicle- and SKF-treated mice aligned to CA entry, OA entry and HD revealed a trend to increased firing rate at CA entry (*p*=0.0601), and decreased firing rate at HD (\**p*=0.0214) following D1 activation, relative to vehicle (two-way ANOVA: Main effect Trial type: F_2,2112_=6.61, \*\**p*=0.0014, Main effect Group: F_1,2112_=0.176, *p*=0.6753, with a Region x Group interaction: F_2,2112_=4.38, \**p*=0.0126). **e,** Proportions of ‘excited’, ‘inhibited’, and ‘non-responding’ units across trial types and groups. Generalized linear mixed models were performed on the proportion of excited, inhibited and non-responding units to assess the effects of trial type and group on unit class proportions. For excited units, in vehicle mice, the proportion of excited units was higher during HD compared to CA entry (\**p*=0.028), and compared to OA entry (\**p*=0.020). D1 activation tended to decrease the proportion of excited units during HD (*p*=0.059). No difference was observed for inhibited and non-responding units. **f,** Higher excited over inhibited ratio during HD relative to OA entry (\**p*=0.0394) and CA entry (\**p*=0.0421) in vehicle condition (one-way ANOVA, F_2,24_=8.206, \**p*=0.0222). No change observed following SKF injection (one-way ANOVA, F_4,40_=4.271, *p*=0.0685). **g,** Sankey diagrams showing transitions of units activity between CA and OA entry in vehicle and SKF mice. Units were classified as ‘excited’ (red), ‘inhibited’ (blue), or ‘neutral’ (gray) to CA entry and OA entry. Violet links indicate stable transitions (same), black links opposite transitions, and other links are colored by the transitional state. The width of each link is proportional to the percentage of units. **h,** Distribution of four outcome categories of unit response to CA entry and OA entry between Vehicle and SKF groups using a permutational multivariate analysis of variance (F=4.839, \**p*=0.038). Among unit classes, systemic D1 activation increases the proportion of ‘opposite’ units (\**p*=0.0489), ‘same’ units (\**p*=0.0264) and decreases the proportion of ‘selective’ units (\**p*=0.0442), relative to vehicle. Data are: mean ± SEM.

To dissect finer behavioral alterations, we used machine-learning-based tracking and supervised classification in the EPM using DeepLabCut^30^ and deepOF^31^. We extracted sniffing, climbing (against the closed walls), immobility, scanning (immobile environmental scanning with active head/nose movements), locomotion, instantaneous velocity and head dips. Head dips are discrete behavioral events in which mice extend their head beyond the open arm borders, and are a classical measure of risk assessment thought to be highly anxiogenic^32^. Systemic D1 activation reduced both the number of head dips and the time spent head-dipping (**Fig. 4b and Extended Data Fig. 4c-d**), whereas other quantified behaviors were unchanged **(Extended Data Fig. 4c-d)**, indicating that systemic D1 activation primarily affected anxiety-related behaviors rather than exploration, locomotion or arousal.

To assess anterior insula neural correlates of anxiety-related behaviors in the EPM, we aligned single-unit activity to three relevant behaviors: the entry to closed arms, the entry to open arms, and head dips in open arms (397 and 309 units for vehicle and SKF mice, respectively, **Fig. 4c**). In vehicle-treated mice, the average firing rate across recorded units after the onset of the three behavior types revealed that, firing rate was higher during head dips than at entries to either open or closed arms (**Fig. 4d**). On the contrary, SKF-treated mice did not exhibit significant differences in population firing rates across these behaviors. Notably, SKF treatment markedly decreased the firing rate during head dips and tended to increase it at closed arm entries (**Fig. 4d and Extended Data Fig. 4h**).

For each behavior, units were then classified as excited, inhibited or non-responding. Under vehicle conditions, the proportion of excited units was higher during head dips than at entries into open or closed arms (**Fig. 4e**). Systemic D1 activation did not alter these proportions, although a trend toward fewer excited units during head dips was observed in SKF-treated mice (**Fig. 4e**). In vehicle-treated mice, the excited-to-inhibited unit (E/I) ratio increased during head dips relative to both closed and open arm entries. After systemic D1 activation, the E/I ratio was similar across behaviors (**Fig. 4f**). We next categorized units into five classes based on their response to entry into closed and open arms: [1] same units, exhibiting similar activity at both arm types entry, [2] opposite units, excited at one entry and inhibited at the other, [3] excited-only units, excited at the entry of only one arm type, [4] inhibited-only units, inhibited at the entry of only one arm-type, and finally [5] non-responding units which have stable firing rates at the entrance of either arm type (**Fig. 4g**). The two largest classes were same units (∼37% and ∼43% for vehicle and SKF groups, respectively) and opposite units (∼28% and ∼30% for vehicle and SKF groups, respectively, **Fig. 4h and Extended Data Fig. 4i**). Systemic D1 activation increased the proportions of units with same and opposite responses at closed and open arm entries, and decreased the proportion of units with a selective response to one arm-type entry (**Fig. 4h**). When units were categorized by their response to open arm entry and head dips, or closed arm entry and head dips, systemic D1 activation did not alter the proportions of each category (**Extended Data Fig. 4j-k**).

Together, these results support a mixed selectivity among anterior insula neurons and indicate that systemic D1 activation reduces the number of selective units, while increasing the number of units with the same or opposite responses to entries into exposed versus safe zones. This reorganization suggests that systemic D1 activation increases the fraction of units encoding salience (‘same units’) and of units encoding anxiety-related behaviors (‘opposite units’).

### Systemic D1 activation disrupts anterior insula population dynamics during anxiety-related behaviors

Our single-unit analyses show that anterior insula population activity is increased during risk-assessment behaviors (**Fig. 4d-f**) and that systemic D1 activation alters the proportion of units responding to anxiety-related behaviors (**Fig. 4g-h**). Yet these observations do not establish how anterior insula neurons encode anxiety-related behaviors. To address this, we analyzed neural population activity in the anterior insula using principal component analysis (PCA) during anxiety-related behaviors (**Fig. 5a**). We constructed 250 ms population activity vectors across 6 seconds windows centered on anxiogenic behaviors (open arm entry and head dips) or protected behaviors (closed arm entry) in vehicle- and SKF-treated mice. In both groups, neural trajectory lengths computed in the first three principal components (PCs) were all longer than those obtained from shuffled data (**Fig. 5b-c**) and explaining between 8% and 14% of the variance (**Extended Data Fig. 5a**). In vehicle-treated mice, neural trajectory lengths were longer for anxiogenic behaviors (open arm entry < head dips) than for closed arm entry (black stars in **Fig. 5b-c and Extended Data Fig. 5b**), indicating more dynamic population coding of anxiogenic behaviors. Following D1 activation, this pattern was inverted: the neural trajectory was longest at closed arm entry and shortest around head dip (blue stars in **Fig. 5b-c and Extended Data Fig. 5b**). Consistently with this opposite pattern of neural dynamics in vehicle- and SKF-treated mice, neural trajectories were longer for closed arm entry and shorter for head dip, in SKF-compared to vehicle-treated mice (grey stars in **Fig. 5b-c and Extended Data Fig. 5b**).

**Fig. 5:**
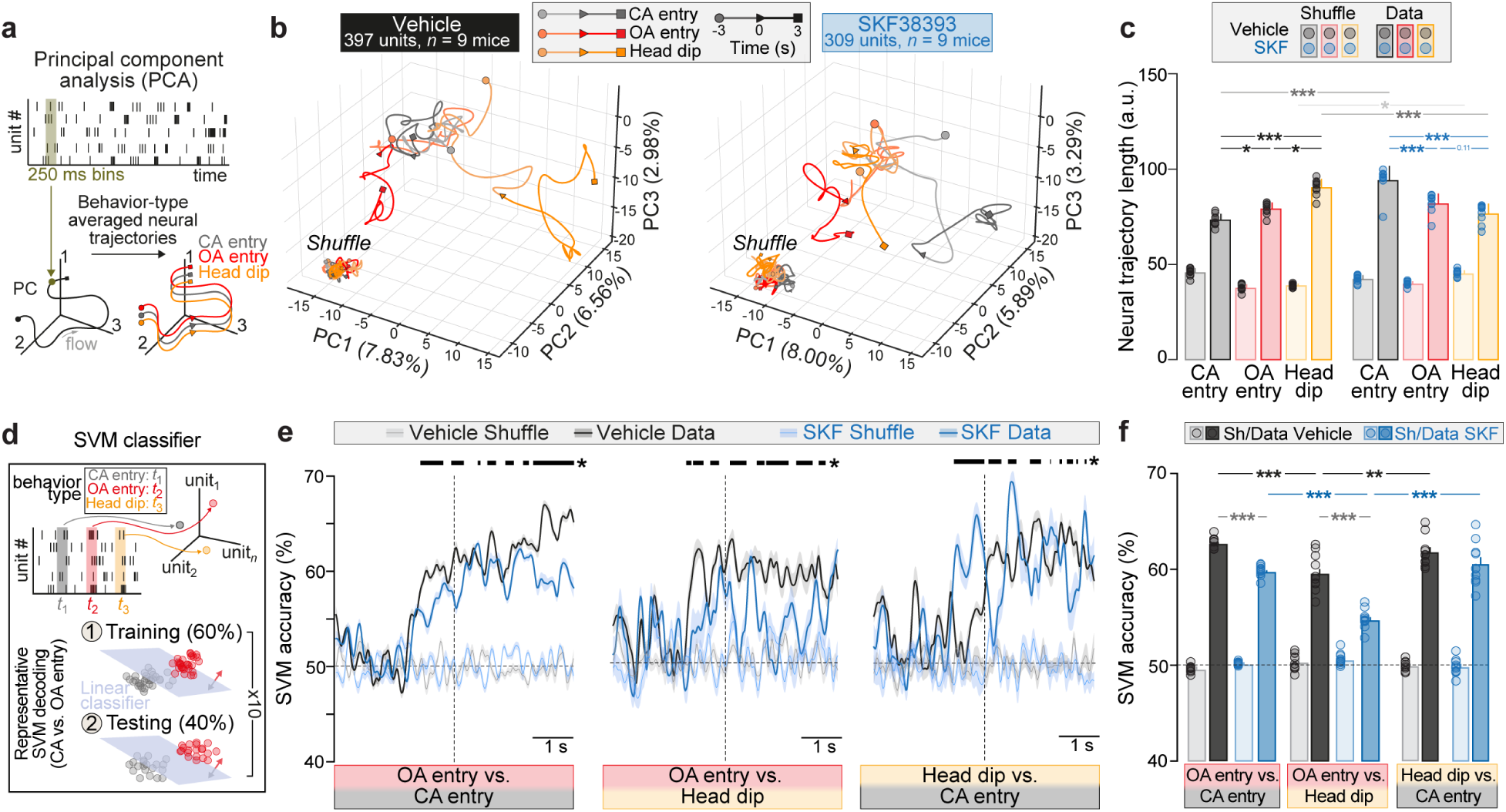
Anterior insula activity reveals coding specificity for risk-assessment behaviors. **a,** Population vectors (250 ms bins) contain spike counts of recorded aIC neurons for each trial. Principal component analysis (PCA) was applied to firing patterns across closed arm (CA) entry, open arm (OA), and head dips (HD) across 6 second behavioral snippets. **b,** Neural trajectories (Shuffle vs Data) for the three trial types projected onto the first three principal components explaining 17.37% (vehicle) and 17.18% (SKF) of the total variance (vehicle: PC1 = 7.83%, PC2 = 6.56% and PC3 = 2.98%; SKF: PC1 = 8%, PC2 = 5.89% and PC3 = 3.29%). a.u.: arbitrary units. **c,** For vehicle-treated mice, neural trajectories were longer for OA entry (\*\*\**p<*0.0001) and HD (\*\*\**p<*0.0001) compared to CA entry. In SKF-treated mice, neural trajectories were longer for CA entry (\*\*\**p*<0.0001) and shorter for HD (\*\*\**p*<0.0001), while no change was observed for OA entry. Neural trajectories from real data were statistically different from Shuffle. (3-way ANOVA: Main effect Event: F_2,96_=12.57, \*\*\**p*<0.0001, Main effect Data type: F_2,96_=3329, \*\*\**p*<0.0001, Main effect Group: F_2,96_=11.68, \*\*\**p*=0.0009, with an Event x Data type x Group interaction: F_2,96_=80.66, \*\*\**p*<0.0001). **d,** Illustration of the Simple Vector Machine (SVM) classifier to decode the activity of each behavior at a given time point. **e,** Temporal evolution of the SVM decoding accuracy from Shuffle and Data in vehicle and SKF mice for OA entry vs. CA entry (left), OA entry vs. Head dip (middle) and Head dip vs. CA entry (right). The dashed line represents the chance level. The thick line represents the permutation test performed on each bin during 1000 iterations (\**p*<0.05). **f,** Mean of the SVM performances from the onset of the event (0 to 3s) between vehicle and SKF, showing a decreased accuracy to decode between CA and OA entries in SKF-treated mice (\*\*\**p*<0.0001) and OA entries and head dips (\*\*\**p*<0.0001), while no change of decoding accuracy was observed between head dip vs CA entry, relative to vehicle (2-way ANOVA: Main effect Group: F_1,36_=47.9, \*\*\**p*<0.0001, Main effect Data type: F_1,36_=63.6, \*\*\**p*<0.0001, with a Group x Data type interaction: F_1,36_=39.5, \*\*\**p*<0.0001). Data are: mean ± SEM.

To directly quantify how anterior insula population activity encodes anxiety-related behaviors, we trained a linear support vector machine (SVM) classifier. The decoder was trained on 60% of the data and tested on the remaining 40% to distinguish between open arm entry, closed arm entry and head dips (Methods, **Fig. 5d**). For shuffled datasets, decoding performance remained at chance level (**Fig. 5e-f and Extended Data Fig. 5c-d**). In vehicle-treated mice, the accuracy to decode open and closed arm entries reached 62.6%, similar to the accuracy to decode closed arm entry and head dips (61.7%) while the accuracy to decode open arm entry vs head dips was lower (59.5%, black stars, **Fig. 5e-f**). Systemic D1 activation reduced the accuracies to decode open and closed arm entries, and to decode open arm entry and head dips (**Fig. 5e-f**). Finally, when decoding all three behaviors simultaneously, decoding accuracy in vehicle mice was 11% higher than chance, which was decreased to 7.2% higher than chance following D1 activation (**Extended Data Fig. 5c-d**). Together, PCA and decoding analyses indicate that under control conditions, anterior insula coding is more dynamic for risk-assessment behaviors. Systemic D1 activation increases neural dynamics for entries to a safe space, while decreasing neural dynamics of risk assessment, leading to a reduction of information content (**Extended Data Fig. 7d**).

### Non-linear dimensionality reduction reveals that systemic D1 activation alters neural representations of anxiety-related behaviors and spaces in the anterior insula

Linear dimensionality reduction using PCA indicated that anterior insula population activity differentiates protected and anxiogenic behaviors, although the first three PCs explained only a modest fraction of total variance. To further probe nonlinear structure in population activity, we applied the recently developed machine-learning algorithm, CEBRA^33^ (**Fig. 6a** and Methods) and trained it solely on single-unit spiking (CEBRA-Time model). Embeddings were computed in a three-dimensional latent space from 50 ms binned activity, and behavioral labels were overlaid post hoc. To quantify how anxiety-related information is organized in the latent space, we computed two complementary metrics on CEBRA latent clusters: the barycenter distance, measuring the separation between cluster centroids, and the Shannon entropy, quantifying how compact or dispersed a cluster is (**Fig. 6a**, see Methods for details).

**Fig. 6:**
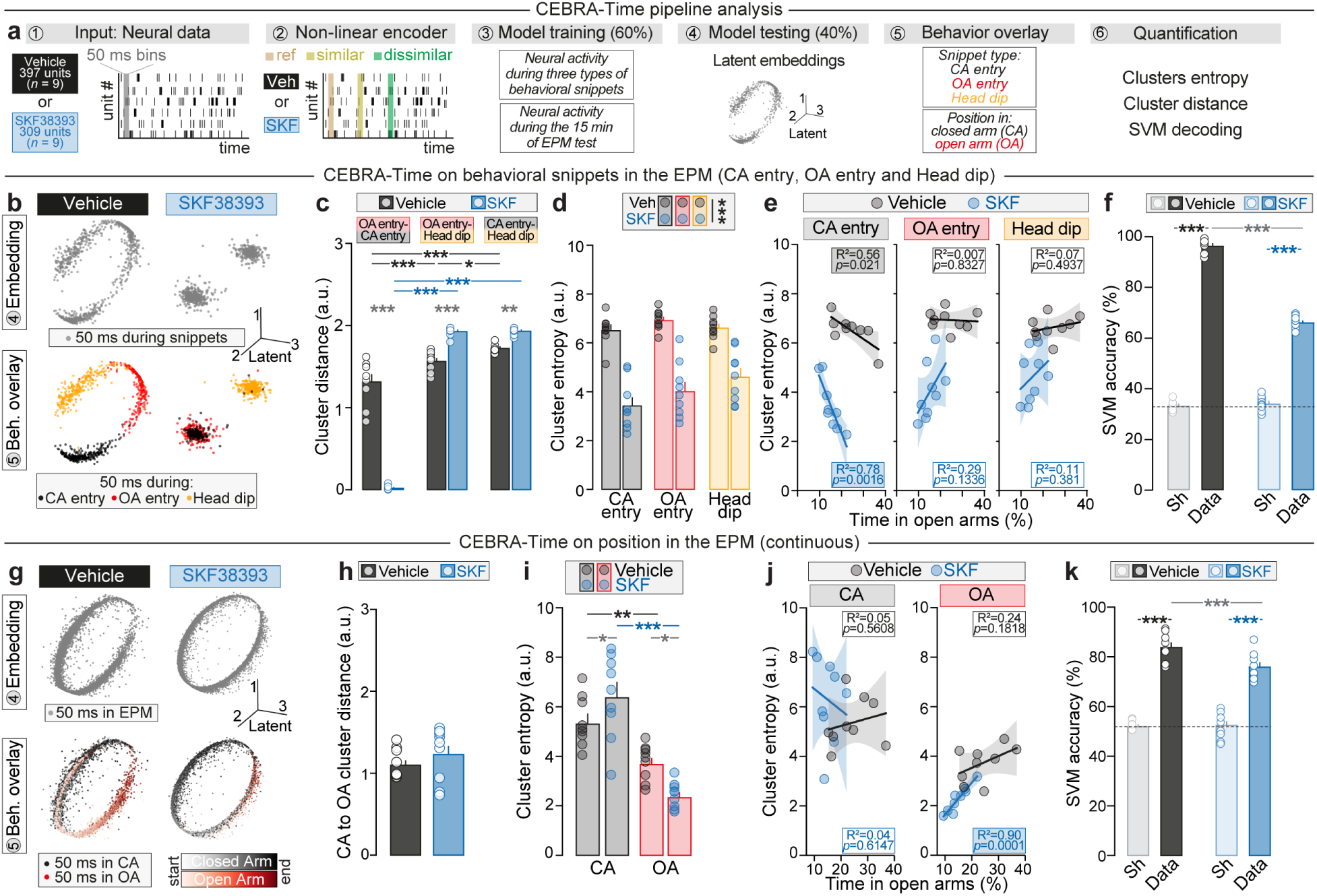
Non-linear dimensionality reduction reveals hidden features of anxiety in the anterior insula. **a,** CEBRA pipeline applied to aIC single-units. Population spike trains were binned (50 ms) and projected into a low-dimensional embedding space using contrastive learning. **b,** Representative CEBRA embeddings for vehicle- and SKF-treated mice representing behavioral snippets. Latent embeddings were plotted in the first three latent spaces and post hoc color coded by trial types (CA entry: black, OA entry: red and head dips (HD): orange). **c,** Barycenter distance of different clusters for vehicle and SKF groups. D1 activation decreased (\*\*\**p*<0.0001) the distance between OA entry and CA entry clusters, increased (\*\*\**p*<0.0001) the distance between OA entry and head dip clusters, and increased (\*\*\**p*<0.0001) the distance between CA entry and head dip clusters (Two-way ANOVA: Main effect Snippets: F_2,48_=468, \*\*\**p*<0.0001, Main effect Group: F_1,48_=48.9, \*\*\**p*<0.0001, with a Snippets x Group interaction: F_2,48_=236, \*\*\**p*<0.0001). **d,** D1 activation decreased the Shannon entropy of all clusters without specific changes within groups (Two-way ANOVA: Snippets: F_2,48_=2.93, *p*=0.0629, Main effect Group: F_1,48_=141, \*\*\**p*<0.0001, with no interaction). **e,** For CA entry latent cluster only, its entropy negatively correlates with the time spent in open arms in vehicle-treated mice (two-tailed Pearson correlation: R²=0.56, \**p*=0.021), and correlates negatively even more following D1 activation (two-tailed Pearson correlation: R²=0.78, \*\**p*=0.0016). No difference was observed for OA entry and HD latent clusters (two-tailed Pearson correlations). **f,** A SVM classifier reached above 95% of accuracy decoding between the three trial types in vehicle group (\*\*\**p*<0.0001), while the decoding decreased to 65% of decoding accuracy (***p<0.0001) following SKF injection (Two-way ANOVA: Main effect Data type: F_1,32_=14.8, \*\*\**p*<0.0001, Main effect Group: F_1,32_=484, \*\*\**p*<0.0001, with a Data type x Group interaction: F_1,32_=4.95, \**p*=0.0333). **g,** Representative CEBRA embeddings for vehicle- and SKF-treated mice representing the continuous EPM recording. Latent embeddings were plotted in the first three latent spaces and post hoc color coded with the mouse position in the EPM (CA: black, OA: red). **h,** No change of barycenter distance between OA and CA latent clusters following D1 activation (two-tailed *t-*test, *t*=1.033, *p*=0.3170). **i,** Shannon entropy increased for CA latent cluster (\**p*=0.0462) but decreased for OA latent cluster (\**p*=0.0133) after SKF injection compared to vehicle (Two-way ANOVA: Main effect Arm: F_1,16_=60.3, \*\*\**p*<0.0001, Group: F_1,16_=0.152, *p*=0.7015, with an Arm x Group interaction: F_1,16_=10.8, \*\**p*=0.0046). **j,** Shannon entropy for the CA latent cluster did not correlate with the time spent in open arms in both vehicle and SKF groups (two-tailed Pearson correlation, vehicle: R²=0.05, *p*=0.5608, SKF: R²=0.04, *p*=0.6147). For the OA latent cluster, only after D1 activation that the OA latent cluster entropy correlates with the time mice spent in open arms (two-tailed Pearson correlation, vehicle: R²=0.24, *p*=0.1818, SKF: R²=0.90, \*\*\**p*=0.0001). **k,** SVM decoding of OA vs. CA position achieved above 90% accuracy in vehicle mice but decreased to 75% (\*\*\**p*<0.0001) after SKF injection (two-way ANOVA: Main effect Data type: F_1,32_=14.8, \*\*\**p*=0.0005, Main effect Group: F_1,32_=484, \*\*\**p*<0.0001, with a Data type x Group interaction: F_1,32_=4.95, \**p*=0.0333). **c-d, f, i, k**: Two-way ANOVA and Tukey’s post hoc tests were performed. Data are: mean ± SEM.

We first applied the CEBRA-Time to neural snippets of anterior insula neural activity aligned to behaviors (**Extended Data Fig. 6a**). In vehicle-treated mice, three clearly separated latent clusters corresponding to closed arm entry, open arm entry and head dips, indicate distinct population codes for protective (closed arm entry) and risk-assessment behaviors (open arm entry and head dips). Strikingly, following systemic D1 activation, latent embedding structure was completely reorganized, with substantial overlap between closed arm entry and open arm entry clusters (**Fig. 6b**). Accordingly, D1 activation nearly abolished the cluster separation between closed and open arm entries, as reflected by a barycenter distance close to zero. On the contrary, systemic D1 activation increased the separation between the head dip cluster and both open and closed arm entry clusters (**Fig. 6c**). Interestingly, systemic D1 activation also reduced the entropy of closed and open arm entry clusters, and of the head dip cluster (**Fig. 6d**). While cluster distances were independent of anxiety levels, estimated by time spent in open arms (**Extended Data Fig. 6b**), closed arm entry cluster entropy positively correlated with anxiety levels in mice with or without D1 activation (**Fig. 6e**). However, when pooling mice of both groups, the entropy of open arm entry and head-dip clusters were positively correlated with the time mice spent in the open arm, meaning that the more anxious mice were the lower the entropy of those cluster was (**Extended Data Fig. 6c**).

Finally, SVM decoding of the three behaviors yielded accuracies above 95% in vehicle mice, which decreased of 30% after D1 activation (**Fig. 6f**). Similar results were obtained using k-nearest neighbors (kNN) decoding (**Extended Data Fig. 6d**), supporting the hypothesis that D1 activation disrupts anxiety-associated representations in the anterior insula. Altogether, CEBRA-Time analysis of behavioral snippets reveal that D1 activation strongly alters neural population dynamics during anxiety-related behaviors, by both blurring the distinction between neural codes of protective and risk-assessment behaviors, and by magnifying the segregation of these codes and the one of high risk-taking behavior.

As behavioral snippets only capture a portion of the neural activity rather than representing the full recording, we next applied CEBRA-Time on the entire EPM recording. When latent embeddings were color-coded by mouse position, we observed segregated clusters depending on the mouse position in safe and exposed spaces (**Fig. 6g** and **Extended Data Fig. 6e-f**). The distance between open and closed arm clusters did not change after systemic D1 activation (**Fig. 6h**), and this distance was independent of the level of anxiety in both vehicle- and SKF-treated mice (**Extended Data Fig. 6i**). Interestingly, systemic D1 activation increased the entropy of the closed arm cluster, while decreasing the entropy of the open arm cluster (**Fig. 6i**). Importantly, matching the number of open arm and closed arm positions yielded similar entropy values, confirming that the increased dispersion of the closed arm cluster is independent of the increased time spent in closed arms induced by systemic D1 activation (**Extended Data Fig. 6j**). Moreover, the open arm cluster entropy correlated negatively with anxiety levels of SKF-treated mice (**Fig. 6j**). This correlation was also present when combining vehicle- and SKF-treated mice, indicating that more anxious mice exhibit sharper, more consistent representations of open arm occupancy in the anterior insula (**Extended Data Fig. 6j**).

Finally, high SVM decoding accuracy for closed arm versus open arm in vehicle mice was decreased in SKF-treated mice (**Fig. 6k**, similar to kNN decoding in **Extended Data Fig. 6k**), supporting that systemic D1 activation decreases the information content of neural firing in an anxiogenic environment (**Extended Data Fig. 7d**).

Together, these complementary metrics of CEBRA latent embeddings reveal neural features of anxiety in anterior insula population activity. At the individual behavior level, D1 activation disrupts latent representations of entries into safe and exposed compartments. At the space level, coding reliability for open arm scales with anxiety levels. Shannon entropy and barycenter distance thus provide quantitative metrics demonstrating that systemic D1 activation reshapes anterior insula population code for anxiety across event-based and continuous timescales.

## Discussion

The anterior insula and dopaminergic transmission both regulate anxiety in humans and in preclinical models, yet the dopaminergic impact on anterior insula function has remained unknown. Using a multi-faceted approach, we identify a D1-dependent mechanism by which dopamine shapes anterior insula neural dynamics for protective and risk-assessment behaviors.

### VTA dopaminergic innervation of the anterior insula

We found that both anterior and posterior insula receive dopaminergic inputs from the VTA and SNc, extending previous work showing both nuclei innervate the anterior insula^34^. Additionally, we show that VTA^DA^ neurons preferentially project to the anterior insula, whereas SNc^DA^ terminals are more uniformly distributed in the rostro-caudal axis of the insula, except in layer 6 where they are denser in the posterior insula. Our findings are consistent with the broader view that midbrain dopamine neurons are organized into projection-defined subpopulations with distinct input-output connections and behavioral roles^35,36^. While SNc^DA^ neurons classically regulate sensorimotor processes via striatal targets^37^, VTA^DA^ neurons are critical for processing both positive and negative valence^38–40^, including anxiety^41^. In this context, the preferential VTA^DA^ innervation of the anterior compared to posterior insula, combined with the established anxiogenic role of the anterior insula^1,2^, supports the idea that dopamine released from VTA^DA^ neurons could be a key contributor to anterior insula functions in anxiety.

### Topography of D1 and D2 neurons of the anterior insula

We showed that D1+ neurons outnumber D2+ neurons in both anterior and posterior insula, consistent with human^42^ and preclinical studies^23^. Along the rostro-caudal axis, we identified a higher density of D1+ neurons in the anterior compared to the posterior insula, suggesting a prominent influence of D1 modulation in the anterior insula. Along the medio-lateral axis, unlike in the mPFC where D1+ neurons are enriched in deeper layers^43–45^, in the anterior insula, D1+ neurons are preferentially located in layer 2/3. This distribution suggests a specialization for modulating cortico-cortical circuits and indirectly shaping layer 5 projection neurons^46^. Although the proportion of D1+ or D2+ neurons is higher among inhibitory neurons, as reported in the mPFC^44,47^, in absolute numbers D1+ excitatory and D2+ excitatory neurons are more than threefold more numerous than D1+ inhibitory and D2+ inhibitory neurons. This indicates that glutamatergic neurons represent the primary target of D1 control in the insula. Hence, the very limited *Drd1/Drd2* co-expression (<10%) supports largely segregated D1 and D2 circuits, reminiscent of the functional dichotomy described in the mPFC^48^, ventral hippocampus^6^, and striatum^49–51^. Finally, long-range mapping of anterior and posterior insula D1+ neurons revealed dense projections to ipsilateral amygdala nuclei and contralateral insula, closely mirroring the projection pattern of insula glutamatergic neurons^2,29^. This consistency supports the existence of D1 projection-defined subpopulations that may differentially contribute to anxiety control, with D1 anterior insula-BLA neurons being strong candidates given the established role of anterior insula-BLA neurons in anxiety^2^. Future work will delineate the contribution of selective anterior insula D1 circuits in encoding anxiety states.

### Brain region and cell-type selectivity of dopaminergic modulation

Dopamine signaling is region-specific and modulates a wide range of behaviors^52^. Only two previous studies have reported increased dopamine signals during exploration of anxiogenic environments, recorded in the ventral hippocampus^6^ and in the interpeduncular nucleus^53^. Here, we report for the first time increased dopamine signals in the insula during exposure to exposed spaces, with stronger signals in the anterior insula relative to the posterior insula. This presence of increased dopamine signals in exposed areas, across multiple brain regions, supports a model in which dopamine acts a key modulator of a distributed anxiety network, integrating both anterior and posterior insula.

Importantly, although dopamine targeted both D1+ and D2+ anterior insula neurons during exposure to exposed spaces, dopamine signals preferentially increased onto D1+ neurons upon entry of these exposed spaces. Consistently, only dopamine signaling onto D1+ neurons of the anterior insula scaled with the level of anxiety of mice. These findings suggest that D1+ neurons of the anterior insula encode anxiety magnitude while D2+ neurons may preferentially signal contextual or state-dependent aspects of anxiety. Together, these results indicate that D1+ and D2+ anterior insula neurons encode distinct components of anxiety, potentially mediated by differential dopaminergic inputs, such as distinct sources (VTA versus SNc) and/or divergent laminar projections preferentially targeting D1+ and D2+ neurons.

### Systemic and insula D1 activation increase anxiety-related behaviors

We repeatedly found that systemic D1 activation elicited robust anxiogenic effects, as recently published in adolescent mice^54^. Conversely, a seminal study had found that systemic D1 blockade produce anxiolytic effects as shown by increased head-dipping in mice^55^. In line with these previous studies, our findings reveal a systemic and bidirectional D1-dependent control of anxiety. Furthermore, systemic D1 agonism induced neural activation in the anterior and posterior insula, as well as NAc, consistent with other studies reporting activation in NAc and PVN^56,57^. Surprisingly, BLA activation remained unchanged, despite high D1 expression^58^ and previously described anxiogenic influence of D1 in this region^4^, suggesting that BLA recruitment might be more transient or context-dependent. Importantly, we observed the highest neural activation in the anterior insula, pinpointing this insula section as a crucial actor of systemic D1-mediated anxiogenesis. We further demonstrated using local D1 activation/blockade or optogenetic D1 neuron activation, that both D1 receptors and D1+ neurons of the anterior insula have anxiogenic properties. Optogenetic activation of D1+ neurons increased anxiety, consistent with recent findings in the ventral hippocampus^6^. However, this contrasts with studies showing that D1 blockade in the dorsal hippocampus^59^ or ventral pallidum^60^ leads to increased or unchanged anxiety, respectively. Moreover, optogenetic activation of D1+ neurons in the VTA has been reported to decrease anxiety^9^. These region-specific effects (**Extended Data Fig. 7c**) underline that behavioral outcomes critically depend on which dopaminoceptive circuit is engaged (**Extended Data Fig. 7c**). Integrating our behavioral, neural activation, pharmacological and optogenetic observations with previous findings, we propose that systemic administration of a selective D1 agonist engages a distributed network of which the anterior insula is a major target. While some of the behavioral effects following systemic injection may be mediated by D1 activation outside the insula, it is reasonable to infer that D1 activation within the anterior insula strongly contributes to anxiogenic effects induced by systemic administration of a D1 agonist.

### Systemic D1 activation modulates anterior insula single-unit responses to anxiety

Anterior insula neurons form a heterogeneous pool distributed across response types during protective and risk-assessment behaviors, as reported in a stress-related paradigm^20^. Systemic D1 activation increased the proportion of the two most prominent pools of neurons responding to those three behaviors (closed arm entry, open arm entry, head dip). Specifically, D1 activation increased the number of units exhibiting either similar responses across entries to both safe and exposed spaces, or opposite responses to these contexts, suggesting additional recruitment of salience-encoding and anxiety-encoding units, respectively. Interestingly, more than twice more extra salience-encoding units were recruited after systemic D1 activation compared to extra anxiety-encoding units. In parallel, decoding accuracy for entries into safe or exposed spaces decreased following systemic D1 activation, indicating that population activity in the anterior insula during these behaviors becomes less discriminable. We hypothesize that the enhanced number of salience-encoding units could partially mask information from anxiety-encoding units that distinguish protective from risk-assessment behaviors, reducing the clarity of the neural representation and contributing to an overall increase in anxiety.

### Nonlinear methods to dissect neural population dynamics

Principal component analysis of anterior insula activity allowed us to visualize variability in population firing constructing neural trajectories computed across all mice. In contrast, the non-linear model CEBRA-Time^33^ preserved inter-individual information in population coding, enabling direct comparisons across individuals. Compared to previous CEBRA applications^61,62^, our approach focuses on the intrinsic structure of population dynamics without using behavioral information^63^. After representing the neural embeddings of each mouse in the latent space, we then overlaid, post hoc, in which behavioral state or physical location mice were for each analyzed time bin. Interestingly, this approach revealed that time bins of anterior insula activity organized into distinct latent clusters, each containing bins of activity sharing similar firing structure, as well as behavioral features. We further extended this data representation by introducing explicit quantitative metrics of latent clusters geometry: the Shannon entropy and barycenter distance. The Shannon entropy captures how compact or dispersed a latent cluster is, which is a proxy for the coding variability of the associated behavioral state. The barycenter distance measures separation between two latent clusters, estimating the neural divergence of pairs of behavioral states. These metrics open avenues to quantify neural codes of virtually any type of behavior, emotional state, sensory perception or motor action. Together, PCA and CEBRA offer complementary tools to quantify neural population dynamics at mouse populations and inter-individual levels, respectively.

### Systemic D1 activation reorganizes anterior insula firing geometry of anxiety-related behaviors and spaces

Under control conditions, high neural decoding accuracy of CEBRA in differentiating risk-assessment and protective behaviors, as well as exposed and protected spaces, supports the hypothesis that the anterior insula generates distinct population codes for high and low anxiety states. Notably, systemic D1 activation profoundly reorganizes the latent geometry of anterior insula activity. Beyond inducing a general decrease in coding variability for the three behaviors analyzed, it markedly reduced the distance between clusters associated with entries into safe or exposed areas, behaviors which share similar sensorimotor components, yet differ in inferred threat value. In addition, in both control and D1-activated mice, higher anxiety was associated with larger coding variability within the cluster linked to the entry into protected spaces. This suggests that anterior insula geometry tracks individual differences in avoidant behaviors.

Interestingly, systemic D1 activation also induced opposite effects on coding variability for protected and exposed spaces. Specifically, it increased the variability of safe spaces representation, while increasing the reliability of anxiogenic spaces representation. Notably, systemic D1 activation installed a positive correlation between anxiety levels and reliability of exposed spaces representation, indicating that coding reliability of anxiogenic contexts scales with the level of anxiety in mice. This aligns with the idea that dopamine through D1 can modulate threat valuation and the assignment of negative value to cues and contexts^64–67^.

Finally, consistent with linear decoding analyses, systemic D1 activation also markedly reduced the accuracy to decode entries to safe and exposed spaces, and to decode those spaces, using a non-linear approach. Altogether, these reduced decoding accuracies supports the idea that D1 signaling reduces neural discrimination of contextual value (safety vs. threat) under similar sensorimotor features. In this framework, D1 activation would blur the population activity patterns that normally distinguish safe from exposed contexts, impairing the anterior insula ability to appropriately differentiate safety from threat representations, which could in turn increase anxiety.

### Conclusions

In summary, we provide the first evidence of a D1-mediated control of anxiety in which the anterior insula plays a crucial role. This key brain structure receives dopaminergic innervation preferentially from the VTA and expresses a higher proportion of D1+ neurons than other insular territories. Anterior insula D1 modulations induced a bidirectional control over anxiety, with activations of D1 receptors or D1+ neurons both increasing anxiety-related behaviors. At the coding level, systemic D1 activation dampens the coding dichotomy between protective and risk-assessment behaviors which share similar sensorimotor components, as well as between safe and exposed contexts. We hypothesize that this alteration of anterior insula coding selectivity could blur the mental delineation between threat and safety, which might contribute to the observed increased anxiety. In parallel, systemic D1 activation linked the coding reliability of exposed spaces to mice level of anxiety, suggesting a direct contribution of anterior insula D1 activation to neural coding of both anxiety-related behaviors as well as anxiety levels. Finally, our work lays out an innovative approach using nonlinear deep learning methods to dissect neural population dynamics underlying specific behaviors and mental states. By defining how dopamine modulates insula control of anxiety in mice, our work will serve as a starting point to understand how dopaminergic modulation contributes to anxiety disorders.

## Methods

### Animals and housing conditions

Adult male and female mice, at least 10 weeks old, were used for all experiments. The cohorts included wild-type C57Bl/6J mice, transgenic D1-cre (Tg(Drd1a-cre)AGsc/KndlJ, JAX 030329), DrD2-cre (Tg(DrD2-cre)ER44Gsat, MMRRC 32108), D1-cre*Ai14, D2-cre*Ai14, and DAT-cre mice (B6.SJL-Slc6a3tm1.1(cre)Bkmn/J, JAX 006660). Animals were maintained under controlled temperature and humidity on a reversed 12 h light/dark cycle, with ad libitum access to food and water. All housing, treatments, and experimental procedures complied with French governmental regulations and were approved by the local ethical committee and the Ministry of Education, Research, and Innovation (APAFIS #30247), following European Communities Council Directives.

### Stereotaxic surgeries

Before surgery, mice received Metacam (5 mg/kg) and all procedures were performed under aseptic conditions. Anesthesia was induced with isoflurane (5%) and maintained at 1.5-2% in a stereotaxic frame, with body temperature controlled using a heating pad. Postoperative health and weight were monitored for three days. Retrograde tracers or viral vectors were delivered by intracranial injection using glass micropipettes (3-000-203-G/X, Drummond Scientific) pulled with a PC-100 puller (Narishige). Injections were made at a rate of 1 nL.s^-1^ with a Nanoliter 2020 injector (WPI). After injection, the pipette was left in place for 5 minutes to facilitate diffusion before being slowly withdrawn, after which the skin was sutured. Following surgery, mice were kept under a heat lamp until they fully recovered from anesthesia and then returned to their home cages. Animals were allowed 1 week for retrograde tracer transport or 3-5 weeks for viral expression before experiments.

<u>Dopaminergic inputs to the insular cortex</u> were mapped using complementary tracing strategies. Retrograde tracers (CTB conjugated to Alexa Fluor 488 or 555) were injected into either the anterior or posterior insular cortex (aIC or pIC, counterbalanced). In DAT-cre mice, anterograde labeling was achieved by injecting Cre-dependent AAV_9_ vectors encoding eYFP (AAV_9_-EF1α-DIO-eYFP) or mCherry (AAV_9_-EF1α-DIO-mCherry) into the ventral tegmental area (VTA) and substantia nigra pars compacta (SNc).

For quantification of synaptic contacts in D1 insula neurons, D1-cre mice received injections of a 1:2 mixture of AAV_9_-EF1α-DIO-eYFP and AAV_8/2_-EF1α-DIO-Synaptophysin-mCherry into the aIC and pIC.

<u>For combined optogenetic manipulations and fiber-photometry recordings of dopamine signal</u>, DAT-cre mice received AAV_9_-hSyn-gDA3m (green GRAB-DA3m) injections into the aIC, and AAV_8_-EF1a-DIO-rsChRmine-oScarlet (or a control vector, AAV_9_-hSyn-DIO-mCherry) into the VTA. An optical fiber (400 μm diameter, NA 0.39, >90% efficiency) mounted in a 1.25 mm ferrule was implanted 50 μm above the aIC and 300 μm above the VTA with a 10° angle, at following stereotactic coordinates: ML ± 0.9 mm, AP −3.2 mm, DV −4.20 ± 0.05 mm from Bregma (Paxinos). Fibers were secured with biocompatible cement and resin. Mice recovered in their home cages for 5 weeks to allow for viral expression.

<u>For fiber photometry recordings of dopamine signal</u>, either AAV_1_-hSyn-dLight1.1 was injected into the aIC and pIC, or AAV_9_-hSyn-DIO-gDA3m was injected into the aIC of D1-cre and D2-cre mice. An optical fiber (400 μm diameter, NA 0.39, >90% efficiency) mounted in a 1.25 mm ferrule was implanted 50 μm above the injection site and secured with biocompatible cement and resin. Mice recovered in their home cages for 3 weeks to allow for viral expression.

<u>For pharmacological experiments</u>, bilateral stainless-steel guide cannulae (4 mm, 26G; Phymep) were implanted 1 mm above the aIC. Cannulae were fixed with dental cement as for fiber photometry implants, using the following coordinates relative to Bregma (mm): AP +1.7, ML ±3.1, DV −2.5. After cementation, protective caps (Phymep) were secured to the guide cannulae.

<u>For optogenetic activation of D1 aIC neurons</u>, D1-cre mice received bilateral injections of AAV_9_-EF1a-DIO-hChR2-eYFP or the control viral vector AAV_9_-EF1a-DIO-eYFP into the aIC. Optical fibers (300 μm diameter, NA >0.80 efficiency) mounted in 1.25 mm ferrules were implanted 300 μm above the aIC injection site and secured with biocompatible cement and resin. Mice recovered for 4 weeks to allow for viral expression before behavioral testing.

**Table 1.** Details of coordinates, anatomical tracing and viral vectors.

| Fig. | Coordinates<br>AP/ML/DV (mm) | Viral vector | V(nL) | Titer<br>(vg/mL) | Provider |
| --- | --- | --- | --- | --- | --- |
| 1a | **aIC:** +1.7 / +3.1 / -3.5<br>**pIC:** -0.35 / +4.0 / -4.2 | CTB AF488 or AF555 | 250 |  | Fisher Scientific<br>488 (11544267)<br>555 (11554267) |
| 1e | **VTA:** -3.2 / +0.4 / -4.35<br>**SNC:** -3.16 / +1.5 / -4.2 | AAV<sub>9</sub>- EF1α-DIO-eYFP<br>AAV<sub>9</sub>- EF1α-DIO-mCherry | 400 | 6.69.10<sup>13</sup><br>(diluted<br>1:10) | Addgene<br>(27056-AAV9)<br>(50462-AAV9) |
| S2n | **aIC:** +1.7 / +3.1 / -3.5<br>**pIC:** -0.35 / +4.0 / -4.2 | AAV<sub>8/2</sub>-hEF1α -DIO-Synaptophysin-mCherry | 250 | 2.5.10<sup>13</sup> | Massachusetts<br>General<br>Hospital<br>AAV-RN1 |
| 1i | **VTA:** -3.4 / 0.35 / -4.0<br>(10°) | AAV<sub>8</sub>- EF1α-DIO-rsChRmine-<br>oScarlet<br>AAV<sub>9</sub>- EF1α-DIO-mCherry | 400 | ≥ 10<sup>13</sup><br>(diluted<br>1:10) | Deisseroth Lab |
| 1m-p | **aIC:** +1.7 / +3.1 / -3.5<br>**pIC:** -0.35 / +4.0 / -4.2 | AAV<sub>1</sub>.hSyn.dLight1.1 | 250 | ≥ 7.10<sup>12</sup> | Addgene<br>(111066-AAV1) |
| 2d-i | **aIC:** +1.7 / +3.1 / -3.5 | AAV<sub>9</sub>.hSyn.DIO.gDA3m | 250 | ≥ 2.10<sup>12</sup> | From Li Lab<br>Vector Builder |
| 3o-u | **aIC:** +1.7 / +3.1 / -3.5 | AAV<sub>9</sub>- EF1α-DIO-hChR2-eYFP<br>AAV<sub>9</sub>- EF1α-DIO-eYFP | 250<br>250 | ≥ 6. 10<sup>13</sup><br>(diluted<br>1:10) | Addgene<br>(35509-AAV9)<br>(27056-AAV9) |
| Red Dil used to coat electrodes before implantation |  |  |  |  | ThermoFisher<br>(V22885) |

<u>For extracellular recordings</u>, a 16-wire electrode array (4×4 grid; 35 μm tungsten wires, 150 μm spacing; Innovative Neurophysiology) was implanted into the right aIC (coordinates from Bregma, mm: AP +1.7, ML +3.1, DV −3.5). Electrodes were referenced to a silver ground placed above the right visual cortex and coated with the fluorescent tracer DiI prior to insertion. Implants were secured with Super-Bond cement (Sun Medical). Mice recovered for 4 days post-surgery and were habituated to handling and head stage connection for 3 days.

<u>For combined photoactivation of D1 aIC neurons and single-unit recordings</u>, D1-cre mice received unilateral injection of AAV_9_-hSyn-DIO-hChR2-eYFP (*n* = 2) or a control viral vector AAV_9_-hSyn-DIO-eYFP (*n* = 2) into the right aIC (coordinates from Bregma, mm: AP +1.7, ML +3.1, DV −3.5). 2 weeks after, mice were implanted with an electrode and optic fiber (200 μm diameter, NA >0.90 efficiency, mounted in 1.25 mm ferrules and implanted 50 μm above the aIC injection site). Mice recovered for another 2 weeks before behavioral testing.

### Brain extraction and sectioning

Animals were euthanized with pentobarbital (300 mg/kg, i.p.) and perfused transcardially with Ringer’s solution followed by 4% paraformaldehyde (PFA, Antigenfix, F/P0014, MM France) at 4°C. Brains were post-fixed overnight in 4°C, then cryoprotected in 30% sucrose in PBS 1X at 4°C, and embedded in OCT and sectioned into 50-100 µm coronal or horizontal slices using a freezing sliding microtome (Thermo Scientific, 12062999). Sections were incubated in Hoechst for 10 minutes, washed in PBS 1X twice for 5 minutes each, and mounted with PVA-DABCO mounting medium (Sigma-Aldrich, France). All processed sections, including those containing injection sites, were imaged using a 10x objective.

#### Immunohistochemistry

##### Staining tyrosine hydroxylase

To label dopaminergic neurons in the VTA and SNc, 40 µm coronal slices between Bregma -2.80 mm and -3.52 mm were selected. Sections were permeabilized with 3% Triton-X100 in PBS 1X, then blocked for 1 h at room temperature (RT) in 10% normal goat serum (NGS) with 3% Triton X-100 in PBS 1X. Sections were incubated overnight at 4°C with the mouse anti-Tyrosine Hydroxylase primary antibody (1:2000, Millipore, MAB318) in PBS containing 3% Triton-100x and incubated overnight at 4°C. After three washing steps in 3% Triton X-100 in PBS 1X, sections were incubated for 2 h at RT with the goat anti-mouse Alexa Fluor 647 secondary antibody (1:500, Fisher Scientific A21236), rinsed three times in PBS 1X, incubated in Hoechst (1:20000, Fisher Scientific 11544876) for 10 min, and given a final PBS wash. Sections were then mounted and cover slipped with PVA-DABCO (Sigma Aldrich).

##### Double NeuN and GAD67 staining in section expressing TdTomato

To identify glutamatergic and GABAergic neurons in the insular cortex, NeuN was used to label neurons and GAD67 to label GABAergic somata. Putative glutamatergic neurons were defined as NeuN+ and GAD67-cells, and GABAergic neurons as GAD67+ cells, consistent with cortical composition reported previously^68^. Horizontal sections from D1-cre x Ai14 and D2-cre x Ai14 mice were washed in PBS 1X and blocked for 1h at RT in 10% NGS with 0.5% Triton X-100 in PBS. Primary and secondary antibodies were diluted in 5% NGS with 0.5% Triton X-100 in PBS. Sections were incubated overnight at 4°C with guinea pig anti-NeuN (1:500, Synaptic Systems 266004) and mouse anti-GAD67 (1:500, Millipore MAB5406), washed 3 x 5 min in PBS, then incubated for 3 h at RT with Alexa Fluor 555 goat anti-guinea pig IgG (1:1000, Fisher Scientific A21435) and Alexa Fluor 647 goat anti-mouse IgG (1:1000, Fisher Scientific A21236). After three 5 min PBS washes, Hoechst (1:20000, Fisher Scientific 11544876) was applied for 10 min, sections were washed in PBS, mounted, and cover slipped with PVA-DABCO (Sigma Aldrich).

##### cFos immunohistochemistry

To assess cFOS expression as a marker of neuronal activation, mice received an intraperitoneal injection of vehicle or the D1 receptor agonist SKF38393 (10 mg/kg) and were returned home cage. 90 minutes after, animals were anesthetized with pentobarbital (300 mg/kg, i.p.) and transcardially perfused. Following cryoprotection in 30% sucrose in PBS 1X, brains were sectioned at 50 µm coronal brain sections using a frozen sliding microtome (12062999, Thermo-Scientific). Sections were washed in PBS 1X (3 x 5 min, RT), then blocked for 2 h at RT in 3% NGS in PBS 1X with 0.3% Triton X-100. Sections were incubated overnight at 4 °C with the rabbit polyclonal anti-cFos primary antibody (1:500, Synaptic Systems 226-003), washed in PBS 1X (3 x 10 min), and incubated for 2 h at RT with Alexa Fluor 555 goat anti-rabbit IgG secondary antibody (1:1000, Fisher Scientific 1890860). After PBS washes (3 x 10 min), sections were incubated in Hoechst (1:20000, Fisher Scientific 11544876) for 10 min, washed once in PBS, mounted, and cover slipped with PVA-DABCO (Sigma Aldrich).

### Fluorescent *in situ* hybridization (FISH)

Staining for *Drd1* and *Drd2* mRNAs was performed using single molecule FISH (smFISH). Brains from four C57Bl/6J 8-week-old male mice were rapidly extracted and snap-frozen on dry ice and stored at −80 °C until use. Anterior and posterior insular cortex coronal sections (16 μm) were collected directly onto Superfrost Plus slides (Fisherbrand). RNAscope Fluorescent Multiplex labeling kit (ACDBio; Cat# 323110) was used to perform the smFISH assay according to the manufacturer’s recommendations. Probes used for staining are mm-DrD1+C2 (ACDBio, Cat# 406491-C2), mm-DrD1+C1 (ACDBio, Cat# 461901), mm-DrD2+C3 (ACDBio, Cat# 406501-C3), mm-Slc17a7-C1 (ACDBio; Cat# 416631) and mm-Slc32a1-C2 (ACDBio, Cat# 319191-C2). After incubation with fluorescent-labeled probes, slides were counterstained with DAPI and mounted with ProLong Diamond Antifade mounting medium.

#### Confocal imaging and data analysis

Confocal images were acquired on a Leica SP8 confocal microscope (Leica Microsystems) using a 10x dry objective (NA 0.70) and a 40x oil-immersed objective (NA 1.30).

##### Retrograde mapping and tyrosine hydroxylase staining

Images were acquired with a Leica SP8 confocal microscope (Leica Microscopy) and analyzed in ImageJ. To quantify dopaminergic cell bodies in the VTA and SNc projecting to the insula, images were collected at 40x magnification (field size: 276,79 µm², resolution: 1024x1024 pixels, z-step of ∼2-4 µm). Regions of interest were defined using the Allen Brain Atlas, with brain sections spanning Bregma -2.80 to -3.52 mm. Cell bodies double-labeled with tracer (CTB488 or CTB555) and tyrosine hydroxylase were classified as dopaminergic neurons.

##### Anterograde mapping of dopaminergic outputs

To quantify axonal densities from VTA (ipsi- and contralateral merged) and SNc neurons targeting the insula, eYFP- or mCherry-labeled terminals were imaged bilaterally in the aIC and pIC of coronal sections. Cortical layers were outlined as ROIs, and terminal density was measured per layer, using custom FIJI macros. For each structure, DA terminal signal was normalized to the layer with the highest fluorescence.

##### Double immunofluorescence for NeuN and GAD67

Confocal images were acquired on a Leica SP8 confocal microscope (Leica Microsystems) using a 10x dry objective (NA 0.70) and a 40x oil-immersed objective (NA 1.30).

##### cFos immunohistochemistry and imaging

Z-stacks of regions of interest (ROIs) were acquired with a z-step of ∼2–4 µm at 1024 × 1024 pixels using a 40x objective, and counts were performed on maximum-intensity projections. ROIs were the right anterior insula (aIC, +1.70 mm), right somatosensory cortex (S1, +1.70 mm), right posterior insula (pIC, -0.35 mm), right nucleus accumbens (NAc, +1.21 mm), and the right basolateral amygdala (BLA, +1.31 mm).

##### Quantification of *Drd1* and *Drd2* mRNAs in the insula (FISH)

A cell was classified as positive for *Drd1* and *Drd2* only if it contained at least 5 puncta for each transcript independently.

### Behavioral assays

The temperature and humidity of the experiment room were controlled at 20 to 24°C and 45% to 65%, respectively. Animals were handled at least once a day, and habituated to being connected one week before fiber-photometry, *in vivo* electrophysiology and optogenetic experiments. All tests were performed during the dark phase (active phase) of the reversed light/dark cycle. Behavioral mazes were cleaned with acetic acid 2% and distilled water before each animal session. One camera on top of the experimental apparatus was used for all tests to track the animal’s position and then synchronized with related activities.

<u>Anxiety assays</u>: two recognized tests with predictive validity were used: the elevated plus maze (EPM, Lister, 1987) and the open field test (OFT, Prut & Belzung, 2003). <u>Elevated plus maze</u>: The arena has a plus shape (75 x 75 cm) at 60 cm height, consisting of two open arms (exposed spaces) and two closed arms (protected areas) of 5 cm width. Each animal was placed at the center and facing one open arm and left to explore it for 15 minutes. <u>Open field test</u>: the arena is squared (60 x 60 cm) and the exposed space is the center, while the protected spaces are the borders of the field. The center is defined as 50% of the total open-field arena. Each animal was put in the corner of the open field and left to explore it for 20 minutes.

#### Fiber-photometry recordings

<u>Acquisition</u>: dLight1.1 and gDA3m signals were recorded under low-power blue excitation (470 nm LED, 30-40 µW measured at the patch-cord tip) delivered through a 0.29 NA patch cords. Emitted green fluorescence was collected through the same cord, relayed through a 20x objective, and imaged onto a CMOS sensor after band-pass filtering. An isosbestic 405 nm channel was acquired in parallel to remove motion and non-dopamine-related fluctuations artifacts since it is independent of DA concentration. Photometry and video were sampled at 20 Hz.

<u>Data analysis</u>: photometry signals were processed and analyzed using custom-written scripts in Matlab R2024b, see^2^. The 405 nm reference was fitted to the raw trace by polynomial fit, and normalized fluorescence was computed as dF/F. For z-scoring, the baseline was defined as the dF/F trace with suprathreshold peaks filtered out, as in^71^. Suprathreshold peaks used for baseline exclusion were identified on the z-scored trace with a minimum peak prominence of 2 z and a minimum peak amplitude of 2.58 z. All samples spanning each detected peak width were excluded. The baseline mean and standard deviation were then computed on this baseline for the z-score. Detection of dopamine transients were detected on the dF/F trace and defined as high-amplitude events (amplitude 1.5 median absolute deviation (MAD) above the dF/F). For each transient, peak time and amplitude were extracted, and averaged peak amplitude and frequency were compared across groups.

<u>For entry-based analyses of dopamine signals</u> (Fig. 1m-n and Extended Data Fig. 2h-i), photometry traces were expressed as z-scored dF/F as described above. For each open- or closed-arm entry event, mean activity was quantified in a pre-entry center (CT) window (-3s to - 1s relative to entry) and in an arm entry window (0s to 2 s), and these values were used for statistical analyses.

#### *In vivo* pharmacology

<u>For intraperitoneal (i.p.) injection of the selective D1 agonist SKF38393</u>, mice were habituated for 2 days to the i.p. injections. On the test day, mice received either sterile saline (0.9% NaCl) or the D1 agonist SKF38393 (10 mg/kg, diluted with saline). 30 minutes after, mice were placed in the EPM for 15 minutes.

<u>For bilateral intracranial infusion of the D1 agonist SKF38393, or D1 antagonist SCH23390</u>, mice were handled for 5 days to habituate them to restraint and cannula connection. Infusions were performed using 33GA stainless steel internal canulae (5 mm long, Phymep) projecting 1 mm beyond 26 GA guide cannula, connected via tubing to Hamilton syringes mounted on a micro-infusion pump (KDS Legato 101, Phymep). Internal cannula tip coordinates relative to Bregma (in mm) were AP +1.7, ML ± 3.1, DV -3.5. SKF38393 hydrobromide (0.25 mg/mL, Tocris) or SCH23390 hydrochloride (0.5 mg/ml, Tocris) was diluted in sterile saline (0.9% NaCl). To verify injection sites, drugs were mixed with 0.5% CTB-AF555 (0.4%, Fisher Scientific). Mice received 250 nL per hemisphere of SKF38393, SCH23390 or vehicle at a rate of 1 µl.min^-1^. Internal cannulae were withdrawn 1 minute after infusion, and mice were placed in the center of the EPM 5 min later, for 15 minutes.

#### Optogenetics

<u>For combined optogenetic manipulations and fiber-photometry recordings of dopamine signals</u>, unilateral stimulations of VTA dopaminergic neurons were delivered by coupling the implanted optical fiber to a patch cord connected to a 635 nm laser source (Intelligent Optogenetics Laser System, RWD; IOS-635 nm). In rsChRmine-oScarlet and mCherry control mice, light output was calibrated to 10 mW at the fiber tip and delivered in 36 epochs consisting of 2 s ON trains (20 Hz, 10 ms pulses; 40 pulses per epoch) followed by 3 s OFF. Photometry signals were expressed as z-scored dF/F, using the -3s to -1s pre-stimulation period as baseline. Mean z-scored dopamine signal in the baseline window (-3s to -1s) and the stimulation window (0s to 2 s) was extracted for statistical analyses.

<u>For optogenetic activation of D1 aIC neurons in the EPM</u>, bilateral stimulations of D1 aIC neurons were achieved by connecting fiber implants to two patch cords connected to a 465 nm laser source (Intelligent Optogenetics Laser System, RWD, IOS-465 nm). ChR2 and eYFP mice received pulse trains (465 nm, 20 mW at the tip of the fiber, 5 ms pulse duration, 20 Hz repetition rate for 3 minutes).

<u>For combined photoactivation of D1 aIC neurons and single-unit recordings</u>, *in vivo* recordings and optogenetic stimulations were synchronized. Unilateral stimulations of D1 aIC neurons were achieved by connecting the fiber implant to a patch cord connected to the 465 nm laser source (Intelligent Optogenetics Laser System, RWD, IOS-465 nm). ChR2 and eYFP mice received pulse trains (465 nm, 20 mW at the tip of the fiber, 5 ms pulse duration, 20 Hz repetition rate for 3 minutes). The entire recording session was 15 minutes. The first 3 minutes served to record baseline firing rate.

#### *In vivo* electrophysiology

##### Single-unit recordings

16-channel electrodes were connected to an 18-pin nanoconnector (Omnetics A79014-001) and a unity-gain 16-channel headstage (Intan RHD2132). Signals were acquired via an Open Ephys board at 30 kHz, hardware high-pass 300 Hz and and low-pass 6 kHz, synchronized to behavior.

##### Spike sorting

Spike detection and sorting used Kilosort 4^72^ and custom-written python scripts. Common reference removal and per-channel normalization were applied before template matching. Principal component analysis was applied per channel. Units were accepted as single-units if they formed an isolated cluster in PC space, had a refractory period violation consistent with a refractory period longer than 1 ms and passed visual inspection of waveforms and auto-correlograms. Multi-units that failed these criteria were excluded.

##### Behavioral tracking

Mouse pose was tracked using markerless pose estimation with DeepLabCut 2.3.10 (single animal mode for pose estimation^30^). Around 5-20 frames per video were manually annotated for: nose, ears, neck, three backbone points, and three tail points, and a ResNet-50 model was trained over 80% (100 000 iterations) of the manually annotated frames. To account for head dips, the effective open arm border was widened by a tolerance equal to the average snout-to-shoulder offset. Time points with coordinates outside the EPM were removed. Outliers were defined as any body part at a particular timepoint *t* larger than 5 cm between consecutive frames; outliers were replaced by linear interpolation over *t*−10 to *t*+10 frames. Head dips closer than 1 s to a previous head dip were merged and counted once.

<u>Individual behaviors were analyzed during the elevated-plus maze data using DeepOF</u> version 0.8.1^31,73^ for supervised behavioral analysis of nine behaviors including: head dips, climbing in closed arms (climbing-arena in DeepOF module), arena sniffing, immobility, stationary look around, stationary active, stationary passive, moving, instantaneous speed (description of behaviors can be found here: https://deepof.readthedocs.io/). In addition, the instantaneous speed (speed-p95) was computed by calculating the Euclidean displacement of the Center key point between consecutive frames divided by the frame interval (frames per second specific), which was then defined as the empirical 95th percentile of these per-frame speeds, providing a robust summary of high-end movement bursts. All behaviors were analyzed throughout the different zones, in which a separation was made based on the center key point of the animal in the open arm, closed arm and center zone. Data were analyzed for the total 15 min.

##### Event definition, binning and trial snippets

Behavioral events were defined as closed arm (CA) entry, open arm (OA) entry and head dip (HD) over the borders of the open arms of the EPM, and all behavioral entries into the closed or open arm take the center of the EPM as the origin. To avoid any false positive, mice had to stay at least 2 seconds in the center of the EPM for entry events and 1 second in the open arms for the head dips. For each event, a 6.5 s snippet was extracted and centered on the event time. Spikes were binned at 25 ms, then converted to a sliding 250 ms window by summing over 10 successive 25 ms bins with a 25 ms step. Population matrices were stored as (units, time) shape.

##### Single-unit response classification

For each event type, per-unit peri-event time histograms were z-scored using a baseline of -3 to -1 s relative to event onset. Within the post-event window (0, +3 s), responses were classified using a sliding window of 200 ms (4 consecutive 50 ms bins after the 50 ms resampling step): units were considered as excited when all samples in any 200 ms window exceed z = 1.65, inhibited when all samples in any 200 ms window are below z = -1.65 and neutral for neither criterion met.

##### Single-unit response classification following photoactivation of D1 aIC neurons

To test whether units were expressing ChR2, we applied a Wilcoxon signed-rank test with a significance threshold of \*\**p*<0.01 to compare the firing of each unit during a baseline window (1 s before light onset) to the firing in an experimental window of 10 ms, starting at light onset. If the average z-score during the experimental window was positive, and the Wilcoxon signed rank test was significant (\*\**p*<0.01), the unit was classified as excited, whereas, if the average z-score during the experimental window was negative and the Wilcoxon signed-rank test was significant (\*\**p*<0.01), the unit was classified as inhibited.

##### Analysis of unit class proportions

To compare the proportion of unit response categories between Vehicle and SKF groups, units were classified into four mutually exclusive categories (opposite, same, selective, neutral) based on their activity across pairs of behaviors: CA entry and OA entry (Fig. 4h), or OA entry and head dip, or CA entry and head dip (Extended Data Fig. 4j-k). Group differences in class composition were assessed using a permutational multivariate analysis of variance. Post hoc comparisons were performed using Welch’s t-tests.

##### Population matrix construction and principal component analysis

For each mouse and trial type, z-scored, trial-averaged population vectors were concatenated along time to build trial-type blocks. Blocks for open arm entry, closed arm entry, and head dips were concatenated along time to form a global matrix across units, then concatenated across mice along the unit axis to obtain a single analysis matrix. Principal component analysis (PCA) was performed on the covariance of this matrix. Trajectories were obtained by projecting time points onto the principal components. <u>Leave-one-mouse-out projection and trajectory metrics</u>. To estimate across-animal generalization, we used a leave-one-mouse-out procedure. For each mouse, we removed its units from the eigenvector matrix and re-ordered eigenvectors by the remaining eigenvalues, then projected that mouse concatenated snippets into the PCA space derived from the other mice. Trajectory length per condition was computed as the sum of step sizes across the first 3 principal components after light Gaussian smoothing. We report on the scalar trajectory length. As a control, a shuffled dataset was generated by randomizing spikes per unit and bin using a binomial process matched to firing statistics. These data were binned and processed identically as the real data.

##### Decoding

To quantify how well aIC population activity discriminated against behavioral conditions, we trained linear support vector machine (SVM) classifiers on single-trial instantaneous population vectors. For each neuron, spike counts were computed in 250 ms bins before and after the onset of behavioral events (closed arm entry, open arm entry, and head dips), yielding one population vector per trial and time bin (units and trials pooled across mice). Vectors were z-scored across trials, and for each decoding analysis (for example, closed arm entry versus open arm entry, or a given event versus the rest of the recording), we used the same number of trial type events between trials by subsampling each condition to the global minimum trial count. Data were then randomly split into training (60%) and testing (40%) sets. SVM model training was performed on the training set using C-type classification with a linear kernel and fivefold cross-validation. The trained model was subsequently evaluated on the held-out testing set, and decoding accuracy was defined as the proportion of correctly classified trials. This procedure was repeated 10 times with new random train/test splits to obtain a distribution of accuracies for statistical comparison. To assess significance, we used a two-sided permutation test: for each decoding comparison, we computed the difference in mean decoding accuracy between condition (\**p*<0.05). Analogous decoding analyses were performed with a k-Nearest Neighbors (kNN) classifier to obtain similar accuracies.

##### Non-linear dimensionality reduction (CEBRA)

To characterize nonlinear population dynamics, we used CEBRA-Time (Consistent EmBeddings of high-dimensional Recordings using auxiliary^33^), a self-supervised contrastive learning algorithm. CEBRA-Time models were trained on 50 ms spike-count vectors using a multi-session (multi animals) setup that learns embeddings invariant across animals, based solely on single-unit spiking and without behavioral labels. CEBRA-Time employs a nonlinear encoder (deep neural network) to map high-dimensional neural activity into a low-dimensional latent space. Through contrastive learning, the model pulls together temporally neighboring activity patterns while pushing apart dissimilar activity patterns, thereby capturing the intrinsic temporal structure of neural population dynamics. For each analysis, models were trained on 60% of the data, and training loss was monitored to ensure how good our model can predict values based on 60% of the data. The remaining 40% of the data were projected into the three-dimensional latent space in an embedding (50 ms bins). Behavioral labels were then overlaid post hoc and were not used during training. Decoding analyses were then performed on the projected latent clusters. We applied CEBRA-Time in two complementary ways: (1) on the behavioral snippets (discrete: closed arm entry, open arm entry, head dips), and (2) on the entire EPM recording (continuous). Both analyses used identical architecture and hyperparameters (offset5 architecture, learning rate 3.10^-4^, 20 hidden units, temperature 1). The number of training iterations was adjusted according to dataset size: 4 000 iterations for snippet analysis and 40 000 for the entire recording analysis. To quantify behavioral information in the latent space, we performed decoding analysis using a support vector machine (SVM), with an analogous k-nearest neighbors (kNN) classifier for comparison. Decoding was performed on neural activity projected onto the first three latent dimensions to discriminate behavioral states (closed arm entry vs open arm entry vs head dips; or closed arm vs open arm position). Two complementary metrics were computed from the latent embeddings:

(1) the <u>Shannon entropy</u>, computed from the empirical distribution of embedded points within each condition, to quantify the dispersion of the population state. The Shannon entropy *H* is defined as:

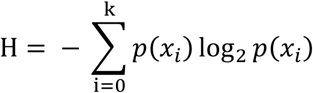

where 𝑝(𝑥_𝑖_) represents the probability of observing point 𝑥_𝑖_in the latent space. Higher entropy reflects greater dispersion and variability of the embedded population activity, whereas lower entropy indicates more compact and consistent/reliable representations.

(2) the <u>barycenter distance</u>, defined as the Euclidean distance between the centroid of two latent clusters. This metric quantified the separation between neural representations associated with different behavioral states and serves as an estimate of neural divergence between conditions.

#### Histological verification of photometry, pharmacology, optogenetic and electrophysiology targets

At the end of behavioral experiments, mice were anesthetized and transcardially perfused with PBS followed by 4% PFA. Brains were post-fixed in 4% PFA and cryoprotected in 30% sucrose in PBS before sectioning. Coronal sections (100 µm) of the insular cortex were collected and mounted for fluorescence imaging. Sections were imaged on a Leica fluorescence microscope to verify viral expression and the location of optic fiber, cannula, or electrode placements using the Paxinos Brain Atlas. For each animal, histological images were overlaid onto atlas templates to reconstruct maps of viral infusion and implant locations. Representative examples and overlaid histological verifications are shown in the corresponding Extended Data Figures for photometry implants (Extended Data Fig. 1h and Extended Data Fig. 2g), pharmacology injection sites (Extended Data Fig. 3a), optogenetic manipulations (Extended Data Fig. 3e-f) and *in vivo* electrophysiology recordings (Extended Data Fig. 4a).

## Supporting information

Statistical_Table

## Code availability

Custom-written codes used to analyze fiber photometry data and *in vivo* electrophysiology data are available on the GitHub page of the Beyeler Lab: https://github.com/beyelerlab/FiberPhotometry_anxiety https://github.com/beyelerlab/in_vivo_ephys.

## Statistical analysis

For statistical analyses, GraphPad Prism 10 (version 10.6.1) was used. Normality and homogenous variance of the data were assessed by Shapiro-Wilk and F-tests, respectively. Paired or unpaired two-tailed *t*-tests were used for comparisons between two conditions. For analyses involving more than two groups and/or multiple factors, one-way or two-way ANOVA was performed followed by post hoc multiple-comparisons tests. For correlation analyses, Pearson’s correlation coefficients (R^2^) and linear regression were calculated. Grubb’s tests were performed for each test to identify potential outliers. Significance was set at \**p*<0.05. A table of all statistical analyses is reported in Table 2.

## Acknowledgments

We want to acknowledge the past and present members from the Beyeler Lab for discussions and advice, especially Débora Jacky and Anes Ju. We thank Sara Laumond, Julie Tessaire, and the technical staff of the Neurocentre Magendie animal facility, especially Delphine Gonzales, Elizabeth Huc, Mégane Marion, and Jean-Baptiste Bernard, for their invaluable support. We thank Cyril Herry for the DAT-cre mouse line. We are grateful to Yulong Li for providing the cre-dependent gDA3m plasmid. We thank the Bordeaux Imaging Center (BIC) for using their confocal microscope, and especially Jérémie Teillon for advice. We also thank Adeline Cathala and Luigi Bellocchio for their guidance and assistance with pharmacological experiments. Finally, we thank Sébastien Delcasso and Stéphane Valerio from Aquineuro for their support regarding *in vivo* electrophysiology experiments.

## Authorship contributions

YC and AB conceptualized the study. AB provided resources to perform all experiments except the FISH staining, supported by EV. YC contributed to the analysis of all experiments.

*Dopaminergic architecture of the insula*. MA and GV conducted D1+ neurons and D2+ neurons mapping of the insula, respectively. EV performed FISH experiments, and DR analyzed the data. *Dopamine signal recordings*. YC and GV performed and analyzed dLight1.1 / gDA3m recordings. *Pharmacology*. YC, GV, DR, CN, and TH conducted pharmacological experiments.

*AI-based behavioral phenotyping.* JB performed the deepOF analysis.

*Single-unit recordings and analysis*. YC performed *in vivo* single-unit recordings while TDR, AG, and DR assisted in analyzing these data. TDR, AG and YC wrote computational analysis scripts for the study, with help from YKW and JG.

*Manuscript*. YC and AB wrote the original draft of the manuscript. YC made all the figures. All authors critically reviewed and approved the final version before submission.

## Supplementary Figures and Legends

**Extended Data Fig. 1:**
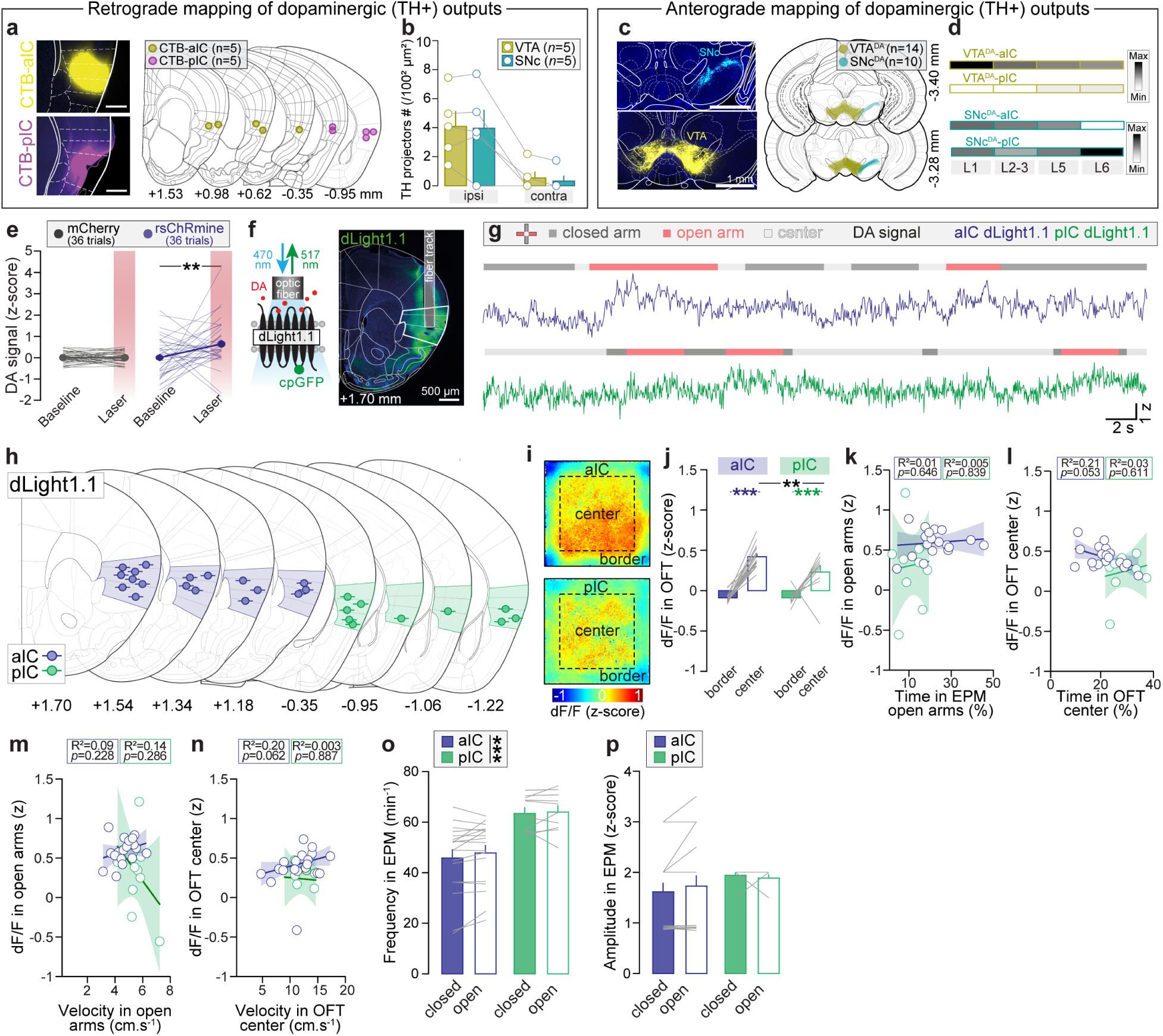
Dopaminergic system architecture in the insular cortex. a-c, Histological verification of retrograde (**a**) and anterograde (**c**) mapping of dopaminergic outputs to the aIC and pIC. **b,** Total number of TH projectors from the VTA and SNc in the ipsilateral and contralateral side (Two-way ANOVA: Region: F_1,16_=0.04, *p=*0.8454, Main effect Hemisphere: F_1,16_=17.62, \*\*\**p=*0.0007, with no interaction: F_1,16_=0.002, *p=*0.9617). **d,** Heatmaps showing the fluorescent intensity of VTA (top) and SNc (bottom) dopaminergic terminals in the aIC and pIC cortical layers. Gray scale illustrates the magnitude of the fluorescence intensity (black is the highest intensity). **e,** VTA^DA^ cell body optogenetic stimulations in rsChRmine mice (20 Hz, 10 mW, 2s, 10 ms pulses) increase DA signal (\*\**p=*0.0062), relative to mCherry (Group: F_1,356_=2.33, *p=*0.1278, Main effect Period: F_1,356_=4.22, \**p=*0.0407, with a Group x Period interaction: F_1,356_=4.41, \**p=*0.0365). **f,** dLight1.1 sensor schematic and representative confocal image of dLight1.1 expression in the aIC. **g,** Representative dopamine signal (dLight1.1) traces from aIC (top) and pIC (bottom) in the EPM. **h,** Histological verification of dLight1.1 viral injections into aIC and pIC. **i-j,** Heatmaps (**i**) and quantification (**j**) of averaged z-scored DA signal in aIC and pIC during the OFT showed that DA signals are increased in center vs. borders in aIC (\*\*\**p<*0.0001) and pIC (\*\*\**p<*0.0001), with a stronger signal in aIC (\*\**p*=0.0065) relative to pIC (Main effect Zone: F_1,52_=120, \*\*\**p<*0.0001, Main effect Region: F_1,52_=5.82, \**p=*0.0194, with a Zone x Region interaction: F_1,52_=5.88, \**p=*0.0189). **k-l,** DA signals in the aIC and pIC do not correlate with the time mice spent in open arms (**k,** two-tailed Pearson’s correlation, aIC: R²=0.0135, *p=*0.6462; pIC: R²=0.0055, *p=*0.8391) or OFT center (**l,** aIC: R²=0.2151, *p=*0.0525; pIC: R²=0.0338, *p=*0.6109), except for the amplitude of DA signal in OFT center that tends to negatively correlate with the time mice spent in OFT center (**l**). **m-n,** Z-scored DA signals in aIC and pIC did not correlate with the velocity in EPM open arms (two-tailed Pearson’s correlation, aIC: R²=0.09, *p=*0.228; pIC: R²=0.20, *p=*0.286) or OFT center (aIC: R²=0.20, *p=*0.062; pIC: R²=0.003, *p=*0.887). **o,** aIC and pIC dopamine transient frequency and amplitude in the EPM (Frequency: Main effect Region: F_1,52_=27.5, \*\*\**p*<0.0001, Zone: F_1,52_=0.15, *p=*0.7003, with no interaction; Amplitude: Region: F_1,52_=1.77, *p=*0.1886, F_1,52_=0.0305, *p=*0.8620, with no interaction). **a-b:** aIC and pIC (n=5 mice). **c-d:** VTA (n=14 mice) and SNc (n=10 mice). **e:** rsChRmine (n=3 mice) and mCherry (n=2 mice). **f-o:** aIC (n=18 mice) and pIC (n=10 mice). **b, e, j, o:** Two-way ANOVA and Tukey’s post-hoc tests were performed. Data are: mean ± SEM.

**Extended Data Fig. 2:**
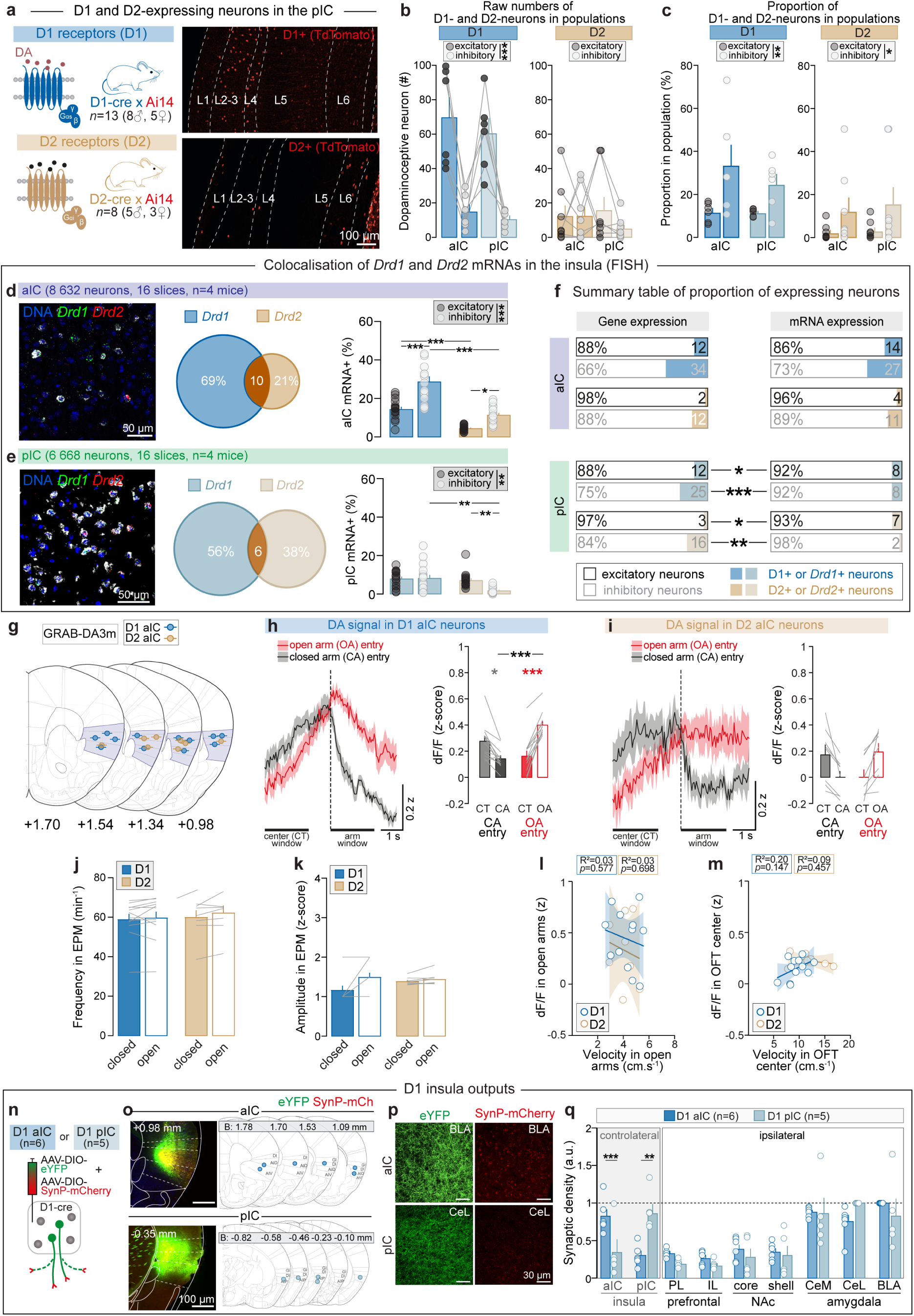
Role of D1+ and D2+ anterior insula neurons in anxiety-related behaviors. **a,** Confocal images of tdTomato-expressing neurons in D1 (top) and D2 (bottom) subpopulations of the pIC among cortical layers. **b,** Absolute numbers of D1+ neurons (Region: F_1,20_=0.7961, *p=*0.3829, Main effect Subpopulation: F_1,20_=46.06, \*\*\**p<*0.0001, with no Region x Subpopulation interaction: F_1,20_=0.1018, *p=*0.7530) and D2+ neurons in excitatory and inhibitory subpopulations in the insula (Region: F_1,28_=0.1208, *p=*0.7308, Subpopulation: F_1,28_=0.8323, *p=*0.3694, with no Region x Subpopulation interaction: F_1,28_=0.8724, *p=*0.3583). **c,** Proportion of D1+ neurons (Region: F_1,20_=0.711, *p=*0.409, Main effect Subpopulation: F_1,20_=10.8, \*\**p=*0.0037, with no Region x Subpopulation interaction: F_1,20_=0.659, *p=*0.4266) and D2+ neurons within excitatory and inhibitory subpopulations in aIC and pIC (Region: F_1,28_=0.169, *p=*0.6844, Main effect Subpopulation: F_1,28_=4.99, \**p=*0.0337, with no Region x Subpopulation interaction: F_1,28_=0.066, *p=*0.7987). **d-e,** Colocalization and proportion of *Drd1* and *Drd2* mRNAs among excitatory (*Slc17a7*+) and inhibitory (*Slc32a1*+) neurons in the aIC (**d,** Main effect Receptor: F_1,60_=42.6, \*\*\**p<*0.0001, Main effect Subpopulation: F_1,60_=69.8, \*\*\**p<*0.0001, with a Receptor x Subpopulation interaction: F_1,60_=4.89, \**p=*0.0308), and pIC (**e,** Main effect Receptor: F_1,60_=4.70, \**p=*0.0341, Main effect Subpopulation: F_1,60_=9.76, \*\**p=*0.0027, with Receptor x Subpopulation interaction: F_1,60_=5.9, \**p=*0.0182). **f,** Summary diagram of the proportion of D1 and D2 neurons expression in the insula (two-tailed unpaired *t*-tests, *cf.* Table 2 for statistical values). **g,** Histological verification of gDA3m viral injections into aIC and pIC. **h,** Peri-event z-scored DA signal onto D1 aIC neurons aligned to closed or open arm entries showing a decrease of DA signals at CA entry (\**p=*0.0108) and increased at OA entry (\*\*\**p=*0.0005), with a stronger increase (\*\*\**p*<0.0001) at OA entry compared to CA entry (Space: F_1,68_=2.48, *p=*0.1221, Main effect Event: F_1,68_=4.56, \**p=*0.0384, with a Space x Event interaction: F_1,68_=31.1, \*\*\**p*<0.0001). **i,** No difference of DA signals was found onto D2 aIC neurons (Space: F_1,36_=0.0243, *p=*0.8772, Event: F_1,36_=0.0498, *p=*0.8251, with Space x Event interaction: F_1,36_=8.74, \*\**p=*0.0063). **j,** Dopamine transient frequency onto D1 and D2 aIC neurons in the EPM (Region: F_1,36_=0.306, *p=*0.5836, Zone: F_1, 36_=0.22, *p=*0.6418, with no interaction). **k,** Dopamine transient amplitude onto D1 and D2 aIC neurons in the OFT (Region: F_1,36_=0.0342, *p=*0.8544, Main effect Zone: F_1, 36_=4.3, \**p=*0.0453, with no interaction). **l-m,** Z-scored DA signal onto D1 or D2 aIC did not correlate with the velocity in EPM open arms (two-tailed Pearson’s correlation, aIC: R²=0.03, *p=*0.577; pIC: R²=0.03, *p=*0.698) or OFT center (aIC: R²=0.20, *p=*0.147; pIC: R²=0.09, *p=*0.457). **n,** Viral strategy to map D1 aIC and pIC outputs using a mix of two viral vectors to label axons and synaptic contacts in D1-cre mice. **o,** Histological verification of viral injections into the aIC and pIC of D1-cre mice. **p,** Representative confocal images of axonal (green) and synaptic (red) densities in the BLA and CeL arising from D1 aIC (*top*) and D1 pIC (*bottom*) neurons. **q,** Relative synaptic density in different target regions from D1 aIC and pIC outputs (Main effect Outputs: F_11,107_=18.5, \*\*\**p*=0.0001, Insula section: F_1,107_=0.0174, *p=*0.8954, with a Outputs x Insula section interaction: F_11,107_=3.82, \*\*\**p*<0.0001). **a**: D1 (n=13 mice) and D2 (n=8 mice). **b-c:** D1 (n=6 mice) and D2 (n=6 mice). **d-f:** aIC (n=4 mice) and pIC (n=4 mice). **g-m:** D1 aIC (n=12 mice) and D2 aIC (n=8 mice). **n-q:** D1 aIC (n=6 mice) and D1 pIC (n=5 mice). **b-e, h-k, q:** Two-way ANOVA and Tukey’s post hoc tests. Data are: mean ± SEM.

**Extended Data Fig. 3:**
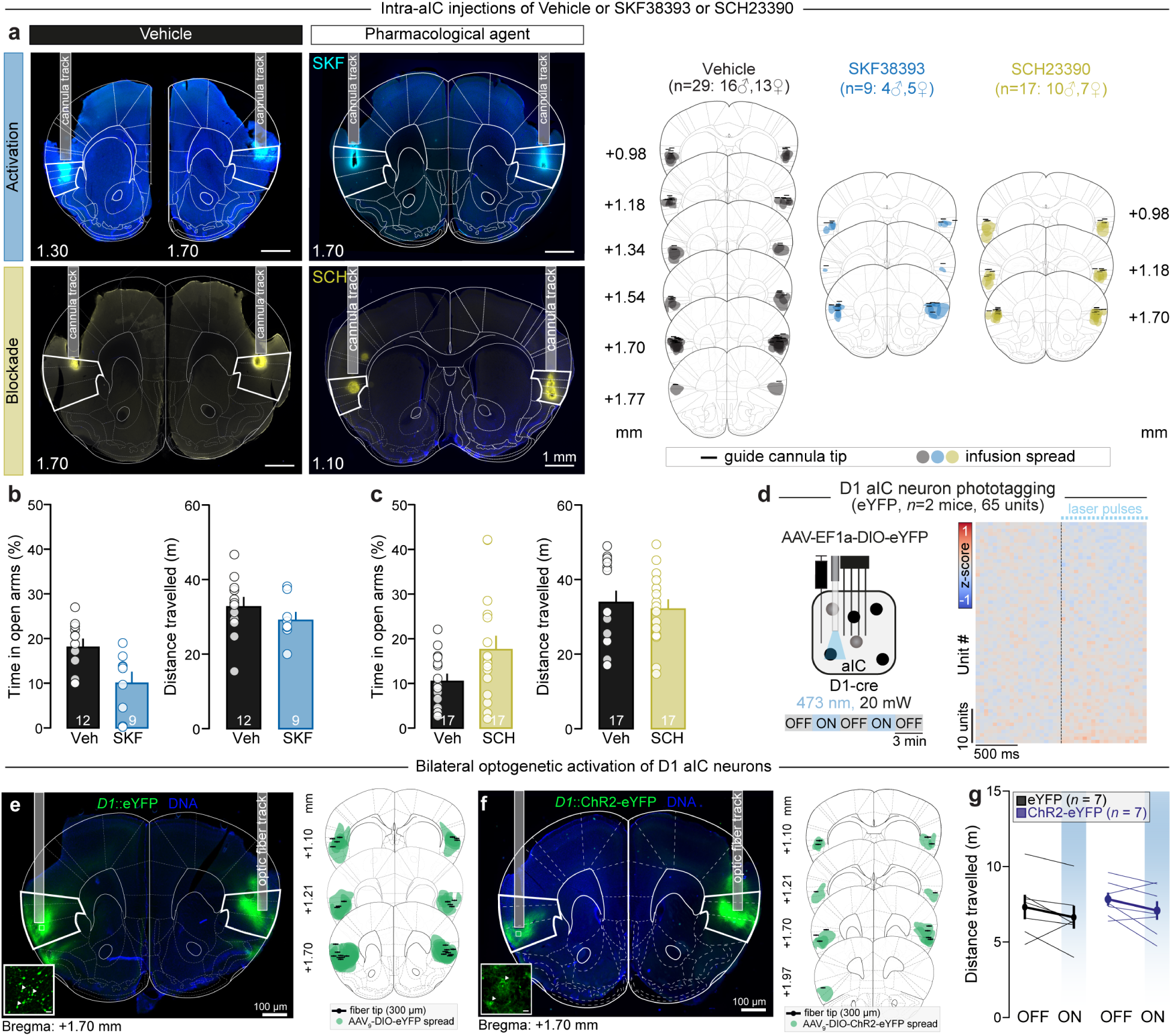
Additional analyses of D1 manipulations in the anterior insula. **a,** Representative images illustrating the location of bilateral cannulae implants and the infusion of the vehicle, the D1 agonist SKF38393 and the D1 antagonist SCH23390 in the aIC. Local infusions were mixed with a fluorescent marker (CTB). Antero-posterior levels (distance from Bregma, in millimeters) are indicated next to each slice. **b-c,** Raw time spent in open arms and total distance travelled in the EPM for SKF- (**b**) and SCH-treated (**c**) mice related to the Main Fig. 3k-m. **d,** Photoactivation of D1 neurons expressing the fluorescent protein eYFP in the aIC (n=2 mice, 65 units) showing no effect of laser stimulations on population activity. **e-f,** Histological verification of viral injections and fiber implants for eYFP (**e**) and ChR2 (**f**) mice. **g,** No change of total distance travelled in eYFP and ChR2 mice during the light stimulation (Two-way ANOVA, Group: F_1,24_=0.395, *p=*0.5357, Light mode: F_1,24_=1.27, *p=*0.27, with no Group x Light mode interaction: F_1,24_=0.0023, *p=*0.9569).

**Extended Data Fig. 4:**
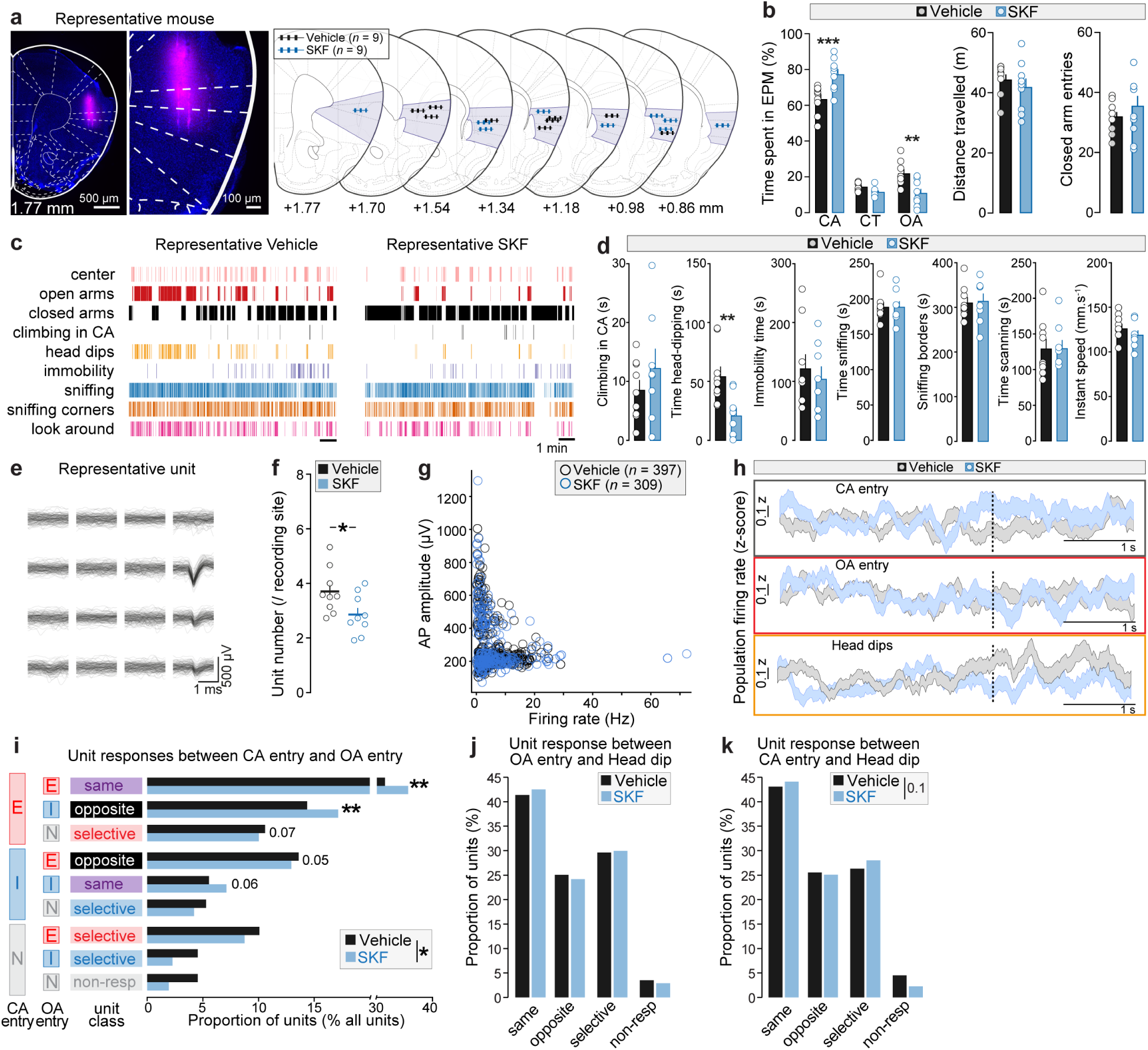
Additional analyses of single-unit anterior insula activity during anxiety-related behaviors. **a,** Representative image of the probe track coated with a red fluorescent dye (DiI) in aIC and histological verification for vehicle and SKF mice. Antero-posterior levels (distance from Bregma, in millimeters) are indicated at the bottom. **b,** SKF injection increased and decreased the time mice spent in the closed arms (CA) and open arms (OA), respectively compared to vehicle-injected mice (two-way ANOVA, Group: F_1,48_=1.2x10^-14^, *p>*0.99, Main effect Location: F_2,48_=467, \*\*\**p<*0.0001, with Group x Location interaction: F_2,48_=18.2, \*\*\**p*<0.0001), without altering the total distance travelled and the number of total entries into the closed arms. **c,** Representative behavioral raster plots from one vehicle and SKF-treated mice in the EPM. **d,** No alteration of solitary behaviors was found in the EPM between vehicle- and SKF-treated mice, except for a decrease of time spent in head-dipping following SKF, relative to vehicle injection (two-tailed unpaired *t*-test, *t*=3.299, \*\**p*=0.0051). **e,** Waveform of a representative unit shown on the 4 x 4 electrode. **f,** Averaged unit number per channel on the electrode (vehicle: 3.71 ± 0.30 units, SKF: 2.86 ± 0.21 units). **g,** Scatterplot representing the action potential amplitude as a function of the baseline firing rate for all units recorded (vehicle: n=397 units, SKF: n=309 units). The averaged firing rate for vehicle-treated mice and SKF-treated mice are: 6.44 ± 0.28 Hz, 6.53 ± 0.38 Hz, respectively. **h,** Peri-event time course of the z-scored population firing rate centered on three behaviors: CA entry (top), OA entry (middle) and head dips (bottom) for vehicle and SKF mice. **i,** Distribution of nine outcome categories analyzed by multinomial logistic regression with non-responding classification as the reference (F=2.363, \**p*=0.05). Among unit classes, D1 systemic activation increased the percentage of ‘saliency-coding’ (\*\**p*=0.0061) and ‘anxiety-coding’ units (\*\**p*=0.0071). **j-k,** Distribution of four outcome categories of unit response to OA entry and head dip (**j**) and between CA entry and head dip (**k**) Vehicle and SKF groups using a permutational multivariate analysis of variance. **a-k:** vehicle (397 units, n=9 mice) and SKF38393 (309 units, n=9 mice). Data are: mean ± SEM.

**Extended Data Fig. 5:**
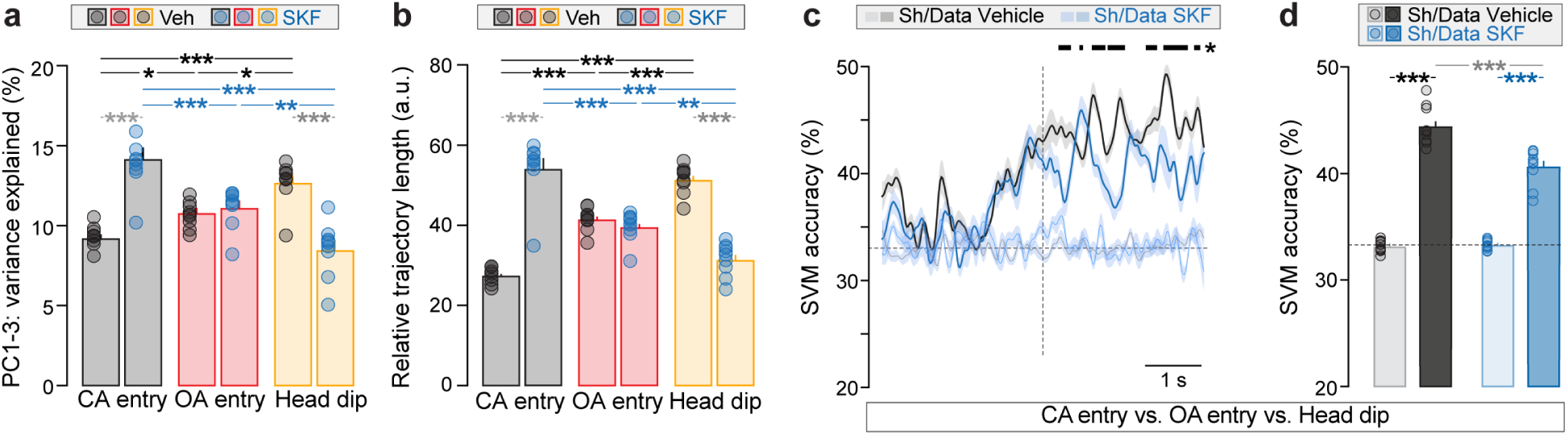
Additional linear analyses on population anterior insula activity during anxiety-related behaviors. **a,** Variance explained by the first three PCs for the three trial types (Event: F_2,48_=0.653, *p=*0.5250, Group: F_1,48_=3.77, *p=*0.0582, with an Event x Group interaction: F_2,48_=54, \*\*\**p*<0.0001) was higher for CA entry (\*\*\**p*<0.0001) and shorter for HD (\*\*\**p*<0.0001) after SKF compared to vehicle. **b,** Relative neural trajectory lengths (Event: F_2,48_=0.198, *p=*0.8214, Group: F_2,48_=1.79, *p=*0.1868, with an Event x Group interaction: F_2,48_=135, \*\*\**p*<0.0001) were longer for CA entry (\*\*\**p*<0.0001) and shorter for HD (\*\*\**p*<0.0001) after SKF compared to vehicle. **c,** A SVM classifier was used to decode the activity between CA entry, OA entry and Head dip for vehicle- and SKF-treated mice (permutation test, \**p*<0.05). **d,** SVM accuracy differentiating CA entry, OA entry and head dip in SKF-treated mice was decreased following D1 activation (\*\*\**p*<0.0001), relative to vehicle (Two-way ANOVA, Main effect Data type: F_1,36_=556, \*\*\**p<*0.0001, Main effect Group: F_1,36_=21.6, \*\*\**p<*0.0001, with an Data type x Group interaction: F_1,36_=24.1, \*\*\**p*<0.0001). The dashed line represents the chance level. **a-d:** vehicle (397 units, n=9 mice) and SKF38393 (309 units, n=9 mice). **a-b, d:** Two-way ANOVA and Tukey’s post hoc tests. Data are: mean ± SEM.

**Extended Data Fig. 6:**
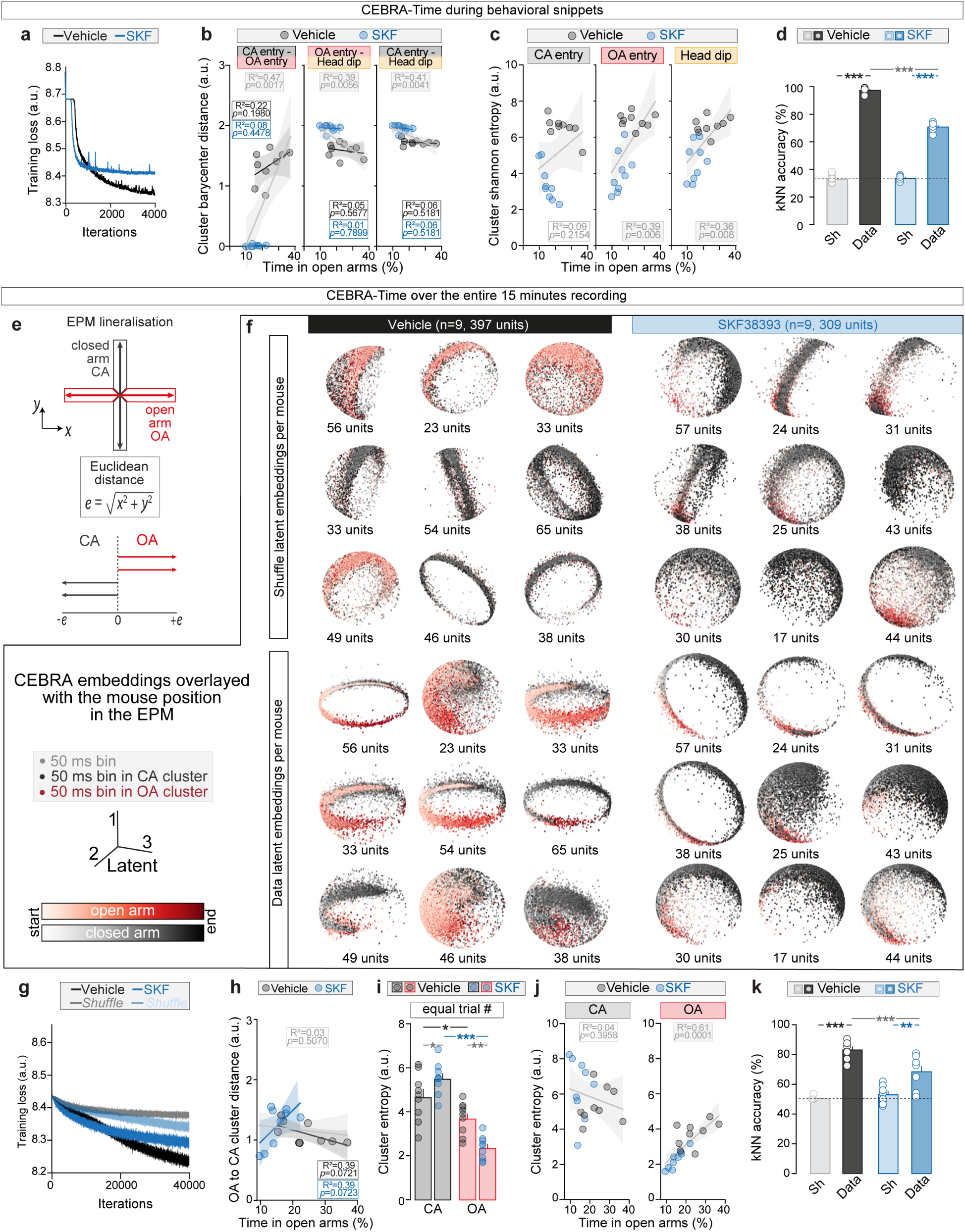
Additional nonlinear analyses on population anterior insula activity in anxiety-related behaviors and spaces. a-d, CEBRA-Time during behavioral snippets. **a,** InfoNCE training loss curve from vehicle and SKF groups (4000 iterations). **b,** The distances between cluster barycenters (vehicle and SKF) associated with different pairs of behaviors for: CA entry-OA entry, OA entry-head dip, and CA entry-head dip, do not correlate with the time spent in the open arms of the EPM, except when the barycenter distances from both vehicle and SKF groups are pooled together (i.e., the shaded gray area representing the 95% confidence interval). **c,** Pooled (vehicle and SKF) correlation analyses of Shannon entropy for CA entry, OA entry, and head dip clusters with time spent in the open arms. Shannon entropy for OA entry and head dip clusters negatively correlates with time spent in the open arms (OA entry: R²=0.39, \*\**p*=0.006; head dip: R²=0.36, \*\**p*=0.008). **d,** k-Nearest Neighbor decoder achieved above 95% accuracy in distinguishing OA entry, CA entry and head dip trials in vehicle conditions (\*\*\**p*<0.0001), while it decreased to 70% (\*\*\**p*<0.0001) following D1 activation (Two-way ANOVA, Main effect Data type: F_1,32_=3746, \*\*\**p<*0.0001, Main effect Group: F_1,32_=247, \*\*\**p<*0.0001, with an Data type x Group interaction: F_1,32_=265, \*\*\**p*<0.0001). **e,** EPM linearization to quantify the mouse position in the EPM. **f,** CEBRA latent embeddings representing population anterior insula activity (shuffled and non-shuffled) binned in 50 ms and projected onto the first three latent spaces, for vehicle and SKF38393 mice. Each embedding represents the neural activity from one mouse with the number of recorded units per mouse at the bottom of each embedding. We post hoc color coded each embedding based on the mouse position in the EPM (black: closed arms, red: open arms). **g,** InfoNCE training loss curve from vehicle and SKF groups (40000 iterations). **h,** Barycenter distance between OA and CA latent clusters do not correlate with the time mice spent in open arms for vehicle and SKF groups (two-tailed Pearson correlations, Vehicle: R²=0.39, *p*=0.0721; SKF: R²=0.39, *p*=0.0723 and pooled data: R²=0.03, *p*=0.5070). **i,** Shannon entropy was computed on the same number of trials in the closed arms and the open arms. Higher entropy for the CA latent cluster (\**p*=0.0462) but shorter for the OA latent cluster (\**p*=0.0133) after SKF compared to vehicle (two-way ANOVA, Arm: F_1,16_=60.3, \*\*\**p*<0.0001, Group: F_1,16_=0.152, *p*=0.7015, with an Arm x Group interaction: F_1,16_=10.8, \*\**p*=0.0046). **j,** Shannon entropy for the CA latent cluster does not correlate with time spent in the open arms when vehicle and SKF groups are pooled (two-tailed Pearson correlation, *R²*=0.04, *p*=0.3958). In contrast, Shannon entropy for the OA latent cluster positively correlates with time spent in the open arms (pooled data; two-tailed Pearson correlation, *R²* = 0.61, \*\*\**p*=0.0001). **k,** k-Nearest Neighbor decoder achieved above 80% accuracy in distinguishing open arm vs. closed entry trials (\*\*\**p*<0.0001) in vehicle conditions, while it decreased to 70% of accuracy (\*\*\**p*<0.0001) following D1 activation (two-way ANOVA). Data are: mean ± SEM.

**Extended Data Fig. 7:**
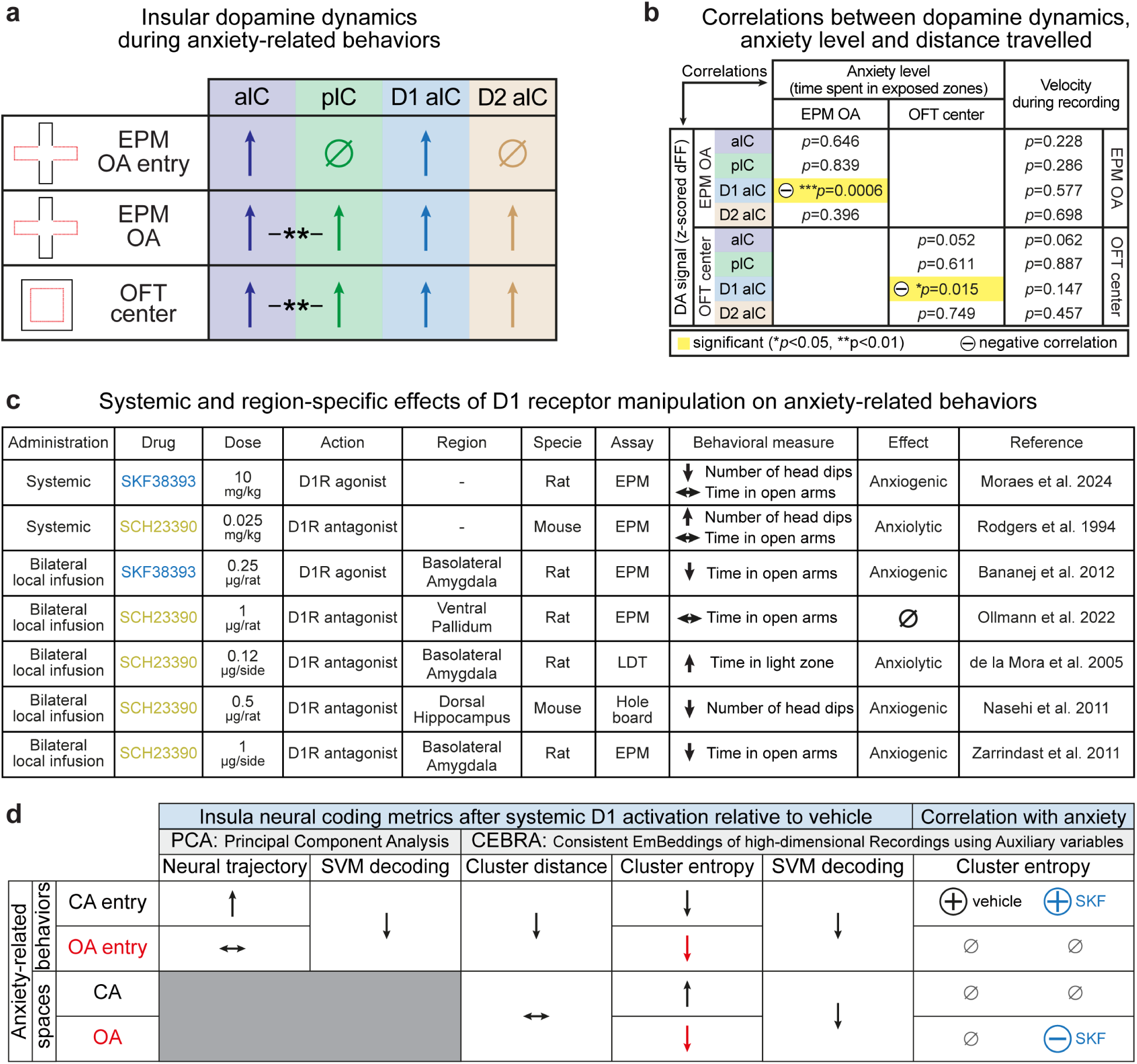
Summary tables. **a,** Dopamine dynamics in the anterior and posterior insula upon the entrance to EPM open arms, within open arms and in the center of the OFT. **b,** Correlations between dopamine signal, anxiety level and distance travelled. **c,** Table summarizing systemic and region-specific effects of D1 receptor pharmacological manipulations on anxiety-related behaviors. **d,** Anterior insula neural coding metrics after systemic D1 activation relative to vehicle and cluster entropy correlations with anxiety (summarizing Main results from Fig. 5 and 6).

