## Supplementary material for "Dopamine D1 receptor activation shapes anterior insula neural coding of anxiety": Statistical_Table

**Table 2. Statistical analyses**

| Figure | Variable | Group | Mice # | Statistical test | Comparison | DF | test value | p-value |
|---|---|---|---|---|---|---|---|---|
| 1c | VTADA - insula projector # | aIC<br>pIC | 5<br>5 | 2-way ANOVA | Hemisphere (ipsi/contra) main effect | 1,16 | F=15.7 | **p=0.0011 |
| | | | | | Brain region (aIC/pIC) main effect | 1,16 | F=4.91 | *p=0.0415 |
| | | | | | Hemisphere x Brain region interaction | 1,16 | F=4.44 | *p=0.0512 |
| | | | | Tukey's multiple comparison te | Ipsi VTADA-aIC vs. ipsi VTADA-pIC | 1,16 | q=4.32 | *p=0.0342 |
| | | | | | Ipsi VTADA-aIC vs. contra VTADA-aIC | 1,16 | q=6.08 | **p=0.0028 |
| | | | | | Ipsi VTADA-aIC vs. contra VTADA-pIC | 1,16 | q=6.18 | *p=0.0024 |
| 1d | SNcDA - insula projector # | aIC<br>pIC | 5<br>5 | 2-way ANOVA | Hemisphere (ipsi/contra) main effect | 1,16 | F=11 | **p=0.0044 |
| | | | | | Brain region (aIC/pIC) | 1,16 | F=3.34 | p=0.0863 |
| | | | | | Hemisphere x Brain region interaction | 1,16 | F=1.57 | p=0.2288 |
| 1g | VTADA-insula terminals | aIC<br>pIC | 14<br>14 | 2-way ANOVA | Layer | 3,104 | F=0.175 | p=0.9131 |
| | | | | | Brain region (aIC/pIC) main effect | 1,104 | F=8.87 | **p=0.0036 |
| | | | | | Layer x Brain region interaction | 3,104 | F=0.489 | p=0.6909 |
| 1h | SNcDA-insula terminals | aIC<br>pIC | 10<br>10 | 2-way ANOVA | Layer | 3,72 | F=0.127 | p=0.9440 |
| | | | | | Brain region (aIC/pIC) | 1,72 | F=2.17 | p=0.1450 |
| | | | | | Layer x Brain region interaction | 3,72 | F=3.19 | *p=0.0288 |
| | | | | Tukey's multiple comparison te | Layer 6: aIC vs. pIC | 3,72 | q=4.78 | **p=0.0012 |
| 1l | VTA-aIC evoked DA release | mCherry<br>rsChRmin<br>e | 2<br>3 | 2-way ANOVA | Group | 1, 356 | F=2.33 | p=0.1278 |
| | | | | | Period (baseline/laser) main effect | 1, 356 | F=4.22 | *p=0.0407 |
| | | | | | Group x Period interaction | 1, 356 | F=4.41 | *p=0.0365 |
| | | | | Tukey's multiple comparison te | Laser: rsChRmine vs. mCherry | 1, 356 | q=4.64 | **p=0.0062 |
| 1m | dLight aIC entries into CA and OA | aIC | 18 | 2-way ANOVA | Space (center/CA/OA) | 1,68 | F=0.46 | p=0.4997 |
| | | | | | Event (CA entry/OA entry) | 1,68 | F=3.55 | p=0.0638 |
| | | | | | Space x Event interaction | 1,68 | F=27.2 | ***p<0.0001 |
| | | | | Tukey's multiple comparison te | Center (CT) to closed arm (CA) entry | 1,68 | q=4.53 | *p=0.0108 |
| | | | | | Center (CT) to closed arm (OA) entry | 1,68 | q=5.89 | ***p=0.0005 |
| | | | | | CA entry to OA entry | 1,68 | q=7.10 | ***p<0.0001 |
| 1n | dLight pIC entries into CA and OA | pIC | 10 | 2-way ANOVA | Space (center/CA/OA) | 1,36 | F=0.155 | p=0.6957 |
| | | | | | Event (CA entry/OA entry) | 1,36 | F=0.515 | p=0.4774 |
| | | | | | Space x Event interaction | 1,36 | F=1.78 | p=0.1902 |
| 1p | DA release in the EPM | aIC<br>pIC | 18<br>10 | 2-way ANOVA | Brain region (aIC/pIC) | 1,52 | F=0.35 | p=0.5565 |
| | | | | | Arm main effect | 1,52 | F=92.6 | ***p<0.0001 |
| | | | | | Brain region x Arm interaction | 1,52 | F=21.6 | ***p<0.0001 |
| | | | | Tukey's multiple comparison te | aIC closed vs. aIC open | 1,52 | q=16.9 | ***p<0.0001 |
| | | | | | pIC closed vs. pIC open | 1,52 | q=4.38 | *p=0.0160 |
| | | | | | aIC open vs. pIC open | 1,52 | q=5.24 | **p=0.0028 |

| Figure | Variable | Group | Mice # | Statistical test | Comparison | DF | test value | p-value |
|---|---|---|---|---|---|---|---|---|
| 2b | Dopaminoceptive neuron density | D1<br>D2 | 13<br>8 | 2-way ANOVA | Recepteur (D1/D2) main effect | 1,40 | F=134 | ***p<0.0001 |
| | | | | | Brain region (aIC/pIC) | 1,40 | F=0.821 | p=0.3703 |
| | | | | | Recepteur x Brain region interaction | 1,40 | F=4.36 | *p=0.0432 |
| | | | | Holm-Sidak's multiple comparison test | aIC: D1 vs. D2 | 1,40 | t=9.66 | ***p<0.0001 |
| | | | | | pIC: D1 vs. D2 | 1,40 | t=6.71 | ***p<0.0001 |
| | | | | | D1: aIC vs. pIC | 1,40 | t=2.34 | *p=0.0480 |
| 2c | Dopaminoceptive neuron density | D1<br>D2 | 13<br>8 | 3-way ANOVA | Layer main effect | 3,160 | F=19,02 | ***p<0.0001 |
| | | | | | Brain region | 1,160 | F=1.462 | p=0.2284 |
| | | | | | Receptor main effect | 1,160 | F=238.7 | ***p<0.0001 |
| | | | | | Layer x Brain region x Receptor interaction | 3,160 | F=3.454 | *p=0.0180 |
| | | | | Tukey's multiple comparison test | L1 D1 aIC vs. D2 aIC | 3,160 | q=2.003 | p=0.989 |
| | | | | | L1 D1 pIC vs. D2 pIC | 3,160 | q=0.4690 | p>0.999 |
| | | | | | L2/3 D1 aIC vs. D2 aIC | 3,160 | q=16.88 | ***p<0.0001 |
| | | | | | L2/3 D1 pIC vs. D2 pIC | 3,160 | q=8.724 | ***p<0.0001 |
| | | | | | L2/3: D1 aIC vs. D1 pIC | 3,160 | q=9.408 | ***p=0.0002 |
| | | | | | L5 D1 aIC vs. D2 aIC | 3,160 | q=9.309 | ***p<0.0001 |
| | | | | | L5 D1 pIC vs. D2 pIC | 3,160 | q=6.501 | ***p=0.0009 |
| | | | | | L6 D1 aIC vs. D2 aIC | 3,160 | q=8.274 | ***p<0.0001 |
| | | | | | L6 D1 pIC vs. D2 pIC | 3,160 | q=9.886 | ***p<0.0001 |
| 2e | DA signal in EPM (D1 and D2 neurons aIC) | D1<br>D2 | 12<br>8 | 2-way ANOVA | Recepteur (D1/D2) | 1,36 | F=0.695 | p=0.4100 |
| | | | | | Arm main effect | 1,36 | F=48.9 | ***p<0.0001 |
| | | | | | Recepteur x Arm interaction | 1,36 | F=0.414 | p=0.5243 |
| 2g | DA signal in OFT (D1 and D2 neurons aIC) | D1<br>D2 | 12<br>8 | 2-way ANOVA | Recepteur (D1/D2) | 1,36 | F=1.84 | p=0.1833 |
| | | | | | Zone main effect | 1,36 | F=131 | ***p<0.0001 |
| | | | | | Recepteur x Zone interaction | 1,36 | F=2.90 | p=0.0973 |
| 2h | DA signal in open arms (D1 and D2 aIC) | D1<br>D2 | 12<br>8 | Two-tailed Pearson correlation | aIC D1:z(OA) vs. time in open arms | 1,10 | $R^2$ = 0.71 | ***p=0.0006 |
| | | | | | aIC D2: z(OA) vs. time in open arms | 1,6 | $R^2$ = 0.1222 | p=0.3960 |
| 2i | DA signal in OFT center (D1 and D2 aIC) | D1<br>D2 | 12<br>8 | Two-tailed Pearson correlation | aIC D1:z(OFT CT) vs. time in OFT CT | 1,10 | $R^2$ = 0.4596 | *p=0.0154 |
| | | | | | aIC D2: z(OFT CT) vs. time in OFT CT | 1,6 | $R^2$ = 0.01832 | p=0.7493 |

| Figure | Variable | Group | Mice # | Statistical test | Comparison | DF | test value | p-value |
|---|---|---|---|---|---|---|---|---|
| 3c | Time in open arms | Vehicle<br>SKF | 25<br>25 | Two-tailed unpaired t-test | Vehicle vs. SKF | | t=2.106 | *p=0.0405 |
| 3d | Distance travelled | Vehicle<br>SKF | 25<br>25 | Two-tailed unpaired t-test | Vehicle vs. SKF | | t=0.003 | p=0.9980 |
| 3f | Time in OFT center | Vehicle<br>SKF | 13<br>13 | Two-tailed unpaired t-test | Vehicle vs. SKF | | t=2.811 | **p=0.0097 |
| 3g | Distance travelled | Vehicle<br>SKF | 13<br>13 | Two-tailed unpaired t-test | Vehicle vs. SKF | | t=1.042 | p=0.3078 |
| 3j | cFOS+ cell density | Vehicle<br>SKF | 12<br>11 | 2-way ANOVA | Region main effect | 4,103 | F=34.2 | ***p<0.0001 |
| | | | | | Group (Veh vs SKF) main effect | 1,103 | F=49.2 | ***p<0.0001 |
| | | | | | Region x Group interaction | 4,103 | F=6.54 | ***p<0.0001 |
| | | | | Tukey's multiple comparison test | Vehicle: | | | |
| | | | | | aIC vs. pIC | 4,103 | 3.79 | p=0.0640 |
| | | | | | aIC vs. BLA | 4,103 | 1,91 | p=0.6621 |
| | | | | | aIC vs. S1 | 4,103 | 7,39 | ***p<0.0001 |
| | | | | | aIC vs. Nac | 4,103 | 5 | **p=0.0054 |
| | | | | | SKF: | | | |
| | | | | | aIC vs. pIC | 4,103 | 8.73 | ***p<0.0001 |
| | | | | | aIC vs. BLA | 4,103 | 10.9 | ***p<0.0001 |
| | | | | | aIC vs. S1 | 4,103 | 14.9 | ***p<0.0001 |
| | | | | | aIC vs. Nac | 4,103 | 11.5 | ***p<0.0001 |
| | | | | | Vehicle vs. SKF: | | | |
| | | | | | aIC | 4,103 | 10.5 | ***p<0.0001 |
| | | | | | pIC | 4,103 | 4.89 | ***p=0.0008 |
| | | | | | Nac | 4,103 | 3.25 | *p=0.0234 |
| | | | | | BLA | 4,103 | 1.18 | p=0.4074 |
| | | | | | S1 | 4,103 | 2.43 | p=0.0889 |
| 3m | Time in open arms | Vehicle<br>SKF<br>SCH | 29<br>9<br>17 | One-way ANOVA | Vehicle vs. SKF vs. SCH | 2,52 | F=11.37 | ***p<0.0001 |
| | | | | Tukey's multiple comparison test | Vehicle vs. SKF | 2,52 | q=4.201 | *p=0.0123 |
| | | | | | Vehicle vs. SCH | 2,52 | q=3.796 | *p=0.0259 |
| | | | | | SKF vs SCH | 2,52 | q=6.702 | ***p<0.0001 |
| 3n | Distance travelled | Vehicle<br>SKF<br>SCH | 29<br>9<br>17 | One-way ANOVA | Vehicle vs. SKF vs. SCH | 2,52 | F=2.014 | p=0.1437 |
| 3r | Absolute firing rate | Excitated<br>Inhibited | 8<br>19 | Two-tailed Mann-Whitney test | Excited vs. Inhibited units | | U=11 | ***p=0.0002 |
| 3t | Time in open arms | eYFP OFF<br>eYFP ON | 7<br>7 | Two-tailed paired t-test | eYFP OFF vs. ON | | t=0.08 | p=0.9388 |
| | | ChR2 OFF<br>ChR2 ON | 7<br>7 | Two-tailed paired t-test | ChR2 OFF vs. ON | | t=3.361 | *p=0.0152 |
| 3u | Distance travelled | eYFP<br>ChR2 | 7<br>7 | 2-way ANOVA | Opsin | 1,24 | F=0.395 | p=0.5357 |
| | | | | | Light mode (OFF/ON) | 1,24 | F=1.27 | p=0.27 |
| | | | | | Opsin x Light mode interaction | 1,24 | F=0.0023 | p=0.9569 |

| Figure | Variable | Group | Mice # | Statistical test | Comparison | DF | test value | p-value |
|---|---|---|---|---|---|---|---|---|
| 4b | Open arm entries # | Vehicle<br>SKF | 9<br>9 | One-tailed unpaired t-test | Vehicle vs. SKF | | t=1.870 | *p=0.0399 |
| 4b | Time in open arms | Vehicle<br>SKF | 9<br>9 | Two-tailed unpaired t-test | Vehicle vs. SKF | | t=3.144 | **p=0.0063 |
| 4b | Head dips | Vehicle<br>SKF | 9<br>9 | Two-tailed unpaired t-test | Vehicle vs. SKF | | t=2.856 | *p=0.0114 |
| 4d | Population firing rate | Vehicle<br>SKF | 9<br>9 | 2-way ANOVA | Trial type main effect<br>Group<br>Trial type x Group interaction | 2,2112<br>2,2112<br>2,2112 | F=6.61<br>F=0.176<br>F=4.38 | **p=0.0014<br>p=0.6753<br>*p=0.0126 |
| | | | | Tukey's multiple comparison test | CA entry Vehicle vs. SKF<br>OA entry Vehicle vs. SKF<br>HD Vehicle vs. SKF<br>Vehicle: CA entry vs. HD<br>Vehicle: OA entry vs. HD | 2,2112<br>2,2112<br>2,2112<br>2,2112<br>2,2112 | q=2.66<br>q=0.43<br>q=3.26<br>q=5.28<br>q=5.39 | *p=0.0601<br>p=0.7615<br>*p=0.0214<br>***p=0.0006<br>***p=0.0004 |
| 4e | Proportion of units | Vehicle<br>SKF | 9<br>9 | Generalized linear mixed model | Excited units:<br>CA entry vs. HD<br>OA entry vs. HD<br>CA entry vs. OA entry<br><br>Inhibited units:<br>CA entry vs. HD<br>CA entry vs. OA entry<br><br>Non-respondong units:<br>CA entry vs. HD<br>CA entry vs. OA entry | | <br>z=2.199<br>z=2.332<br>z=-0.324<br><br><br>z=-1.49<br>z=-4.67.10-15<br><br>z=-0.83<br>z=0.407 | <br>*p=0.028<br>*p=0.020<br>p=0.746<br><br><br>p=0.136<br>p=1<br><br>p=0.41<br>p=0.684 |
| 4f | E/I ratio | Vehicle | 9 | One-way ANOVA | | 2,24 | F=4.484 | *p=0.0222 |
| | | | | Tukey's multiple comparison test | CA entry vs. OA entry<br>CA entry vs. HD<br>OA entry vs. HD | 2,24<br>2,24<br>2,24 | q=0.0439<br>q=3.646<br>q=3.69 | p=0.9995<br>*p=0.0421<br>*p=0.0394 |
| 4f | E/I ratio | SKF | 9 | One-way ANOVA | | 2,24 | F=1.879 | p=0.1745 |
| 4h | Proportion of units | Vehicle<br>SKF | 9<br>9 | Multivariate analysis of variance | | | F=4.839 | *p=0.03796 |
| | | | | Welch's t-tests | Same<br>Opposite<br>Selective<br>Non-responsive | | t=-2.473<br>t=-2.183<br>t=-2.214<br>t=-1.772 | *p=0.0264<br>*p=0.0489<br>*p=0.0442<br>p=0.0965 |

| Figure | Variable | Group | Mice # | Statistical test | Comparison | DF | test value | p-value |
|---|---|---|---|---|---|---|---|---|
| 5c | Neural trajectory length | Vehicle<br>SKF | 9<br>9 | 3-way ANOVA | Event main effect | 2,96 | F=12.57 | ***p<0.0001 |
| | | | | | Group main effect | 1,96 | F=11.68 | ***p=0.0009 |
| | | | | | Data type main effect | 1,96 | F=3329 | ***p<0.0001 |
| | | | | | Event x Group x Data type interaction | 2,96 | F=80.86 | ***p<0.0001 |
| | | | | Tukey's multiple comparison test | Vehicle: CA entry vs. HD | 2,96 | q=4.757 | *p=0.0481 |
| | | | | | Vehicle: CA entry vs. OA entry | 2,96 | q=13.84 | ***p<0.0001 |
| | | | | | Vehicle: OA entry vs. HD | 2,96 | q=9.082 | ***p<0.0001 |
| | | | | | SKF: CA entry vs. OA entry | 2,96 | q=9.866 | ***p<0.0001 |
| | | | | | SKF: CA entry vs. HD | 2,96 | q=14.19 | ***p<0.0001 |
| | | | | | SKF: OA entry vs. HD | 2,96 | q=4.329 | p=0.1075 |
| | | | | | CA entry: Data vs. Shuffle | 2,96 | q=16.83 | ***p<0.0001 |
| | | | | | OA entry: Data vs. Shuffle | 2,96 | q=2.24 | p=0.9193 |
| | | | | | HD: Data vs. Shuffle | 2,96 | q=11.21 | **p<0.0001 |
| 5f | SVM accuracy | Vehicle<br>SKF | 9<br>9 | 2-way ANOVA | Data type main effect | 1,108 | F=2252 | ***p<0.0001 |
| | | | | | Event main effect | 5,108 | F=23.2 | ***p<0.0001 |
| | | | | | Data type x Event interaction | 5,108 | F=37.7 | ***p<0.0001 |
| | | | | Tukey's multiple comparison test | Vehicle: | | | |
| | | | | | OA entry-CA entry vs. OA entry-HD | 5,108 | q=8.81 | ***p<0.0001 |
| | | | | | OA entry-HD vs. HD-CA entry | 5,108 | q=2.87 | p=0.6712 |
| | | | | | OA entry-CA entry vs. HD-CA entry | 5,108 | q=2.45 | p=0.8480 |
| | | | | | SKF: | | | |
| | | | | | OA entry-CA entry vs. OA entry-HD | 5,108 | q=0.606 | p=0.9999 |
| | | | | | OA entry-HD vs. HD-CA entry | 5,108 | q=1.85 | p=0.9767 |
| | | | | | OA entry-CA entry vs. HD-CA entry | 5,108 | q=16 | ***p<0.0001 |
| | | | | | Vehicle vs. SKF | | | |
| | | | | | OA entry-CA entry | 5,108 | q=8.2 | ***p<0.0001 |
| | | | | | OA entry-HD | 5,108 | q=13.5 | ***p<0.0001 |
| | | | | | HD-CA entry | 5,108 | q=3.48 | p=0.3745 |

| Figure | Variable | Group | Mice # | Statistical test | Comparison | DF | test value | p-value |
|---|---|---|---|---|---|---|---|---|
| 6c | Cluster distance | Vehicle<br>SKF | 9<br>9 | 2-way ANOVA | Snippet main effect<br>Group main effect<br>Snippet x Group interaction | 2,48<br>1,48<br>2,48 | F=468<br>F=48.9<br>F=236 | ***p<0.0001<br>***p<0.0001<br>***p<0.0001 |
| | | | | Tukey's multiple comparison test | Vehicle:<br>OA entry-CA entry vs. OA entry-HD<br>OA entry-CA entry vs. CA entry-HD<br>OA entry-HD vs. CA entry-HD | <br>2,48<br>2,48<br>2,48 | <br>q=5.84<br>q=9.60<br>q=3.75 | <br>***p=0.0004<br>***p<0.0001<br>*p=0.0285 |
| | | | | | SKF:<br>OA entry-CA entry vs. OA entry-HD<br>OA entry-CA entry vs. CA entry-HD<br>OA entry-HD vs. CA entry-HD | <br>2,48<br>2,48<br>2,48 | <br>q=45.1<br>q=45.2<br>q=0.09 | <br>***p<0.0001<br>***p<0.0001<br>p=0.9977 |
| | | | | | Vehicle vs. SKF<br>OA entry-CA entry<br>OA entry-HD<br>CA entry-HD | <br>2,48<br>2,48<br>2,48 | <br>q=30.7<br>q=8.61<br>q=4.95 | <br>***p<0.0001<br>***p<0.0001<br>**p=0.0010 |
| 6d | Cluster entropy | Vehicle<br>SKF | 9<br>9 | 2-way ANOVA | Snippet<br>Group main effect<br>Snippet x Group interaction | 2,48<br>1,48<br>2,48 | F=2.93<br>F=141<br>F=2.21 | p=0.0629<br>***p<0.0001<br>p=0.1203 |
| 6e | Cluster entropy<br>CA entry | Vehicle<br>SKF | 9<br>9 | Two-tailed Pearson correlation | CA entry Vehicle entropy vs. time open arms<br>CA entry SKF entropy vs. time open arms | 1,7<br>1,7 | $R^2$ = 0.56<br>$R^2$ = 0.78 | *p=0.0210<br>***p=0.0008 |
| 6e | Cluster entropy<br>OA entry | Vehicle<br>SKF | 9<br>9 | Two-tailed Pearson correlation | OA entry Vehicle entropy vs. time open arms<br>OA entry SKF entropy vs. time open arms | 1,7<br>1,7 | $R^2$ = 0.007<br>$R^2$ = 0.291 | p=0.8327<br>p=0.0668 |
| 6e | Cluster entropy<br>HD | Vehicle<br>SKF | 9<br>9 | Two-tailed Pearson correlation | HD Vehicle entropy vs. time open arms<br>HD SKF entropy vs. time open arms | 1,7<br>1,7 | $R^2$ = 0.07<br>$R^2$ = 0.11 | p=0.4937<br>p=0.3810 |
| 6f | SVM accuracy | Vehicle<br>SKF | 9<br>9 | 2-way ANOVA | Group main effect<br>Data type main effect<br>Group x Data type interaction | 1,32<br>1,32<br>1,32 | F=288.2<br>F=3004<br>F=322.4 | ***p<0.0001<br>***p<0.0001<br>***p<0.0001 |
| | | | | Tukey's multiple comparison test | Vehicle: Shuffle vs. Data<br>SKF: Shuffle vs. Data<br>Vehicle vs. SKF Data | 1,32<br>1,32<br>1,32 | q=72.76<br>q=36.85<br>q=34.93 | ***p<0.0001<br>***p<0.0001<br>***p<0.0001 |
| 6h | CA-OA barycenter distance | Vehicle<br>SKF | 9<br>9 | Two-tailed unpaired t-test | Vehicle vs. SKF | | t=1.033 | p=0.3170 |

| | | | | | | | | |
|---|---|---|---|---|---|---|---|---|
| 6f | Shannon entropy | Vehicle<br>SKF | 9<br>9 | 2-way ANOVA | Arm main effect<br>Group<br>Arm x Group interaction | 1,32<br>1,32<br>1,32 | F=61.4<br>F=0.150<br>F=11 | ***p<0.0001<br>p=0.7015<br>**p=0.0023 |
| | | | | Holm-Sidak's multiple comparison test | CA: Vehicle vs. SKF<br>OA: Vehicle vs. SKF<br>CA vs. OA Vehicle<br>CA vs. OA SKF | 1,32<br>1,32<br>1,32<br>1,32 | t=2.07<br>t=2.62<br>t=3.19<br>t=7.89 | *p=0.0462<br>*p=0.0264<br>**p=0.0095<br>***p<0.0001 |
| 6j | Cluster entropy<br>CA position | Vehicle<br>SKF | 9<br>9 | Two-tailed Pearson correlation | CA position Vehicle entropy vs. time open arms<br>CA position SKF entropy vs. time open arms | 1,7<br>1,7 | $R^2$ = 0.05<br>$R^2$ = 0.038 | p=0.5608<br>p=0.6147 |
| 6j | Cluster entropy<br>OA position | Vehicle<br>SKF | 9<br>9 | Two-tailed Pearson correlation | OA position Vehicle entropy vs. time open arms<br>OA position SKF entropy vs. time open arms | 1,7<br>1,7 | $R^2$ = 0.239<br>$R^2$ = 0.898 | p=0.1818<br>***p=0.0001 |
| 6k | SVM accuracy | Vehicle<br>SKF | 9<br>9 | 2-way ANOVA | Group main effect<br>Data type main effect<br>Group x Data type interaction | 1,32<br>1,32<br>1,32 | F=484<br>F=14.8<br>F=4.95 | ***p<0.0001<br>***p=0.0005<br>*p=0.0333 |
| | | | | Tukey's multiple comparison test | Vehicle: Shuffle vs. Data<br>SKF: Shuffle vs. Data<br>Vehicle vs. SKF Data | 1,32<br>1,32<br>1,32 | q=24.2<br>q=19.08<br>q=6.07 | ***p<0.0001<br>***p<0.0001<br>***p=0.0008 |

| Figure | Variable | Group | Mice # | Statistical test | Comparison | DF | test value | p-value |
|---|---|---|---|---|---|---|---|---|
| S1b | TH projector # | VTA<br>SNc | 5<br>5 | 2-way ANOVA | Region (VTA/SNc)<br>Hemisphere main effect<br>Brain region x Zone interaction | 1,16<br>1,16<br>1,16 | F=0.039<br>F=17.62<br>F=0.002 | p=0.8454<br>***p=0.0007<br>p=0.9617 |
| S1e | VTA-aIC evoked DA release | mCherry<br>rsChRmine | 2<br>3 | 2-way ANOVA | Opsin main effect<br>Period (baseline/laser) main effect<br>Opsin x Period interaction | 1, 140<br>1, 140<br>1, 140 | F=6.425<br>F=6.552<br>F=6.574 | *p=0.0124<br>*p=0.0115<br>*p=0.0114 |
| | | | | Tukey's multiple comparison test | rsChRmine: baseline vs. Laser stim | 1, 140 | q=5.124 | **p=0.0023 |
| S1j | DA signal in the OFT | aIC<br>pIC | 18<br>10 | 2-way ANOVA | Brain region main effect<br>Zone main effect<br>Brain region x Zone interaction | 1,52<br>1,52<br>1,52 | F=5.82<br>F=120<br>F=5.88 | *p=0.0194<br>***p<0.0001<br>*p=0.0189 |
| | | | | Tukey's multiple comparison test | aIC border vs. aIC center<br>pIC border vs. pIC center<br>aIC center vs. pIC center | 1,52<br>1,52<br>1,52 | q=15.8<br>q=7.53<br>q=4.84 | ***p<0.0001<br>***p<0.0001<br>**p=0.0065 |
| S1k | DA signal in open arms (aIC and pIC) | aIC<br>pIC | 18<br>10 | Two-tailed Pearson correlation | aIC: z(OA) vs. time in open arms<br>pIC: z(OA) vs. time in open arms | 1,16<br>1,8 | $R^2$ = 0.01349<br>$R^2$ = 0.00547 | p=0.6462<br>p=0.8391 |
| S1l | DA signal in OFT center (aIC and pIC) | aIC<br>pIC | 18<br>10 | Two-tailed Pearson correlation | aIC: z(OFT CT) vs. time in OFT CT<br>pIC: z(OFT CT) vs. time in OFT CT | 1,16<br>1,8 | $R^2$ = 0.2151<br>$R^2$ = 0.03384 | p=0.0525<br>p=0.6109 |
| S1m | DA signal in open arms (aIC and pIC) | aIC<br>pIC | 18<br>10 | Two-tailed Pearson correlation | aIC: z(OA) vs. velocity in open arms<br>pIC: z(OA) vs. velocity in open arms | 1,16<br>1,8 | $R^2$ = 0.0895<br>$R^2$ = 0.1407 | p=0.2279<br>p=0.2855 |
| S1n | DA signal in OFT center (aIC and pIC) | aIC<br>pIC | 18<br>10 | Two-tailed Pearson correlation | aIC: z(OFT CT) vs. velocity in OFT CT<br>pIC: z(OFT CT) vs. velocity inOFT CT | 1,16<br>1,8 | $R^2$ = 0.2015<br>$R^2$ = 0.0027 | p=0.0617<br>p=0.8866 |
| S1o | aIC and pIC DA transient frequency in the EPM | aIC<br>pIC | 18<br>10 | 2-way ANOVA | Brain region main effect<br>Arm<br>Brain region x Arm interaction | 1,52<br>1,52<br>1,52 | F=27.5<br>F=0.15<br>F=0.0482 | ***p<0.0001<br>p=0.7003<br>p=0.8271 |
| S1p | aIC and pIC DA transient amplitude in the EPM | aIC<br>pIC | 18<br>10 | 2-way ANOVA | Brain region main effect<br>Arm<br>Brain region x Arm interaction | 1,52<br>1,52<br>1,52 | F=1.78<br>F=0.0305<br>F=0.2 | p=0.1886<br>p=0.8620<br>p=0.6564 |

| Figure | Variable | Group | Mice # | Statistical test | Comparison | DF | test value | p-value |
|---|---|---|---|---|---|---|---|---|
| S2b | Raw number of D1-neurons in subpopulations | aIC<br>pIC | 6<br>6 | 2-way ANOVA | Subpopulation main effect | 1,20 | F=10.8 | ***p<0.0001 |
| | | | | | Brain region | 1,20 | F=0.7961 | p=0.3829 |
| | | | | | Subpopulation x Brain region interaction | 1,20 | F=0.1018 | p=0.7530 |
| S2b | Raw number of D2-neurons in subpopulations | aIC<br>pIC | 8<br>8 | 2-way ANOVA | Subpopulation | 1,28 | F=0.8323 | p=0.3694 |
| | | | | | Brain region | 1,28 | F=0.1208 | p=0.7308 |
| | | | | | Subpopulation x Brain region interaction | 1,28 | F=0.8724 | p=0.3583 |
| S2c | Proportion of D1-neurons in subpopulations | aIC<br>pIC | 6<br>6 | 2-way ANOVA | Subpopulation main effect | 1,20 | F=10.8 | **p=0.0037 |
| | | | | | Brain region | 1,20 | F=0.711 | p=0.409 |
| | | | | | Subpopulation x Brain region interaction | 1,20 | F=0.659 | p=0.4266 |
| S2c | Proportion of D2-neurons in subpopulations | aIC<br>pIC | 8<br>8 | 2-way ANOVA | Subpopulation main effect | 1,28 | F=4.99 | *p=0.0337 |
| | | | | | Brain region | 1,28 | F=0.169 | p=0.6844 |
| | | | | | Subpopulation x Brain region interaction | 1,28 | F=0.0663 | p=0.7987 |
| S2d | Drd1 and Drd2 mRNA expression in the aIC | *Drd1*<br>*Drd2* | 18<br>18 | 2-way ANOVA | Receptor main effect | 1,60 | F=69.8 | ***p<0.0001 |
| | | | | | Subpopulation main effect | 1,60 | F=42.6 | ***p<0.0001 |
| | | | | | Receptor x Region interaction | 1,60 | F=4.89 | *p=0.0308 |
| | | | | Tukey's multiple comparison test | vGLUT Drd1 mRNA vs. vGLUT Drd2 mRNA | 1,60 | q=6.14 | ***p=0.0003 |
| | | | | | vGLUT Drd1 mRNA vs. vGAT Drd1 mRNA | 1,60 | q=8.74 | ***p<0.0001 |
| | | | | | vGAT Drd1 mRNA vs. vGAT Drd2 mRNA | 1,60 | q=10.6 | ***p<0.0001 |
| | | | | | vGLUT Drd2 mRNA vs. vGAT Drd2 mRNA | 1,60 | q=4.32 | *p=0.0174 |
| S2e | Drd1 and Drd2 mRNA expression in the pIC | *Drd1*<br>*Drd2* | 18<br>18 | 2-way ANOVA | Receptor main effect | 1,60 | F=9.76 | **p=0.0027 |
| | | | | | Subpopulation main effect | 1,60 | F=4.7 | *p=0.0341 |
| | | | | | Receptor x Region interaction | 1,60 | F=5.90 | *p=0.0182 |
| | | | | Tukey's multiple comparison test | vGLUT Drd2 mRNA vs. vGAT Drd2 mRNA | 1,60 | q=4.60 | **p=0.0099 |
| | | | | | vGAT Drd1 mRNA vs. vGAT Drd2 mRNA | 1,60 | q=5.55 | **p=0.0013 |
| S2f | Summary diagram | | | Two-tailed unpaired t tests | vGLUT: D1 aIC vs. Drd1 aIC | | t=0.8057 | p=0.4287 |
| | | | | | vGAT: D1 aIC vs. Drd1 aIC | | t=0.9481 | p=0.3529 |
| | | | | | vGLUT: D2 aIC vs. Drd2 aIC | | t=1.761 | p=0.0906 |
| | | | | | vGAT: D2 aIC vs. Drd2 aIC | | t=0.3156 | p=0.7550 |
| | | | | | vGLUT: D1 pIC vs. Drd1 pIC | | t=2.215 | *p=0.0375 |
| | | | | | vGAT: D1 pIC vs. Drd1 pIC | | t=4.687 | ***p=0.0001 |
| | | | | | vGLUT: D2 pIC vs. Drd2 pIC | | t=2.431 | *p=0.0229 |
| | | | | | vGAT: D2 pIC vs. Drd2 pIC | | t=2.867 | **p=0.0083 |
| S2h | D1 aIC EPM entries into CA and OA | D1 | 12 | 2-way ANOVA | Space (center/CA/OA) | 1,44 | F=2.48 | p=0.1221 |
| | | | | | Event (CA entry/OA entry) main effect | 1,44 | F=4.56 | *p=0.0384 |
| | | | | | Space x Event interaction | 1,44 | F=31.1 | ***p<0.0001 |
| | | | | Tukey's multiple comparison test | Center (CT) to closed arm (CA) entry | 1,44 | q=4 | *p=0.0339 |
| | | | | | Center (CT) to open arm (OA) entry | 1,44 | q=7.16 | ***p<0.0001 |
| | | | | | CA entry to OA entry | 1,44 | q=7.72 | ***p<0.0001 |

| | | | | | | | | |
|---|---|---|---|---|---|---|---|---|
| S2i | D2 aIC EPM entries into CA and OA | D2 | 8 | 2-way ANOVA | Space (center/CA/OA)<br>Event (CA entry/OA entry)<br>Space x Event interaction | 1,28<br>1,28<br>1,28 | F=0.0243<br>F=0.0498<br>F=8.74 | p=0.8772<br>p=0.8251<br>**p=0.0063 |
| | | | | Tukey's multiple comparison test | Center (CT) to closed arm (CA) entry<br>Center (CT) to open arm (OA) entry<br>CA entry to OA entry | 1,28<br>1,28<br>1,28 | q=2.8<br>q=3.11<br>q=3.18 | p=0.2194<br>p=0.1477<br>p=0.1351 |
| S2j | DA transient frequency onto D1 or D2 aIC neurons in the EPM | D1<br>D2 | 12<br>8 | 2-way ANOVA | Recepteur (D1/D2)<br>Arm<br>Recepteur x Arm interaction main effect | 1,36<br>1,36<br>1,36 | F=0.306<br>F=0.22<br>F=0.0541 | p=0.5836<br>p=0.6418<br>p=0.8173 |
| S2k | DA transient amplitude onto D1 or D2 aIC neurons in the EPM | D1<br>D2 | 12<br>8 | 2-way ANOVA | Recepteur (D1/D2)<br>Arm<br>Recepteur x Arm interaction | 1,36<br>1,36<br>1,36 | F=0.741<br>F=3.76<br>F=2.02 | p=0.3950<br>p=0.0602<br>p=0.1643 |
| S2l | DA release onto D1 or D2 aIC neurons in EPM open arms | D1<br>D2 | 12<br>8 | Two-tailed Pearson correlation | aIC D1: z(OA) vs. velocity in open arms<br>aIC D2: z(OA) vs. velocity in open arms | 1,10<br>1,6 | $R^2$ = 0.0322<br>$R^2$ = 0.0268 | p=0.5768<br>p=0.6982 |
| S2m | DA release onto D1 or D2 aIC neurons in OFT center | D1<br>D2 | 12<br>8 | Two-tailed Pearson correlation | aIC D1: z(OFT CT) vs. velocity in OFT CT<br>aIC D2: z(OFT CT) vs. velocity in OFT CT | 1,10<br>1,6 | $R^2$ = 0.1984<br>$R^2$ = 0.0952 | p=0.1468<br>p=0.4572 |
| S2q | Relative synaptic density of D1 aIC and pIC outputs | aIC<br>pIC | 5<br>6 | 2-way ANOVA | Output region main effect<br>Input<br>Output region x Input interaction | 11,107<br>1,107<br>11,107 | F=18.54<br>F=0.0174<br>F=3.82 | ***p<0.0001<br>p=0.8954<br>***p=0.0001 |
| | | | | Tukey's multiple comparison test | contra-aIC D1+ aIC vs. D1+ pIC<br>contra-pIC D1+ aIC vs. D1+ pIC | 11,107<br>11,107 | q=5.318<br>t=6.186 | ***p=0.0003<br>***p<0.0001 |

| Figure | Variable | Group | Mice # | Statistical test | Comparison | DF | test value | p-value |
|---|---|---|---|---|---|---|---|---|
| S3b | Time in open arms | Vehicle<br>SKF | 12<br>9 | Two-tailed unpaired t-test | Vehicle vs. SKF | | t=2.972 | **p=0.0078 |
| S3b | Distance travelled | Vehicle<br>SKF | 12<br>9 | Two-tailed unpaired t-test | Vehicle vs. SKF | | t=1.136 | p=0.2702 |
| S3c | Time in open arms | Vehicle<br>SCH | 17<br>17 | Two-tailed unpaired t-test | Vehicle vs. SCH | | t=2.218 | *p=0.0337 |
| S3c | Distance travelled | Vehicle<br>SCH | 17<br>17 | Two-tailed unpaired t-test | Vehicle vs. SCH | | t=0.4946 | p=0.6242 |
| S3g | Distance travelled | eYFP<br>ChR2 | 7<br>7 | 2-way ANOVA | Opsin<br>Light mode (OFF/ON)<br>Opsin x Light mode interaction | 1,24<br>1,24<br>1,24 | F=0.395<br>F=1.27<br>F=0.0023 | p=0.5357<br>p=0.27<br>p=0.9569 |

| Figure | Variable | Group | Mice # | Statistical test | Comparison | DF | test value | p-value |
|---|---|---|---|---|---|---|---|---|
| S4b | Time spent in EPM | Vehicle<br>SKF | 9<br>9 | 2-way ANOVA | Zone main effect<br>Group main effect<br>Zone x Group interaction | 2,48<br>1,48<br>2,48 | F=6.61<br>F=1.16x10-14<br>F=4.38 | ***p<0.0001<br>p>0.9999<br>***p<0.0001 |
| | | | | Tukey's multiple comparison test | CA Vehicle vs. SKF<br>OA Vehicle vs. SKF<br>CT Vehicle vs. SKF | 2,48<br>2,48<br>2,48 | q=6.61<br>q=5.22<br>q=1.39 | ***p=0.0003<br>**p=0.0071<br>p=0.9219 |
| S4b | Distance travelled | Vehicle<br>SKF | 9<br>9 | Two-tailed unpaired t-test | Vehicle vs. SKF | | t=0.4946 | p=0.6242 |
| S4b | Closed arm entries # | Vehicle<br>SKF | 9<br>9 | Two-tailed unpaired t-test | Vehicle vs. SKF | | t=1.208 | p=0.2446 |
| S4d | Climbing in CA | Vehicle<br>SKF | 9<br>9 | Two-tailed unpaired t-test | Vehicle vs. SKF | | t=1.035 | p=0.3170 |
| S4d | Time head-dipping | Vehicle<br>SKF | 9<br>9 | Two-tailed unpaired t-test | Vehicle vs. SKF | | t=3.122 | **p=0.007 |
| S4d | Time immobile | Vehicle<br>SKF | 9<br>9 | Two-tailed unpaired t-test | Vehicle vs. SKF | | t=0.5581 | p=0.5850 |
| S4d | Time sniffing | Vehicle<br>SKF | 9<br>9 | Two-tailed unpaired t-test | Vehicle vs. SKF | | t=0.0119 | p=0.9906 |
| S4d | Time sniffing borders | Vehicle<br>SKF | 9<br>9 | Two-tailed unpaired t-test | Vehicle vs. SKF | | t=0.1686 | p=0.8684 |
| S4d | Time scanning | Vehicle<br>SKF | 9<br>9 | Two-tailed unpaired t-test | Vehicle vs. SKF | | t=0.0196 | p=0.9846 |
| S4d | Instantaneous speed | Vehicle<br>SKF | 9<br>9 | Two-tailed unpaired t-test | Vehicle vs. SKF | | t=0.9897 | p=0.338 |
| S4f | Averaged unit number | Vehicle<br>SKF | 9<br>9 | Two-tailed unpaired t-test | Vehicle vs. SKF | | t=2.283 | *p=0.0364 |
| S4i | Proportion of units | Vehicle<br>SKF | 9<br>9 | Multivariate analysis of variance | | | F=2.363 | *p=0.05 |
| | | | | | Exc OA & Exc CA<br>Inh OA & Exc CA<br>Neu OA & Exc CA<br>Exc OA & Inh CA:<br>Inh OA & Inh CA:<br>Neu OA & Neu CA:<br>Neu OA & Inh CA:<br>Inh OA & Neu CA:<br>Exc OA & Neu CA | | t = -3.2981<br>t = -3.1261<br>t = -1.9729<br>t = -2.2034<br>t = -2.0423<br>t = 1.8626<br>t = 1.7368<br>t = 0.9762<br>t = -0.6028 | **p = 0.0061<br>**p = 0.0071<br>p = 0.0708<br>p = 0.0500<br>p = 0.0687<br>p = 0.0820<br>p = 0.1190<br>p = 0.3435<br>p = 0.5551 |
| S4j | Unit response between OA entry and HD | Vehicle<br>SKF | 9<br>9 | Multivariate analysis of variance | | | F=0.001 | p=0.9906 |
| | | | | Welch's t-tests | Same<br>Opposite<br>Selective<br>Non-responsive | | t=-0.085<br>t=-0.022<br>t=0.026<br>t=0.028 | p=0.9330<br>p=0.9830<br>p=0.9800<br>p=0.9783 |

| | | | | | | | |
|---|---|---|---|---|---|---|---|
| S4h | Unit response between CA entry and HD | Vehicle<br>SKF | 9<br>9 | Multivariate analysis of variance | | F=2.344 | p=0.1099 |
| | | | | Welch's t-tests | Same<br>Opposite<br>Selective<br>Non-responsive | t=-1.526<br>t=-1.626<br>t=-1.541<br>t=-1.325 | p=0.1471<br>p=0.1246<br>p=0.1432<br>p=0.2038 |

| Figure | Variable | Group | Mice # | Statistical test | Comparison | DF | test value | p-value |
|---|---|---|---|---|---|---|---|---|
| S5a | PC1-3: variance explained | Vehicle<br>SKF | 9<br>9 | 2-way ANOVA | Event | 2,48 | F=0.653 | p=0.5250 |
| | | | | | Group | 1,48 | F=3.77 | p=0.0582 |
| | | | | | Event x Group interaction | 2,48 | F=54 | ***p<0.0001 |
| | | | | Tukey's multiple comparison te | Vehicle: CA entry vs. OA entry | 2,48 | q=4.73 | *p=0.0189 |
| | | | | | Vehicle: CA entry vs. HD | 2,48 | q=9.26 | ***p<0.001 |
| | | | | | Vehicle: OA entry vs. HD | 2,48 | q=4.53 | *p=0.0274 |
| | | | | | SKF: CA entry vs. OA entry | 2,48 | q=6.16 | ***p=0.0009 |
| | | | | | SKF: CA entry vs. HD | 2,48 | q=11.5 | ***p<0.0001 |
| | | | | | SKF: OA entry vs. HD | 2,48 | q=5.36 | **p=0.0053 |
| | | | | | CA entry: Vehicle vs. SKF | 2,48 | q=8.97 | ***p<0.0001 |
| | | | | | OA entry: Vehicle vs. SKF | 2,48 | q=1.92 | p=0.7524 |
| | | | | | HD: Vehicle vs. SKF | 2,48 | q=11.8 | ***p<0.0001 |
| S5b | Relative trajectory length | Vehicle<br>SKF | 9<br>9 | 2-way ANOVA | Event | 2,48 | F=0.198 | p=0.198 |
| | | | | | Group | 1,48 | F=1.79 | p=0.8214 |
| | | | | | Event x Group interaction | 2,48 | F=135 | ***p<0.0001 |
| | | | | Tukey's multiple comparison te | Vehicle: CA entry vs. OA entry | 2,48 | q=9.77 | ***p<0.0001 |
| | | | | | Vehicle: CA entry vs. HD | 2,48 | q=16.7 | ***p<0.0001 |
| | | | | | Vehicle: OA entry vs. HD | 2,48 | q=6.94 | ***p=0.0001 |
| | | | | | SKF: CA entry vs. OA entry | 2,48 | q=10.2 | ***p<0.0001 |
| | | | | | SKF: CA entry vs. HD | 2,48 | q=15.9 | ***p<0.0001 |
| | | | | | SKF: OA entry vs. HD | 2,48 | q=5.70 | **p=0.0026 |
| | | | | | CA entry: Vehicle vs. SKF | 2,48 | q=18.6 | ***p<0.0001 |
| | | | | | OA entry: Vehicle vs. SKF | 2,48 | q=1.35 | p=0.9291 |
| | | | | | HD: Vehicle vs. SKF | 2,48 | q=14 | ***p<0.0001 |
| S5d | SVM accuracy | Vehicle<br>SKF | 9<br>9 | 2-way ANOVA | Data type main effect | 1,36 | F=556 | ***p<0.0001 |
| | | | | | Event main effect | 1,36 | F=21.6 | ***p<0.0001 |
| | | | | | Data type x Event interaction | 1,36 | F=24.1 | ***p<0.0001 |
| | | | | Tukey's multiple comparison test | Vehicle: shuffle vs. Data | 1,36 | F=28.5 | ***p<0.0001 |
| | | | | | SKF: shuffle vs. Data | 1,36 | F=18.7 | ***p<0.0001 |
| | | | | | Data: Vehicle vs. SKF | 1,36 | F=9.56 | ***p<0.0001 |

| Figure | Variable | Group | Mice # | Statistical test | Comparison | DF | test value | p-value |
|---|---|---|---|---|---|---|---|---|
| S6b | Cluster distance CA entry-OA entry | Vehicle<br>SKF | 9<br>9 | Two-tailed Pearson correlation | Cluster distance Vehicle vs. time open arms<br>Cluster distance SKF vs. time open arms<br>Cluster distance Vehicle + SKF vs. time open arms | 1,7<br>1,7<br>1,7 | $R^2$ = 0.224<br>$R^2$ = 0.085<br>$R^2$ = 0.47 | p=0.1980<br>p=0.4478<br>**p=0.0017 |
| S6b | Cluster distance OA entry-HD | Vehicle<br>SKF | 9<br>9 | Two-tailed Pearson correlation | Cluster distance Vehicle vs. time open arms<br>Cluster distance SKF vs. time open arms<br>Cluster distance Vehicle + SKF vs. time open arms | 1,7<br>1,7<br>1,7 | $R^2$ = 0.049<br>$R^2$ = 0.011<br>$R^2$ = 0.39 | p=0.568<br>p=0.789<br>**p=0.0056 |
| S6b | Cluster distance CA entry-HD | Vehicle<br>SKF | 9<br>9 | Two-tailed Pearson correlation | Cluster distance Vehicle vs. time open arms<br>Cluster distance SKF vs. time open arms<br>Cluster distance Vehicle + SKF vs. time open arms | 1,7<br>1,7<br>1,7 | $R^2$ = 0.06<br>$R^2$ = 0.06<br>$R^2$ = 0.41 | p=0.5181<br>p=0.5181<br>**p=0.0041 |
| S6c | Cluster entropy CA entry | Vehicle<br>SKF | 9<br>9 | Two-tailed Pearson correlation | Cluster entropy Vehicle + SKF vs. time open arms | 1,7 | $R^2$ = 0.09 | p=0.2154 |
| S6c | Cluster entropy OA entry | Vehicle<br>SKF | 9<br>9 | Two-tailed Pearson correlation | Cluster entropy Vehicle + SKF vs. time open arms | 1,7 | $R^2$ = 0.39 | **p=0.006 |
| S6c | Cluster entropy HD | Vehicle<br>SKF | 9<br>9 | Two-tailed Pearson correlation | Cluster entropy Vehicle + SKF vs. time open arms | 1,7 | $R^2$ = 0.36 | **p=0.008 |
| S6d | SVM accuracy | Vehicle<br>SKF | 9<br>9 | 2-way ANOVA | Group main effect<br>Data type main effect<br>Group x Data type interaction | 1,32<br>1,32<br>1,32 | F=247<br>F=3745<br>F=265 | ***p<0.0001<br>***p<0.0001<br>***p<0.0001 |
| | | | | Tukey's multiple comparison test | Vehicle: Shuffle vs. Data<br>SKF: Shuffle vs. Data<br>Vehicle vs. SKF Data | 1,32<br>1,32<br>1,32 | q=77.5<br>q=44.9<br>q=32 | ***p<0.0001<br>***p<0.0001<br>***p<0.0001 |
| S6h | Cluster distance CA-OA position | Vehicle<br>SKF | 9<br>9 | Two-tailed Pearson correlation | Cluster distance Vehicle vs. time open arms<br>Cluster distance SKF vs. time open arms<br>Cluster distance Vehicle + SKF vs. time open arms | 1,7<br>1,7<br>1,7 | $R^2$ = 0.3901<br>$R^2$ = 0.3898<br>$R^2$ = 0.09 | p=0.0721<br>p=0.0723<br>p=0.5070 |
| S6i | Shannon entropy | Vehicle<br>SKF | 9<br>9 | 2-way ANOVA | Arm main effect<br>Group<br>Arm x Group interaction | 1,32<br>1,32<br>1,32 | F=59.9<br>F=0.893<br>F=16.9 | ***p<0.0001<br>p=0.3517<br>***p=0.0003 |
| | | | | Holm-Sidak's multiple comparison test | CA: Vehicle vs. SKF<br>OA: Vehicle vs. SKF<br>CA vs. OA Vehicle<br>CA vs. OA SKF | 1,32<br>1,32<br>1,32<br>1,32 | t=2.24<br>t=3.58<br>t=2.56<br>t=8.38 | *p=0.0322<br>**p=0.0034<br>*p=0.0304<br>***p<0.0001 |
| S6j | Cluster entropy CA position | Vehicle<br>SKF | 9<br>9 | Two-tailed Pearson correlation | Cluster entropy Vehicle + SKF vs. time open arms | 1,7 | $R^2$ = 0.04 | p=0.3958 |
| S6j | Cluster entropy OA position | Vehicle<br>SKF | 9<br>9 | Two-tailed Pearson correlation | Cluster entropy Vehicle + SKF vs. time open arms | 1,7 | $R^2$ = 0.61 | ***p=0.0001 |
| S6k | SVM accuracy | Vehicle<br>SKF | 9<br>9 | 2-way ANOVA | Group main effect<br>Data type main effect<br>Group x Data type interaction | 1,32<br>1,32<br>1,32 | F=98.7<br>F=3.71<br>F=20.8 | ***p<0.0001<br>p=0.0631<br>***p<0.0001 |
| | | | | Tukey's multiple comparison test | Vehicle: Shuffle vs. Data<br>SKF: Shuffle vs. Data<br>Vehicle vs. SKF Data | 1,32<br>1,32<br>1,32 | q=14.5<br>q=5.37<br>q=6.49 | ***p<0.0001<br>***p<0.0001<br>***p=0.0004 |
